# Glycerol degradation in the thermoacidophilic crenarchaeon *Sulfolobus acidocaldarius* involves an unusual glycerol-3-phosphate dehydrogenase

**DOI:** 10.1101/2024.02.29.582781

**Authors:** Christian Schmerling, Carsten Schroeder, Xiaoxiao Zhou, Tobias Busche, Jörn Kalinowski, Lidia Montero, Oliver Schmitz, Farnusch Kaschani, Sabrina Ninck, Markus Kaiser, Jan Bost, Bianca Waßmer, Sonja V. Albers, Bettina Siebers, Christopher Bräsen

## Abstract

Glycerol is highly abundant in nature and serve as carbon source for many organisms. Also, several Archaea have the genetic capacity to grow on glycerol but its degradation has so far only been studied *Haloferax volcanii.* Herein, the thermoacidophilic crenarchaeon *Sulfolobus acidocaldarius* was shown to grow with glycerol as sole carbon and energy source. After uptake likely involving facilitated diffusion, glycerol is degraded via phosphorylation to glycerol-3-phosphate followed by oxidation to dihydroxyacetone phosphate (DHAP) catalyzed by glycerol kinase (GK) by an unusual quinone reducing FAD-dependent glycerol-3-phosphate dehydrogenase (G3PDH), respectively. The *S. acidocaldarius* genome harbors two paralogous copies of each GK and G3PDH. However, only one of these GK-G3PDH couples (Saci_2031-2033) is highly upregulated on glycerol. Deletion of the *saci_2033* gene encoding GK abolished growth on glycerol and GK activity in crude extracts. In contrast, deletion of the second GK gene (*saci_1117*) had only minor effects indicating that only one of the two GK-G3PDH couples is essential. Biochemical characterization revealed that both isoenzymes of each, GK and G3PDH, were functionally similar. Whereas the GKs showed high similarity to known enzymes from Bacteria and Eukaryotes, the G3PDHs represent unusual homologues of the bacterial GlpA subunit of the GlpABC complex with remarkable C-terminal sequence differences and a novel type of membrane anchoring via a CoxG-like protein (Saci_2031). Further sequence analyzes discovered a higher versatility of G3PDHs in Archaea with respect to interacting proteins, electron transfer, and membrane anchoring likely reflecting tailored evolutionary solutions to meet different requirements caused by life styles and electron acceptors.

## Introduction

Glycerol is an integral constituent of membrane phospholipids and storage lipids and is thus a highly abundant organic compound in nature. Accordingly, many organisms from Archaea and Bacteria to complex Eukarya are able to utilize glycerol as carbon and energy source.

Before glycerol is channeled into metabolism, it needs to be transported across the cytoplasmic membrane which often involves facilitated diffusion via aqua(glycerol)porins, transmembrane proteins catalyzing the rapid equilibration of glycerol concentration gradients across the membrane (Stroud et al., 2003). In Bacteria like *Escherichia coli* these glycerol-uptake facilitator (GUF) proteins are encoded by the *glpF* gene. However, glycerol as a small uncharged molecule can also enter the cell via passive diffusion through the cytoplasmic membrane (Richey and Lin, 1972). In few Bacteria and also some Eukarya, alternative glycerol transporters have been described (Wille et al., 1998; Ding et al., 2012; Blötz and Stülke, 2017; Klein et al., 2017; Mahdizadeh et al., 2021).

Once transported, glycerol metabolism in bacteria and eukaryotes (e.g*. E. coli*, *Pseudomonas aeruginosa,* and yeast) as depicted in Figure 1 essentially follows two biochemical routes: The first and most widespread route is utilized (mainly) by respiring organisms (Li, 1976; Klein et al., 2017; Poblete-Castro et al., 2020), which first convert intracellular glycerol into *sn*-glycerol-3-phsophate (G3P) via a glycerol kinase (GK) (encoded by the *glpK* gene) using ATP as phosphoryl donor (Yeh et al., 2004). GKs have been characterized from several Bacteria and Eukarya. They are dimers or tetramers and some, including the *E. coli* enzyme, show posttranslational regulation via FBP inhibition accompanied by dimer-tetramer transition (Applebee et al., 2011).

**Figure 1:**
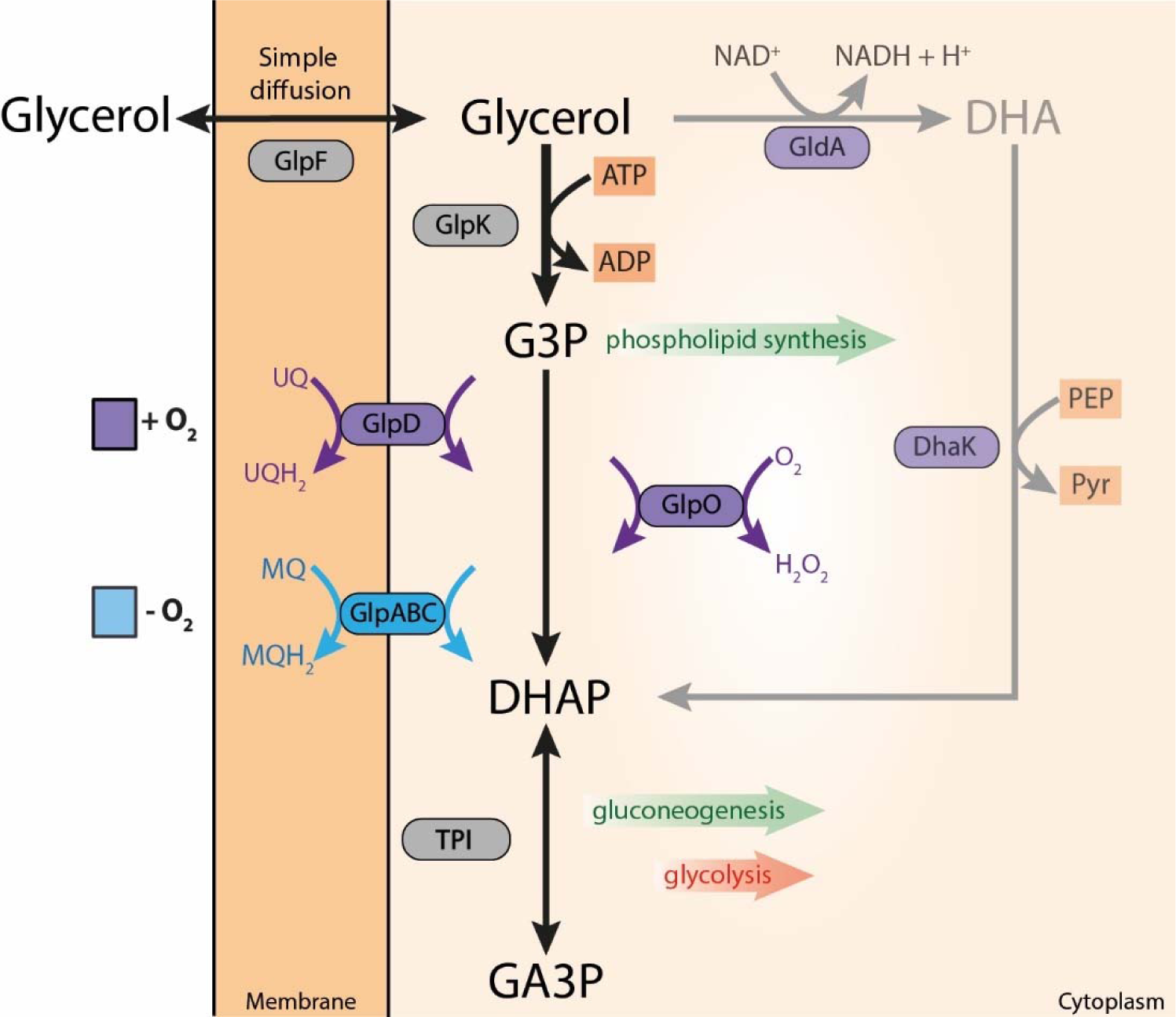
Biochemical pathways involved in glycerol conversion in Bacteria (e.g., *E. coli, Streptococcus or Pseudomonas putida*) and Eukarya forming dihydroxyacetone phosphate (DHAP) which is channeled into central metabolism. Glycerol metabolism starts with its uptake either via facilitated diffusion mediated mainly by the glycerol uptake facilitator (GlpF) (encoded by the *glpF* gene) or via (protein independent) simple diffusion through the cytoplasmic membrane. Following uptake, glycerol is converted finally to DHAP via two basic pathways: In respiring organisms, glycerol is first phosphorylated by the glycerol kinase (GlpK) (encoded by the *glpK* gene) to G3P which is further oxidized by two different membrane bound FAD dependent G3P dehydrogenases (G3PDH), i.e. GlpD (encoded by the *glpD* gene) and GlpABC (encoded by the *glpA*, *B*, and *C* genes). Electrons are transferred via the G3PDH bound FAD cofactor to ubiquinone (UQ) by GlpD or to menaquinone (MQ) (GlpABC) of the respiratory chain. A third mechanism of G3P oxidation mainly known from aerotolerant lactic acid bacteria as well as *Mycoplasma* species is catalyzed by a soluble, cytoplasmic FAD dependent G3P oxidase (GlpO, encoded by the *glpO* gene) which directly utilizes molecular oxygen as electron acceptor. The second basic glycerol converting pathway is known from anaerobic, fermentatively growing organisms. Here, glycerol is first oxidized via an NAD^+^ dependent G3PDH to dihydroxyacetone (GldA, encoded by the *gldA* gene) which is subsequently phosphorylated by dihydroxyacetone kinase (DhaK, encoded by *dhaK* gene) with phosphoenolpyruvate (less frequently ATP) as phosphoryl donor. In all cases, DHAP is further degraded via the lower common shunt of the ED and EMP pathway or utilized for gluconeogenesis. G3P serves as building block for phospholipid/membrane synthesis in Bacteria and Eukarya.

Following phosphorylation, one of two membrane-bound glycerol-3-phosphate dehydrogenases (G3PDH) hereafter designated as GlpD and GlpABC, encoded either by the *glpD* gene or by the operon comprised of the three genes *glpA, B, and C*, respectively, catalyze the oxidation of G3P to dihydroxyacetone phosphate (DHAP), both with the simultaneous reduction of a non-covalently enzyme bound flavin adenine dinucleotide (FAD) to FADH_2_. The electrons are then further transferred from the FADH_2_ to the quinone pool of the respiratory chain (Schryvers et al., 1978; Schryvers, 1981; Austin and Larson, 1991; Yeh et al., 2008). GlpD (also called the aerobic G3PDH) is maximally expressed under aerobic conditions for example in *E. coli*. (Schryvers et al., 1978; Austin and Larson, 1991; Yeh et al., 2008; Poblete-Castro et al., 2020) and transfers electrons to ubiquinone from where they are finally passed on to oxygen or nitrate. GlpD is also the G3P oxidizing enzyme in the mitochondria of Eukarya (Yeh et al., 2008). The second G3PDH (GlpABC) mainly known from Bacteria is induced under anaerobic conditions (the anaerobic G3PDH) and reduces menaquinone with the final acceptors nitrate or fumarate (in *E. coli*) (Cole et al., 1988; Poblete-Castro et al., 2020). The A and B subunits (encoded by *glpA* and *B*) form a soluble and active dimer (Schryvers and Weiner, 1981) which is likely anchored to the membrane via the C subunit (Cole et al., 1988; Varga and Weiner, 1995). A third mechanism of G3P oxidation is less widespread and mainly restricted to aerotolerant/microaerophilic lactic acid bacteria which are heme deficient and thus lack functional respiratory chains. These organisms employ the G3P oxidase (GlpO, encoded by the *glpO* gene), a cytosolic, soluble FAD-dependent enzyme directly reducing oxygen to H_2_O_2_ which is detoxified via peroxidase or catalase (Parsonage et al., 1998; Finnerty et al., 2002; Colussi et al., 2008; Zotta et al., 2017) (Figure 1). Also, the *Mycoplasma* spp. from the Mollicutes use GlpO for G3P oxidation and the resulting H_2_O_2_ has been discussed to contribute to pathogenicity of these organisms (Elkhal et al., 2015; Maenpuen et al., 2015). Finally, the DHAP produced by G3P oxidation is channeled into the central metabolism via glycolysis and gluconeogenesis, respectively. G3P can directly be used for (phospho)lipid biosynthesis in Bacteria and Eukaryotes.

The second route of glycerol processing is less abundant and restricted to organisms growing under fermentative conditions as described for some Enterobacteriaceae including *E. coli* and a few other bacterial species (Clomburg and Gonzalez, 2013; Clomburg et al., 2022). In this pathway glycerol is first oxidized to dihydroxyacetone (DHA) in an NAD^+^ dependent manner (glycerol dehydrogenase encoded by the *gldA* gene) followed by PEP (*dhaK*) or ATP (*glpK*, *Klebsiella pneumoniae*) dependent phosphorylation to DHAP (Clomburg and Gonzalez, 2013; Poblete-Castro et al., 2020) (Figure 1). The DHAP then again enters the lower shunt of the Embden-Meyerhof-Parnas (EMP) pathway finally yielding the major fermentation products of the respective organisms or is utilized for anabolism. The NADH derived from glycerol oxidation is reoxidized with another molecule of glycerol which is first dehydrated to 3-hydroxypropionaldehyde and then reduced to 1,3-propandiol, or converted to methylglyoxal via DHAP followed by reduction to 1,2-propanediol (Clomburg and Gonzalez, 2013).

Whereas the glycerol metabolism is quite well understood in Bacteria and Eukarya, comparably little is known regarding glycerol degradation in Archaea. The haloarchaeon *Haloferax volcanii* has been shown to utilize glycerol as carbon and energy source and to employ homologues of the bacterial GlpK and GlpABC proteins, while GlpD is absent (Sherwood et al., 2009; Rawls et al., 2010; Rawls et al., 2011; Williams et al., 2017). Furthermore, GK activity has been reported in crude extracts of *H. volcanii* and *Halobacterium* sp. to be induced in the presence of glycerol although basal levels have also been observed in its absence (Nishihara et al., 1999; Sherwood et al., 2009). Comparative sequence analyses revealed GK (GlpK) and G3PDH (GlpABC) homologues in various other Halobacteriales (Villanueva et al., 2017; Williams et al., 2017). However, other representatives from the archaeal domain have so far not been shown to grow with glycerol as sole source of carbon and energy although genes encoding putative GKs and G3PDHs were identified in aerobic and anaerobic representatives of the orders Thermococcales, Thermoplasmata, Thermoproteales and Lokiarchaeota (Villanueva et al., 2017). In accordance with the distribution of sequence homologues, GK activities in crude extracts from *Thermoplasma* strains as well as from *Pyrococcus* strains were shown, although the cells were grown in the absence of glycerol (Nishihara et al., 1999). For *Thermococcus kodakarensis*, induction of the G3PDH homologue in the stationary phase of growth was reported (Gagen et al., 2016). However, although both recombinant GK (also homologous to the bacterial/eukaryal GlpK) and G3PDH have been characterized from *T. kodakarensis* and also the GK crystal structure was solved (Koga et al., 1998; Koga et al., 2008; Koga et al., 2019; Hokao et al., 2020), the organism could not be grown with glycerol as carbon and energy source (Koga et al., 2019). Also, for the other archaeal organisms possessing GK and G3PDH homologues growth on glycerol remains to be shown.

The thermoacidophilic, aerobic crenarchaeal model organism *Sulfolobus acidocaldarius* grows heterotrophically at temperatures around 75°C and a pH of 3.0 with a variety of different organic carbon sources including mono-(e.g. pentoses and hexoses), di-, and polysaccharides, peptides and amino acids, as well as fatty acids. The central metabolic network particularly the carbohydrate metabolism is well understood which employs a modified unusual branched Entner-Doudoroff pathway (ED) and the Weimberg pathway for hexose and pentose degradation, respectively, as well as the EMP pathway for gluconeogenesis (Bräsen et al., 2014; Lewis et al., 2021). The organism’s genome is fully sequenced and genetic tools have been developed (for literature see (Lewis et al., 2021)). In *S. acidocaldarius* and other Sulfolobales, only GK homologues have been identified whereas G3PDH homologues appeared absent (Villanueva et al., 2017) and they have been indicated not to grow on glycerol (Grogan, 1989). Since it has recently been shown that *S. acidocaldarius* cleaves triacylglycerides by means of esterases and to grow with fatty acids as sole carbon and energy source (Wang et al., 2019), we reexamined the growth of *S. acidocaldarius* on the other lipid cleavage product glycerol. Using growth studies, polyomics approaches, crude extract enzyme measurements, biochemical and deletion mutant analyses, we herein demonstrate that *S. acidocaldarius* utilizes glycerol as growth substrate and employs a conserved “classical” GK (homologous to GlpK) for glycerol phosphorylation. However, G3P oxidation is catalyzed by a truncated GlpA-subunit like G3PDH lacking the B and C subunits of the classical, bacterial GlpABC complex. Instead, it shows an unusual type of membrane association facilitated by a small CoxG-related protein.

## Material and Methods

### Cultivation of *S. acidocaldarius* strains

Standard growth of *S. acidocaldarius* MW001 (uracil auxotroph mutant) (Wagner et al., 2012) was performed at 75°C and pH 3 in Brock’s basal salt medium (Brock et al., 1972), supplemented with 0.1 % (w/v) NZ-amine, 0.2 % (w/v) dextrin, and 10 µg/ml of uracil. For growth on glycerol, *S. acidocaldarius* MW001 were adapted for several weeks in Brock’s basal salt medium with 10 µg/ml uracil in the presence of 40 mM glycerol as sole carbon and energy source until growth was observed. For strain purification, cells were streaked on Brock’s basal salt medium plates containing 0.1 % (w/v) NZ-amine, 0.2 % (w/v) dextrin, 10 µg/ml of uracil, and 0.6 % (w/v) Gelrite (for solidification, Carl Roth, Germany) and a single colony was regrown in Brock’s basal salt medium containing 10 mM glycerol. This adapted strain denoted as *S. acidocaldarius* MW00G was stored as glycerol stock culture. To this end, *S. acidocaldarius* MW00G was grown on 10 mM glycerol to an OD_600_ of 0.6, cells were pelleted at 4,300xg and 21°C for 10 min, and resuspended in standard Brock medium containing 50 % (v/v) glycerol to a calculated OD_600_ of 10 and stored at −80°C until used.

Growth experiments with the glycerol-adapted *S. acidocaldarius* MW00G were performed in triplicates at 76°C and 120 rpm under constant shaking (Innova®44, New Brunswick, Germany) in the presence 10 mM glycerol, 20 mM glycerol or 40 mM glycerol as sole carbon and energy source. For comparison the same strain was cultivated with 0.2 % (w/v) D-xylose. Cells were inoculated into Brock’s basal salt medium containing the respective carbon source to an OD_600_ of 0.05 and growth was monitored as increase in OD_600_ over time. For growth experiments with the deletion mutant strains, cells were grown in Brock medium with 0.1 % (w/v) NZ-amine and 0.2 % (w/v) dextrin to an OD_600_ between 0.8 and 1. 50 ml Brock medium supplemented with 10 mM glycerol was then inoculated to an OD_600_ of 0.05 and growth was monitored as increase in OD_600_ over time. The glycerol concentration in the medium was determined enzymatically using the recombinantly produced Saci_2033 glycerol kinase (see below) and ADP formation was quantitatively assayed spectrophotometrically as oxidation of NADH via pyruvate kinase (PK) (Merck, Germany) and lactate dehydrogenase (LDH) (Merck, Germany) at 340 nm (42°C) in a TECAN infinite M200 microplate reader (Tecan Trading AG, Switzerland). Thus, per mole of glycerol one mole of NADH was reduced. The reaction mix contained 0.1 M MOPS-KOH buffer pH 6.5, 0.6 mM NADH, 1 mM PEP, 5 mM ATP, 5 mM MgCl_2_, 25.5 U LDH, 2.8 U PK, 0.4 U Saci_2033, and 20 µl glycerol containing sample corresponding to a maximal glycerol concentration of 0.5 mM in a total volume of 0.2 ml. For poly-omic analysis, *S. acidocaldarius* MW00G was pre-grown in Brock’s basal salt medium containing the respective glycerol concentration for two passages in 50 ml and 200 ml of culture medium, respectively. With these precultures, 400 ml main cultures (four replicates) for poly-omic analysis with the respective carbon sources were inoculated to a starting OD_600_ of 0.05 and grown to exponential phase (OD_600_ of 0.8). Cells were cooled down with liquid nitrogen and harvested at 9,000xg, 15 min, 4°C. Cell pellets were stored at −70°C and used for transcriptomics, proteomics and metabolomics.

### RNA-seq

#### RNA isolation, library preparation, and next-generation cDNA sequencing

RNA was isolated using Zymo Direct-zol RNA Miniprep kit following manufactures instructions. The RNA quality was checked by Trinean Xpose (Gentbrugge, Belgium) and the Agilent RNA Nano 6000 kit using an Agilent 2100 Bioanalyzer (Agilent Technologies, Böblingen, Germany). Pan-Archaea riboPOOL kit from siTOOLs Biotech was used to remove the rRNA. TruSeq Stranded mRNA Library Prep Kit from Illumina was applied to prepare the cDNA libraries. The cDNAs were sequenced paired end on an Illumina NextSeq 500 system (San Diego, CA, USA) using 74 bp read length mid output.

#### Bioinformatics data analysis, read mapping and analysis of differential gene expression

The paired-end cDNA reads were mapped to the *Sulfolobus acidocaldarius* DSM 639/MW001 genome sequence (accession number CP000077.1) using bowtie2 v2.2.7. with default settings for paired-end read mapping. All mapped sequence data were converted from SAM to BAM format with SAMtools v1.3 (Li et al., 2009) and imported to the software ReadXplorer v.2.2 (Hilker et al., 2016).

Differential gene expression analysis of four replicates including normalization was performed using Bioconductor package DESeq2 (Love et al., 2014) included in the ReadXplorer v2.2 software (Hilker et al., 2016). The signal intensity value (A-value) was calculated by the log2 mean of normalized read counts and the signal intensity ratio (M-value) by the log2 fold change. The evaluation of the differential RNA-seq data was performed using an adjusted P-value cutoff of P ≤ 0.01 and a signal intensity ratio (M-value) cutoff of ≥2 or ≤−2. Genes with an M-value/log2 fold change outside this range and adjusted P ≤ 0.01 were considered as differentially up-or downregulated.

### Proteome

Samples for proteomic analysis were prepared using the single-pot, solid-phase-enhanced sample-preparation (SP3) strategy (Hughes et al., 2019). All buffers and solutions were prepared with mass spectrometry (MS)-grade water (Avantor, Radnor, PA, USA). Cell pellets were taken up in 200 µl 1x sample buffer (50 mM HEPES pH 8.0, 1 % (w/v) SDS, 1 % (v/v) NP-40, 10 mM TCEP, 40 mM chloroacetamide) and the samples were heated for 5 min at 95°C prior to sonication with a Bioruptor UCD-200 (Diagenode, Seraing, Belgium) device for ten cycles of 1 min pulse and 30 sec pause at high power. The protein extracts were centrifuged (20,000xg, RT, 20 min) and the protein concentration of the cleared lysates was determined using the Pierce 660 nm Protein Assay Reagent (#22660; Thermo Scientific, Waltham, MA, USA) with the Ionic Detergent Compatibility Reagent (#22663; Thermo Scientific, Waltham, MA, USA) according to the manufacturers’ instructions. Next, 15 µg of total protein in a volume of 47 µl 1x sample buffer was treated with 7 U of benzonase (#70746; Merck Millipore, Burlington, MA, USA) in dilution buffer (20 mM HEPES pH 8.0, 2 mM MgCl_2_; 37 °C, 30 min 1500 rpm), followed by the addition of iodoacetamide to a final concentration of 10 mM to complete alkylation of cysteine residues (RT, 30 min, 1500 rpm, in the dark), resulting a sample volume of 50 µl. Then, 3 µl of a 50 µg µl^-1^ 1:1 mixture of hydrophilic (#45152105050250) and hydrophobic (#65152105050250) carboxylate modified Sera-Mag™ SpeedBeads (Cytiva, Marlborough, MA, USA) that were washed twice with MS-grade water were added to the samples. Protein binding was induced by the addition of an equal sample volume of pure ethanol (24 °C, 20 min, 1500 rpm), the beads were collected using a magnetic stand. Beads were allowed to bind for at least 5 min before the supernatant was removed. The beads were washed thrice with 180 µl 80 % (v/v) ethanol prior to the addition of the digestion enzyme mix (0.6 μg of trypsin (V5111; Promega, Madison, WI, USA) and 0.6 µg LysC (125-05061; FUJIFILM Wako Pure Chemical, Osaka, Japan) in 25 mM ammonium bicarbonate) and incubation of the samples at 37 °C (16 h, 1500 rpm). Next day, the samples were briefly centrifuged (10 sec, 500 rpm) and placed on a magnet for 5 min. The clear solution containing the tryptic peptides was transferred to a new Eppendorf tube. The beads were taken up in 47 µl 25 mM (RT, 10 min, 1500 rpm). After incubation, the tubes were placed once more on a magnetic stand and after 5 min the clear supernatant was again collected and combined with the recovered peptide mix, followed by the addition of trifluoroacetic acid (TFA; 2 % (w/v) final concentration) to the samples. Prior to LC-MS/MS analysis, peptides were desalted as described in (Klaus et al., 2022) with the only modification of not re-applying the flow-through to the C_18_StageTips again.

### LC-MS/MS Analysis

LC-MS/MS analysis of peptide samples was performed on an Orbitrap Fusion Lumos mass spectrometer (Thermo Scientific, Waltham, MA, USA) coupled to an EASY-nLC 1200 liquid chromatography (LC) system (Thermo Scientific, Waltham, MA, USA) that was operated in the one-column mode. The samples were separated on a self-packed analytical column (see supplementary File 1) filled with Reprosil-Pur 120 C18-AQ 1.9 μm (Dr. Maisch, Ammerbuch-Entringen, Germany) that was encased by a PRSO-V2 column oven (Sonation, Biberach an der Riß, Germany) and attached to a nanospray flex ion source (Thermo Scientific, Waltham, MA, USA). During data acquisition, the column oven temperature was adjusted to 50°C. The LC was equipped with solvent A (0.1 % (w/v) FA, in water) and solvent B (0.1 % FA (w/v), in 80 % acetonitrile (ACN)) prepared from UHPLC (ultra-high-performance liquid chromatography)-grade solvents (Honeywell, Charlotte, NC, USA) as mobile phases. Peptides were directly loaded onto the analytical column with a maximum flow rate so that the set pressure limit of 980 bar would not be exceed (usually around 0.5 – 0.8 μl min^−1^) and separated by running gradients with different length and composition (for details see supplementary File 1).

The mass spectrometer was operated in the positive ion mode using Xcalibur software (v4.3.7.3.11). Precursor ions (MS^1^) were scanned in the Orbitrap analyzer (FTMS; Fourier Transform Mass Spectrometry) with the internal lock mass option switched on (lock mass was 445.120025 m/z, polysiloxane (Olsen et al., 2005)). Data dependent product ion spectra (MS^2^) were recorded in the ion trap. All relevant MS settings (resolution, scan range, AGC, ion acquisition time, charge states, isolation window, fragmentation type and details, cycle time, number of scans performed, and various other settings) can be found in supplementary File 1.

### Peptide and Protein Identification Using MaxQuant and Perseus

Recorded RAW data was analyzed with MaxQuant (v. 1.6.17.0 or v.2.0.1.0) using the default settings (Cox and Mann, 2008) with the Label-free quantification (LFQ) (Cox et al., 2014) and match between runs options activated. MS/MS spectra were searched against the UniProt *Sulfolobus acidocaldarius* (DSM639) (UP000001018_330779.fasta; 2222 entries; downloaded on 2020-08-02) database. A search against a contaminants database as implemented in MaxQuant (contains known MS contaminants; 246 sequences) was included to estimate the level of contamination.

For further data analysis and filtering of the MaxQuant output, LFQ intensities were loaded from the proteinGroups.txt file into Perseus (v. 1.6.14.0 or v.1.6.15.0) (Tyanova et al., 2016). Contaminants as well as hits only identified based on peptides with a modification site and hits from the reverse database were removed. To allow comparison of the different treatment groups, biological replicates were combined into categorical groups and the data was transformed to the log_2_(x) scale. For the full proteome analysis, only protein groups (PGs) with a valid LFQ intensity in at least three out of four replicates in a minimum of one categorical group were kept for further analysis. For the identification of interaction partners of the CoxG homologs Saci_2031 and Saci_1119, the data was separately filtered to only keep hits with a valid LFQ intensity in at least two out of three replicates for samples containing the respective HA-tagged protein. The log_2_-fold change in normalized protein group quantities between the different categorical groups was determined based on the mean LFQ intensities of replicate samples (relative quantification). To enable quantification, missing LFQ intensities were imputed from a normal distribution (full proteome analysis: width 0.3, down shift 1.8; identification of interaction partners: width 0.3, down shift 2.0). The statistical significance of the difference in LFQ intensity was determined via a two-sided Student’s t-test. Full MS data for the comparative full proteome analysis and the identification of interaction partners of Saci_2031 and Saci_1119 can be found in supplementary Files 2 and 3 Genes that were up-or downregulated with a log2-fold change ≥ 2 in response to growth on glycerol and are shared between the transcriptomics and proteomics analyses are reported in supplementary Table 1. Proteins enriched by co-IP with a log2-fold change ≥ 2, excluding ribosomal proteins, are reported in supplementary Table 2.

### Metabolome analyses

#### Metabolite extraction

The protocol for the metabolite extraction was based on the method described by *Sellick et al*. with slight modifications (Sellick et al., 2011). It consisted of cell lysis disruption by resuspending the cell pellet in 500 µl of prechilled (−80 °C) methanol and the addition of 20 µl of internal standards (fructose 6-phosphate, arginine 13C6, and succinic acid d4). The mixture was vortexed for 2 min and sonicated for 2 min. After that, the sample was freeze at −80 °C for 5 min and the vortex and sonication steps were repeated. The methanol was evaporated by vacuum concentration (Concentrator plus, Eppendorf, Hamburg, Germany). When the extract was completely dry, 250 µl of water were added and the process was repeated. During the whole process the cells were kept on ice. Then, after the drying the water, the extract was resuspended in 100 µl ACN/water (85:15, v/v), sonicated 2 min, and vortexed 2 min. The mixture was centrifugated at 12000 rpm for 2 min (MiniSpin centrifuge, Eppendorf, Hamburg, Germany). The supernatant was transferred to a LC vial.

#### LC-ESI-QTOF method for the metabolomics analysis

The samples were analyzed by a 1290 Infinity II LC instrument coupled to a 6546 LC/Q-TOF high-resolution mass spectrometer, and the ionization was performed using a Dual Jet Stream source in negative mode (Agilent Technologies, Waldbronn, Germany). For the LC separation, a iHILIC-(P)Classic (150 x 2.1 mm, 5 µm) (Hilicon, Umeå, Sweden) was used. The mobile phases consisted of 5 mM Ammonium acetate, pH 5 (A) and ACN/5 mM ammonium acetate, 85:15 (*v/v*), pH5 (B). The gradient elution was stablished as follows: 0 min, 90% B; 22 min, 80% B, 30 min, 65% B; 35 min, 65% B. The flow rate was set at 0.2 mL/min and the column temperature was kept at 40 °C. The electrospray ion source parameters were: gas temperature, 200 °C; dry gas, 10 ml/min; nebulizer, 40 psi; sheath gas temperature, 300 °C; sheath gas flow, 12 L/min; fragmentor, 125 V; skimmer, 65 V; capillary voltage, 3000 V. Full scan-data dependent acquisition was used to perform the tandem MS experiments.

### Crude extract enzyme measurements

For preparation of crude extracts, *S. acidocaldarius* MW00G was grown in 50 ml cultures on 40 mM glycerol and - for comparison - 0.2 % (w/v) D-xylose to an OD_600_ of 0.6. For crude extract measurements in deletion mutants, the cells were grown in 50 ml culture on either 10 mM glycerol or 0.2 % (w/v) NZ-amine to an OD_600_ between 0.8 and 1. Cells were collected by centrifugation (4,300xg 15 min, 21 °C), and resuspended in 5 ml of 50 mM MES-KOH pH 6.5, 20 mM KCl. Cells were lysed by sonication (3 x 8 min, 60 %, 0.5 s^-1^; on ice, UP200S, Hielscher Ultrasonics) and centrifuged again at 4,300xg, 30 min, 4 °C to remove cell debris. To separate the soluble from the membrane fraction, the supernatant was then centrifuged at 150,000xg, 60 min at 4 °C. After centrifugation, the supernatant represented the soluble fraction. For preparation of membrane fractions the pellet was washed by resuspension in 5 ml of 50 mM MES-KOH, 20 mM KCl, pH 6.5, centrifuged (150,000xg, 60 min at 4 °C), resuspended in 500 µl of 50 mM MES-KOH pH 6.5, 20 mM KCl and finally sonicated with a low amplitude (20 %, 0.5 s^-1^) until membranes were fully suspended. The so prepared membrane fraction was then used for enzyme measurements. Protein concentrations in both, soluble and membrane fractions were determined by a modified Bradford assay (Zor and Selinger, 1996) using bovine serum albumin (Carl Roth, Germany) as standard. Enzyme activities in soluble and membrane fraction were assayed spectrophotometrically using a Specord UV/visible-light (Vis) spectrometer (Analytic Jena, Germany) with protein amounts of 50 µg (crude extract) and 40 µg (membrane fraction) in a total volume of 500 µl. Assay mixtures were preincubated at the respective temperatures before starting the reaction usually with substrate (glycerol, G3P). G3PDH activity was determined as glycerol-3-phosphate (G3P) (Merck, Germany) dependent reduction of DCIPIP at 600 nm (extinction coefficient 20.7 mM^-1^ cm^-1^) and 70 °C in 50 mM MES-KOH pH 6.5, 20 mM KCl containing 50 µM DCPIP and 200 µM G3P. GK activity was assayed in 100 mM TRIS-HCl pH 7, 1 mM MgCl_2_, 1 mM ATP, 5 mM glycerol, 0.2 mM NADH, and 2 mM PEP by coupling the glycerol and ATP dependent formation of ADP to NADH oxidation via LDH (63.7 U) and PK (7 U) at 50 °C and 340 nm (extinction coefficient 6.22 mM^-1^ cm^-1^). It was ensured that the auxiliary enzymes were not rate limiting. One unit (1 U) of enzyme activity is defined as 1 µmole substrate consumed or product formed per minute.

### Molecular cloning

*E. coli* DH5α used for plasmid construction and propagation was cultivated at 37 °C and 180 rpm (INFORS shaker, Switzerland) in Lysogeny broth (LB) containing appropriate antibiotics (150 μg/ml ampicillin, 50 μg/ml kanamycin or 25 μg/ml chloramphenicol). For heterologous overexpression in *E. coli* either the pET15b (*saci_2032*) or the pETDuet-1 vector (co-expresssion of *saci_2032/saci_2031* or *saci_1118/1119,* as well as expression of *saci_2032* or *saci_1118* alone) were used and for homologous overexpression in *S. acidocaldarius* MW001 the pSVAmalFX-SH10 vector (*saci_1117*) and pBS-ara-albaUTR-FX-vector (*saci_2033*) (manuscript in preparation) were used. Plasmids and strains are listed in supplementary Table 3. Open reading frames (ORFs) *saci_1117, saci_1118, saci_1119 saci_2031, saci_2032* and *saci_2033* were amplified from genomic DNA of *S. acidocaldarius* DSM 639 wild type using the primers (Eurofins Genomics) listed in supplementary Table 4 employing the Q5 polymerase (NEB, USA) following the manufacturers’ instructions. After restriction digest with the respective restriction endonucleases (NEB, USA) (supplementary Table 4), the purified PCR products were ligated with the empty vectors in a molar ratio of 1:4 using T4 DNA ligase (NEB, USA) at 16 °C overnight. For construction of pETDuet-1 co-expression vectors the gene pairs *saci_2032* and *saci_2031* or *saci_1118* and *saci_1119* were sequentially ligated into multiple cloning site 1 (*saci_2032/saci_1118*) and 2 (*saci_2031/saci_1119*). For homologous overexpression of *saci_2031* and *saci_1119* in course of the co-immunoprecipitation experiments (see below), the PCR-amplified gene was cloned into pSVA-araFX-HA vector (Tsai et al., 2020) using the *Nco*I/*Xho*I restriction sites. Plasmid constructs (pET15b-*saci_2032*; pET15b-*saci_1118*; pETDuet-1-*saci_2032*; pETDuet-1-*saci_1118,* pETDuet-1-*saci_2031/saci_2032*, pETDuet-1-*saci_1118/saci_1119,* pSVAmalFX-SH10-*saci_1117,* pBS-ara-albaUTR-FX-*saci_2033,* and pSVA13204) were propagated in *E coli* DH5α and successful cloning was confirmed by commercially available sequencing services (Eurofins genomics).

### Construction of markerless single and double glycerol kinase deletion mutants Δ*saci_1117*, Δ*saci_2033* and Δ*saci_1117 Δsaci_2033*

To obtain the markerless deletions of *saci_1117* and *saci_2033*, the plasmids pSVA12818 and pSVA12822 were constructed. Briefly, 500bp of the upstream and downstream region of each gene were amplified by PCR and the respective PCR products were annealed via overlap extension PCR. The resulting products were then cloned into pSVA407 or pSVA431 (for plasmids and strains, see supplementary Table 3). The resulting plasmids were then methylated by transformation in *E. coli* ER1821 to prevent plasmid degradation in the recipient strain. Methylated plasmid was then used to transform MW00G as described previously for MW001 (Wagner et al., 2012). To prepare competent MW00G, a pre-culture was grown in Brock media supplemented with 0.1 % (w/v) NZ-amine and 0.2 % (w/v) dextrin and 20 µg/mL uracil to an OD_600_ of 0.5-0.7 and used to inoculate 50 mL Brock media supplemented with 0.1 % (w/v) NZ-amine and 10mM glycerol. The culture was harvested at an OD_600_ of 0.2 - 0.3 and further prepared as described previously (Wagner et al., 2012).

### Complementation studies

For the complementation of MW1257 *Δ1117* and MW1258 Δ*2033*, the respective genes *saci_1117* and the operon *saci_2033--2034* were amplified with 200 bp of the native promoter region together with 500 bp of the *upsE* (*saci_1494*) upstream and downstream region using the primers listed in the supplementary Table 4. The resulting PCR products were then assembled in an *ApaI* and *SacII* digested pSVA407 vector via Gibson Assembly following the manufacturer’s instructions (New England Biolabs). The respective deletion mutants were then transformed with the resulting knock-in plasmids pSVA407*-ΔupsE:P_1117_ saci_1117* CtSS and pSVA407-*ΔupsE:P_2033_ saci_2033-2034.* Correct integration was confirmed by colony PCR. Complementation was then evaluated by growth comparison in Brock medium supplemented with 10 mM glycerol as described above.

### Homologous overexpression and affinity purification of glycerol kinase

For homologous expression, the pSVAmalFX-SH10-*saci_1117* and the pBS-Ara-albaUTR-FX-*saci_2033*-CtSS vector were transformed in *S. acidocaldarius* MW001. To prevent degradation in the recipient strain, the plasmids were methylated by transformation into *E. coli* ER1821. The methylated plasmid was transformed into electro-competent wild type *S. acidocaldarius* MW001 cells via electroporation using a Gene Pulser Xcell (BioRad, München, Germany) with a constant time protocol with input parameters 2 kV, 25 μF, 600 Ω in 0.5 mm cuvettes. Cells were regenerated for 45 min at 75°C in recovery solution (1 % (w/v) sucrose, 20 mM β-alanine, 1.5 mM malate buffer, pH 4.5, 10mM MgSO_4_) (Wagner et al., 2012), and then plated on the same medium solidified by Gelrite (see above) without uracil. Successful transformation was confirmed via colony-PCR. Clones harboring the desired plasmid construct with the respective gene insert were inoculated in a 50 ml preculture of liquid Brock’s basal salt medium with 0.1% (w/v) NZ-amine, 0.2% (w/v) dextrin. The latter was used then to inoculate a 2 L liquid Brock’s basal salt medium (OD_600_ 0.05) containing 0.1 % (w/v) NZ-amine and 0.2% (w/v) D-xylose for induction and cells were grown for two days at 76°C under constant shaking at 120 rpm. Afterwards, cells were harvested by centrifugation at 7,000 x g and 4°C for 15 minutes and stored at −80°C for further use. For protein purification cells were resuspended in 50 mM TRIS-HCl pH 8.0, containing 250 mM NaCl in a ratio of 3 ml buffer to 1 g of cells (wet weight) and disrupted by sonication (3 x 8 min, 60% amplitude, 0.5 s^-1^; UP200S, Hielscher Ultrasonics). Cell debris was removed by centrifugation at 30,000xg and 4°C for 45 min and the resulting crude extract was applied onto a Strep-Tactin®XT 4Flow® column (IBA Lifesciences, Germany) and purified following the manufacturers’ instructions. Protein purity was analyzed by SDS-PAGE and protein concentration was determined by a modified Bradford assay (Zor and Selinger, 1996) using bovine serum albumin (Carl Roth, Germany) as standard.

### Heterologous overexpression and affinity purification of G3PDHs

For expression the pET15b-vector-*saci_2032* was transformed into *E. coli* Rosetta (DE3) [pLys] and grown in terrific broth (TB) (Carl Roth, Germany) supplemented with 150 µg/ml ampicillin and 25 µg/ml chloramphenicol. After growth at 37 °C to an OD_600_ of 0.8 cells were induced with 1 mM IPTG and further grown at 20 °C for 17 h. Cells were harvested by centrifugation (6,000 x g, 20 min, and 4 °C) and stored immediately at −80 °C for further use. Chromatographic purification was carried out using an ÄktaPurifier system (Cytiva, USA). After thawing, cells were resuspended in 10 ml buffer per g cells (wet weight) containing 50 mM TRIS-HCl pH 7.8, 10 mM imidazole, 150 mM NaCl, supplemented with 1 mM FAD and disrupted by sonication (50 % amplitude, 0.5 s^-1^; UP200S, Hielscher Ultrasonics). Afterwards cell debris was removed by centrifugation (16,000xg for 45 min, 4 °C), the supernatant was filtered (0.45 polyvinylidene fluoride membrane, Carl Roth, Germany) and applied onto a 1 ml Nickel-IDA column (GE Healthcare) equilibrated in 50 mM TRIS-HCl pH 7.8, 10 mM imidazole, 150 mM NaCl and proteins were eluted with a linear gradient from 10-300 mM imidazole in the same buffer. G3PDH containing fractions as judged by SDS-PAGE and activity measurements were pooled and imidazole was removed using a centrifugal concentrator (10 kDa cutoff, Sartorius, Germany). Protein aliquots were frozen in liquid nitrogen and stored at −80 °C for further use. Protein purity was analyzed by SDS-PAGE and protein concentration was determined by a modified Bradford assay (Zor and Selinger, 1996) using bovine serum albumin (Carl Roth, Germany) as standard. To determine the FAD concentration of the purified G3PDH, the protein was denatured with 0.5 % (w/v) SDS and FAD absorbance spectra were determined from 300 nm to 600 nm. The FAD content was calculated using an extinction coefficient of 11.300 mM^-1^ cm^-1^ at 450 nm.

### Determination of the native molecular mass of purified proteins

Size exclusion chromatography was used to determine the native molecular mass of all purified enzymes. To this end, pooled enzyme samples after affinity chromatography were concentrated using centrifugal concentrators (10 kDa cutoff) (Sartorius, Germany) and applied to size exclusion chromatography (Superdex 200 HiLoad 26/60 prep grade column or Superose 6/100 column (Cytavia, USA)) using 50 mM TRIS-HCl pH 7.5, containing 250 mM NaCl as buffer. For calibration, Gel Filtration LMW Calibration Kit and Gel Filtration HMW Calibration Kit (Cytavia, USA) were used.

### Localization of G3PDH

To investigate the role of CoxG (Saci_2031 and Saci_1119) for membrane anchoring of the Saci_2032/Saci_1118 G3PDHs in *S. acidocaldarius*, *saci_2032* or *saci_1118* were heterologously coexpressed with *saci_2031* or *saci_1119* from the respective pETDuet-1 expression vector construct in *E. coli* Rosetta (DE3) as described above. As a control, *saci_2032* or *saci_1118* were expressed alone from the same vector without *saci_2031* or *saci_1119*. After expression, membrane and soluble fractions were separated. To this end cells were disrupted as described above and insoluble cell debris and non-lysed cells were removed by centrifugation at 6,000xg, 4 °C for 15 minutes. Afterwards, the resulting supernatant was centrifuged at 150,000xg at 4 °C for 120 min, and the resulting supernatant (=soluble fraction) was separated from the pelleted cell membranes. The insoluble membrane fraction was washed twice with 50 mM TRIS-HCl, pH 7.0 and centrifuged (150,000xg, 4 °C, 120 min). Finally, the membrane fraction was aliquoted, flash frozen in liquid nitrogen, and stored at −80 °C for further use. For activity measurements and immunodetection, isolated membranes were thawed, mixed with 500 µl of buffer (50 mM TRIS-HCl pH 7.0) and sonicated with a low amplitude (20 %, 0.5 s^-1^) until membranes were fully resuspended. For immunodetection, 50 µg of membrane or soluble protein preparations were separated by SDS-PAGE (12.5 % (w/v) polyacrylamide gels) and transferred to a PVDF membrane using the Trans-Blot Turbo (Biorad, USA) at 25 V and1.0 A for 30 min in 25 mM TRIS, 192 mM glycine, 20 % (v/v) Ethanol, 0.075 % SDS. Membranes were blocked with 5 % (w/v) BSA in TBST (20 mM TRIS, 500 mM NaCl, 0.05 % Tween20) and then incubated with horseradish peroxidase (HRP) conjugated anti-His antibody (1:10,000 dilution; Abcam, United Kingdom) in TBST with 5 % (w/v) BSA for 90 minutes. Afterwards membranes were washed three time with 15 ml TBST for 5 min at 21 °C, and finally with 15 ml of TBS (20 mM TRIS-HCL pH7.4, 500 mM NaCl) for 5 min. Detection of His-tagged proteins was performed with the ClarityTM Western ECL substrate (Biorad, USA) and using a Versadoc (Biorad, USA) for luminescence detection.

### Co-immunoprecipitation (co-IP) Assays with anti-HA-Magnetic Beads

For each pulldown experiment, *saci_2031* or *saci_1119* were homologously expressed from the pSVA13204 vector construct transformed into *S. acidocaldarius* MW00G to yield a C-terminally HA-tagged recombinant protein. For this experiment, 500 ml cell cultures were grown in Brock’s basal salt medium with 10 mM glycerol as carbon source for 72 h to an OD_600_ 0.6 - 0.7 and expression of *saci_2031*/*saci_1119* was induced by the addition of 0.2 % (w/v) L-arabinose. After 4 h of further growth, cells were cooled down on ice and harvested by centrifugation for 20 min at 4 °C and 5,000 xg (OD_600_ 0.8 - 0.9). The pellet was then resuspended to a calculated OD_600_ of 40 in 25 mM TRIS-HCl pH 7, 150 mM KCl, 5 % (v/v) glycerol, 10 mM EDTA, and 2 % (w/v) n-Dodecyl β-maltoside (DDM) and incubated for 1 h at 37 °C while rotating. The lysed cells were then centrifuged for 30 min at 15,000xg to remove cell debris at room temperature and the supernatant was used for Co-immunoprecipitation. Therefore, 35 µl of Pierce™ anti-HA Magnetic Beads (Pierce HA-tag IP/co-IP kit (Pierce)) were added to 10 ml of the cell lysate and incubated for 1 h at room temperature while rotating. The beads were collected on a magnetic stand and washed twice with 25 mM TRIS-HCl pH 7 containing 150 mM KCl, 5 % (v/v) glycerol, 10 mM EDTA and finally once with LC-MS grade water (Merck, Germany). After washing, 35 µl of LC-MS grade water was added and 10 µl of the bead solution was taken for SDS-PAGE analysis, Western blotting and immunodetection of HA-tagged proteins with anti-HA and horse radish peroxidase-coupled secondary antibodies using an iBright™ FL1500 (ThermoFisher). The remaining 25 µl bead suspension was used as input for on-bead digestion (Klaus et al., 2022).

### Characterization of purified enzymes

Enzyme assays were performed spectrophotometrically using a Specord UV/visible-light (Vis) spectrometer (Analytic Jena, Germany) in cuvettes filled with 500 µl assay mixture (unless stated otherwise). Assay mixtures were prewarmed to the respective assay temperature and the reaction was usually started by addition of substrate. Experimental data were fitted, and kinetic constants were determined using the OriginPro 2022 software package.

#### Glycerol kinase

GK activity was determined as described above for the crude extract measurements with following modifications: For Saci_1117, the assays were performed in 0.1 M TRIS-HCl buffer (pH 7 at 50°C), 1 mM MgCl_2_, 1 mM ATP, 0.2 mM NADH, 2 mM PEP, 63.7 U LDH, 7 U PK using 0.1 - 1 µg of purified Saci_1117. For Saci_2033 the assay was performed in 0.1 M MOPS-KOH buffer (pH 6.5 at 50°C), 5 mM MgCl_2_, 5 mM ATP, 0.2 mM NADH, 2 mM PEP, 63.7 U LDH, 7 U PK using 0.1-0.5 µg of purified Saci_2033. For *K_M_* determination varying concentrations of glycerol (0 - 2 mM), DHA (0 - 30 mM), DL-glyceraldehyde (0 - 20 mM) or ATP (0-5 mM), were used at a constant concentration of 1 mM (Saci_1117) and 5 mM (Saci_2033) ATP, 1 mM (Saci_1117) and 5 mM (Saci_2033) MgCl_2_ or 2 mM glycerol, respectively. The pH optimum was determined using the same assay in a mixed buffer system ranging from pH 5.0 to pH 8.0 containing 50 mM MES, 50 mM HEPES and 50 mM TRIS-HCl. The temperature optimum was determined between 60°C to 80°C by coupling the GK mediated G3P formation to DCPIP reduction followed at 600 nm via purified, recombinant G3PDH (0.25 U) from *S. acidocaldarius* (Saci_2032) in an assay containing 0.1 M TRIS-HCl buffer (pH 7, temperature adjusted), 1 mM ATP, 1 mM MgCl_2_, 0.1 mM DCPIP and 2 mM glycerol for Saci_1117 and 0.1 M MOPS-KOH buffer (pH 6.5, temperature adjusted), 5 mM ATP, 5 mM MgCl_2_, 0.1 mM DCPIP and 2 mM glycerol for Saci_2033.To study the thermal stability, the Saci_1117 and Saci_2033 GKs were incubated at 70°C, 80°C, and 90°C in 100 mM TRIS-HCl, pH 7 or MOPS-KOH, pH 6.5 (temperature adjusted), respectively, at a protein concentration of 0.05 mg ml^-1^. After regular time intervals samples were taken and the residual activity of the enzyme was determined in the coupled assay via PK and LDH at 50°C as described above. The effect of fructose-1,6-bisphosphate (F1,6BP) on GK activity was also tested in the PK-LDH coupled assay (see above) in the presence of F1,6BP concentrations up to 1 mM as indicated. Nucleotide specificity was analyzed in the G3PDH coupled assay system using 5 mM GTP, CTP or phosphoenolpyruvate (PEP) instead of ATP at 75°C.

#### Glycerol-3-phosphate dehydrogenase

G3PDH activity was assayed spectrophotometrically at 70 °C by following the G3P dependent reduction of either DCPIP, FAD or ubiquinone-Q1 as electron acceptors. Assays were performed in 50 mM HEPES-KOH pH 6.5, 100 mM KCl. For determination of the reductive half reaction, 70 µg of Saci_2032/Saci_1118 corresponding to a final concentration of 5 µM was incubated with 50 µM of G3P. 5 min after addition of G3P, absorption spectra were measured between 400 and 500 nm in a 96 well plates (BRANDplates®, BRAND, Germany) in a Tecan infinite M200 plate reader (Tecan Trading AG, Switzerland). Loss of absorption at 450 nm indicates the reduction of bound FAD. To test the quinone reactivity, 0.6 µg of Saci_2032 or 1.2 µg of Saci_1118 were preincubated with 30 µM of ubiquinone-Q1. Afterwards 100 µM of G3P was added to the reaction mix and reduction of ubiquinone-Q1 was followed in 15 second intervals by recording the absorption spectra between 260 and 330 nm in the Tecan infinite M200 plate reader. Loss of absorption at 280 nm indicates the reduction of the ubiquinone-Q1. Continuous enzyme assays were performed by following the change (decrease) in absorbance upon reduction of the artificial electron acceptor DCPIP (600 nm) or of the ubiquinone-Q1 (280 nm). Kinetic parameters for G3P were determined in a continuous enzyme assay with 0.06 mM of DCPIP as electron acceptor and varying concentrations of G3P (0-0.3 mM). For determination of the kinetic constants for ubiquinone-Q1 concentrations were varied between 0 mM and 60 mM of the quinone at a fixed G3P concentration of 0.4 mM. Both, the pH and the temperature optimum were determined using 0.4 mM G3P and 0.06 mM DCPIP. For the pH optimum a mixed buffer system containing 50 mM MES, HEPES, and TRIS was used in the range of pH 5.0 to pH 8.0. The same assay system in 50 mM MES-KOH pH 6.5 adjusted at the respective temperature was applied for determination of the temperature optimum between 50 °C and 80 °C. To test the recombinant Saci_2032 and Saci_1118 for glycerol oxidase activity, the G3P dependent hydrogen peroxide formation was coupled to the oxidation of 2,2’-azinobis-(3-ethylbenzothiazoline-6-sulfonate) (ABTS) via horseradish peroxidase (HPR) according e.g. to (Maenpuen et al., 2015). The assay mixture contained 100 mM MES-KOH buffer at pH 6.5, 1 mM ABTS, 0.45 μg of protein and 0.2 U of HRP. The reaction was started by addition of 0.2 mM G3P and the hydrogen peroxide dependent ABTS oxidation was photometrically determined at 420 nm (extinction coefficient 42.3 mM^-1^cm^-1^). The thermostability of the Saci_2032 and Saci_1118 G3PDHs was analyzed by incubating the enzyme at 70 °C, 80 °C, and 90 °C in 50 mM MES, pH 6.5 (temperature adjusted) at a protein concentration of 0.18 mg ml^-1^. After regular time intervals the residual activity of the enzyme was determined with G3P and DCPIP as described above. The influence of detergents and membrane lipids on Saci_2032 activity was also analysed with DCPIP at 50 °C in the presence of either 0.5 % (w/v) DDM, 0.5 % (v/v) of Triton X-100, 50 µg of Phosphatidylcholine (Merck, Germany) or 50 µg of isolated *S. acidocaldarius* MW00G membrane fractions.

### Data availability

The mass spectrometry proteomics data for the on-bead digestions have been deposited to the ProteomeXchange Consortium via the PRIDE (Vizcaíno et al., 2016) partner repository (https://www.ebi.ac.uk/pride/archive/) with the dataset identifier PXD050086. During the review process the data can be accessed via a reviewer account (**Username:****; Password:** kqMtx1el)

## Results

### *S. acidocaldarius* glycerol catabolism involves GK and G3PDH

To analyze growth and glycerol consumption by *S. acidocaldarius* MW001, cells were incubated in standard Brock medium containing glycerol (10 mM, 20 mM, and 40 mM) as sole carbon and energy source (Figure 2a, b, and c, respectively). After a prolonged time of adaption (several weeks), *S. acidocaldarius* MW001 grew exponentially with similar growth rates of 0.0294 h^-1^ (10 mM glycerol), 0.0280 h^-1^ (20 mM glycerol), and 0.0284 h^-1^ (40 mM glycerol) up to cell densities of OD_600_ 4 (at 40 mM glycerol) and glycerol was completely consumed (Figure 2a-c). The adapted strain was referred to as *S. acidocaldarius* MW00G. The final OD_600_ of 4 at 40 mM glycerol corresponds to a cell dry weight (CDW) of approximately 1.33 g/l and hence a molar yield coefficient of 33.3 g CDW/mol of glycerol. Lowering the glycerol concentrations results in enhanced growth yields of 41 g CDW/mol (20 mM glycerol) and 50 g CDW/mol glycerol (10 mM glycerol) (Figure 2a and b). For comparison, *S. acidocaldarius* was grown with 0.2 % (w/v) D-xylose as sole carbon and energy source leading to a final OD_600_ of 0.8 with slower growth rates of 0.0195 h^-1^ (Figure 2d). This corresponded to a growth yield of 0.27 g/l CDW and thus a molar yield coefficient of 20 g CDW/mol D-xylose (Figure 2d).

**Figure 2:**
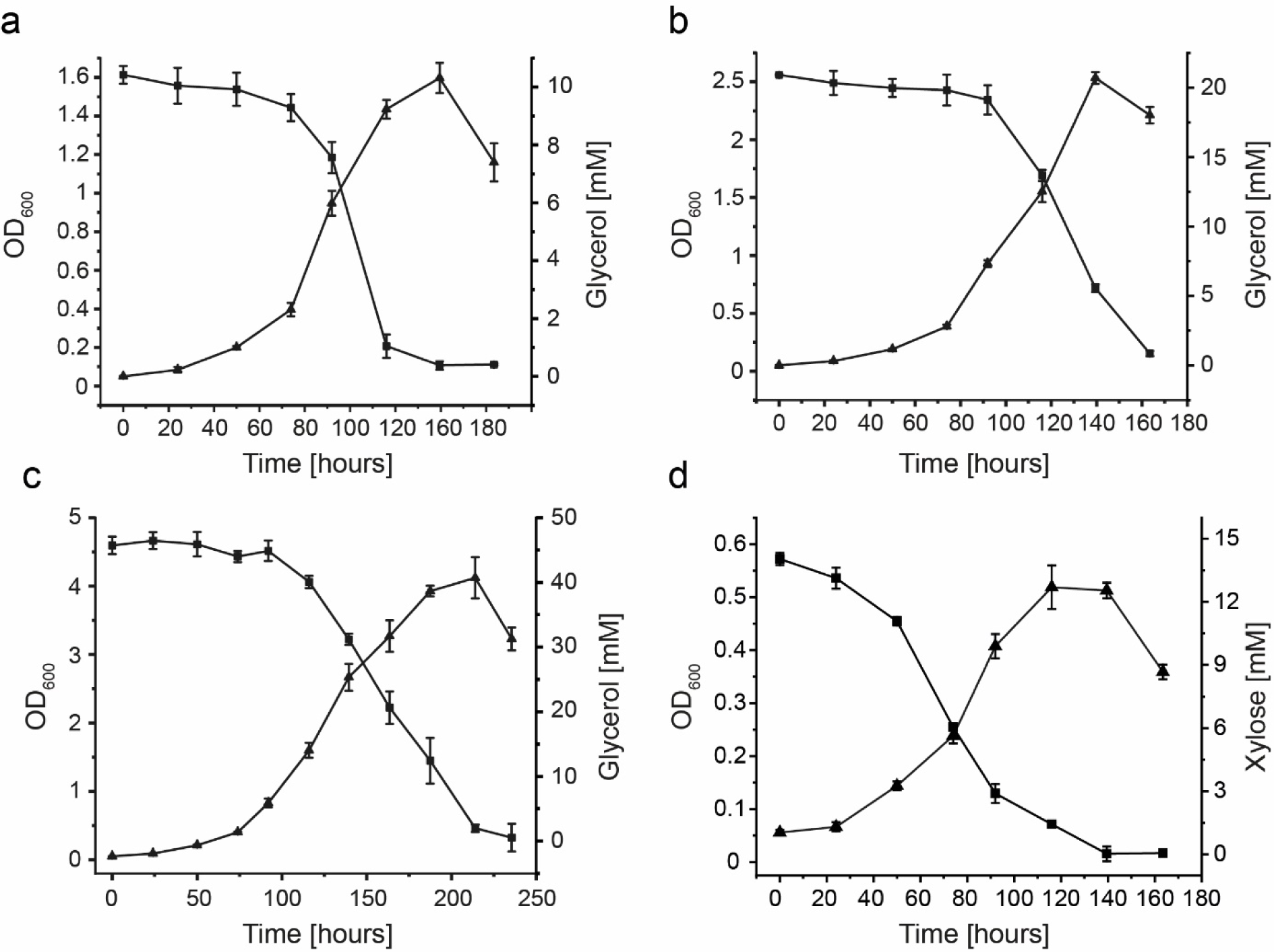
Growth and substrate consumption of *S. acidocaldarius* MW00G. Cells were grown on Brock’s basal medium containing 10 mM (a), 20 mM (b), and 40 mM (c) glycerol as sole carbon and energy source under aerobic conditions. For comparison the growth with 14 mM D-xylose (d) was studied. Growth was monitored as increase in OD_600_. Substrate concentrations in the cell free supernatant were determined enzymatically as described in the materials and methods part. Experiments were performed in triplicate and error bars indicate the SD of the mean.

To further elucidate the glycerol catabolism in *S. acidocaldarius* MW00G both, the carbon source dependent global transcriptional and translational response to glycerol compared to D-xylose was studied using transcriptomics (RNA-Seq) and proteomics (LC-MS-MS). In the presence of glycerol a total of 39 transcripts/proteins were upregulated while a total of 14 transcripts/proteins were downregulated with at least a log2-fold change of 2 (Supplementary Table 1). Downregulation was observed for the Weimberg pathway for pentose degradation (*saci_1938* – α-ketoglutarate semialdehyde dehydrogenase and *saci_1939* - 2-dehydro-3-deoxy-D-arabinonate dehydratase) as well as of the sugar binding subunit of the xylose/arabinose transporter *saci_2122*, while other central carbohydrate metabolic pathways (e.g. glycolytic branched Entner-Doudoroff pathway (ED), and tricarboxylic acid cycle (TCA), as well as the EMP for gluconeogenesis) were not regulated. A significant upregulation of the gene cluster *saci_2031* to *saci_2034* (Figure 3a) and its encoded proteins, a putative G3PDH (Saci_2032) with a downstream encoded protein (Saci_2031) annotated as carbon monoxide dehydrogenase subunit G (CoxG)-like, as well as a putative GK (Saci_2033) and a glycerol uptake facilitator (GUF, Saci_2034) was observed. Both gene couples *saci_2032-2031* and *saci_2033-2034* are divergently oriented as depicted in Figure 3b. In accordance with the observed regulation pattern of the gene cluster *saci_2031-2034*, we also observed the induction of the GK and the G3PDH on activity level in crude extracts of glycerol compared to D-xylose grown cells (Figure 3c). Whereas in D-xylose grown cells only a low GK activity of 0.06 U mg^-1^ and no G3PDH activity was determined, in glycerol grown cells a specific activity of 0.89 U mg^-1^ for GK and 0.12 U mg^-1^ for G3PDH using 2,6-Dichlorophenolindophenol (DCPIP) as artificial electron acceptor was measured. NAD^+^ dependent G3P oxidation was not detectable. Also, neither DCPIP nor NAD^+^ dependent glycerol oxidizing activity could be observed, indicating that no alternative pathway for glycerol dissimilation is present. Notably, the DCPIP dependent G3PDH activity was not only located in the soluble fraction (crude extract) after cell disruption but also in the resuspended pellet/membrane fraction with 0.23 U mg^-1^ (Figure 3c), whereas GK activity was exclusively found in the soluble fraction. In addition to *saci_2031-34*, we identified a second gene cluster comprising *saci_1117* and the divergently oriented *saci_1118* and *saci_1119*, encoding isoenzymes for GK (*saci_1117*), G3PDH (*saci_1118*) and the CoxG-like protein (*saci_1119*). In contrast to *Saci_2031-2034*, in this cluster a homologue encoding a GUF (or other transporter) is missing. However, *saci_1117-1119* were only slightly upregulated on transcript and protein level in response to glycerol.

**Figure 3:**
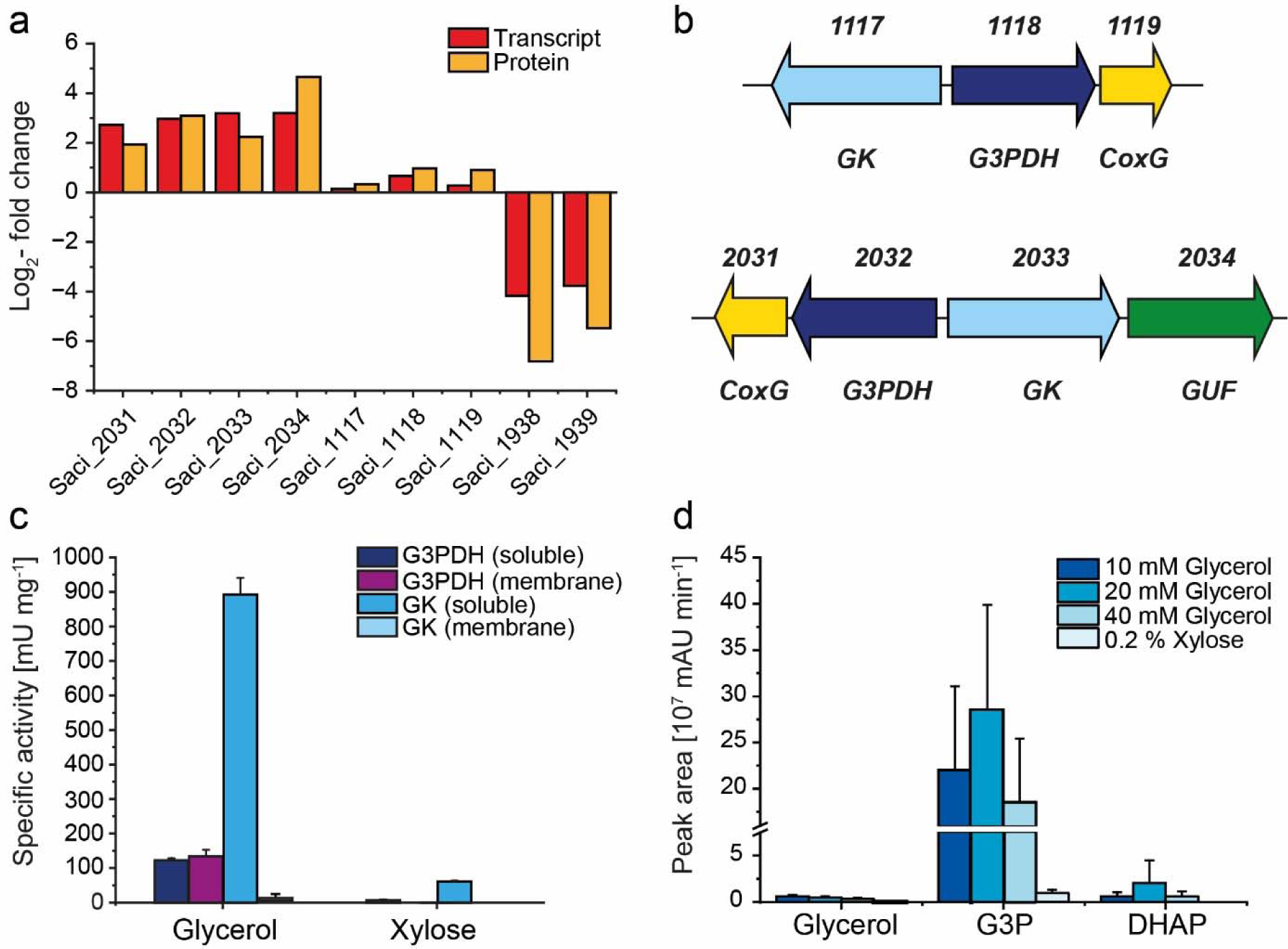
Poly-omics and activity-based analyses of glycerol degradation in *S. acidocaldarius*. (**a**) Log2 fold changes of transcript and protein levels of *saci_2031-saci_2034* and *saci_1938-saci-1939* of *S. acidocaldarius* MW00G grown on 40 mM of glycerol in comparison to 0.2 % (w/v) of D-xylose. (**b**) Divergent orientation of the potential glycerol degradation gene cluster *saci_2031-saci_2034* consisting of a putative glycerol uptake facilitator (GUF), glycerol kinase (GK, GlpK), glycerol-3-phosphate dehydrogenase (G3PDH, GlpA) and CoxG homologue. (**c**) GK and G3PDH activity of *S. acidocaldarius* MW00G grown on 40 mM glycerol in comparison to 0.2 % (w/v) D-xylose in the cytoplasmic and membrane fraction after cell lysis. (**d**) Peak areas of glycerol, G3P and DHAP of glycerol grown *S. acidocaldarius* MW00G (10mM, 20 mM and 40 mM glycerol) compared to D-xylose grown cells determined by targeted metabolomics. All values represent the average of three (growth curve and crude extract activities) or four (poly-omics analyses) independent measurements. Error bars represent the standard deviation of the mean.

Next, we performed comparative targeted LC(HILIC)-MS/MS based metabolome analyses of glycerol and D-xylose grown cells with a special focus on the intermediates of the central metabolic pathways like modified branched ED pathway, gluconeogenesis, TCA cycle, and of course the hypothetical intermediates of the glycerol catabolism. Interestingly, free glycerol was almost absent in glycerol grown cells, while the concentrations particularly of G3P and also of DHAP (∼20% compared to G3P) were significantly increased (Figure 3d). Additionally, in agreement to our transcriptome and proteome data no significant changes in metabolites in the upper EMP pathway for gluconeogenesis (i.e. glucose-6-phosphate (G6P), glucose-1-phosphate (G1P), fructose-6-phosphate (F6), and fructose-1,6-bisphosphate (F1,6BP)) and the common lower shunt of ED and EMP (i.e. glyceraldehyde-3-phosphate (GAP) and 2-phosphoglycerate (2PG)) were observed.

Together, these results show that *S. acidocaldarius* is able to utilize glycerol as sole carbon and energy source and that the glycerol uptake facilitator (Saci_2034, GlpF-like) is likely involved in glycerol transport via facilitated diffusion. Glycerol is then subsequently phosphorylated to G3P via GK (GlpK-like), and further oxidized by an at least loosely membrane associated G3PDH to yield DHAP which is than channelled either into the reversible, lower common shunt of the ED and EMP pathway for glycolysis or the upper EMP shunt for gluconeogenesis.

### Purification and characterization of GK and G3PDH isoenzymes

To confirm the function, C-terminally 10x His tagged GK (Saci_1117) and C-terminally Twin-Strep tagged GK (Saci_2033) were homologously produced in *S. acidocaldarius* MW001 since all expression efforts in *E. coli* were not successful. The putative GKs encoded by *saci_1117* and *saci_2033* comprise 498 and 497 amino acids with a calculated molecular mass of 55.6 and 55.3 kDa, respectively. This coincides well with the molecular mass of approximately 55 kDa experimentally determined under denaturing conditions by SDS-PAGE (supplementary Figure 1). The molecular weight under native conditions as determined via size exclusion chromatography was ∼110 kDa for both proteins, indicating a homodimeric structure (α_2_) (supplementary Figure 2). GK activity was confirmed for both isoenzymes and followed classical Michaelis-Menten kinetics. For Saci_1117 a *V*_max_ value of 45.7 U mg^-1^ and *K_M_* values of 0.024 mM (glycerol) and 0.205 mM (ATP) were determined (Figure 4, Table 1). The kinetic properties of Saci_2033 were similar with a *V*_max_ value of 88.2 U mg^-1^ and *K_M_* values of 0.024 mM (glycerol) and 0.174 mM (ATP). With glyceraldehyde (GA) and dihydroxyacetone (DHA) both enzymes showed only minor activities with catalytic efficiencies of <2% compared to glycerol (supplementary Figure 3a and b). For both GKs, the pH optimum was at 7 and the temperature optimum at 75°C (Table 1, supplementary Figure 3c and d). Furthermore, the Saci GKs were stable against thermal inactivation with half-life times of 45 h and 42 h for Saci_1117 and 12 h and 2 h for Saci_2033 at 70°C and 80°C, respectively. However, no remaining activity could be observed after 1 h of incubation at 90°C for Saci_2033 while Saci_1117 remained active over 3h of incubation (supplementary Figure 3e and f).

**Figure 4:**
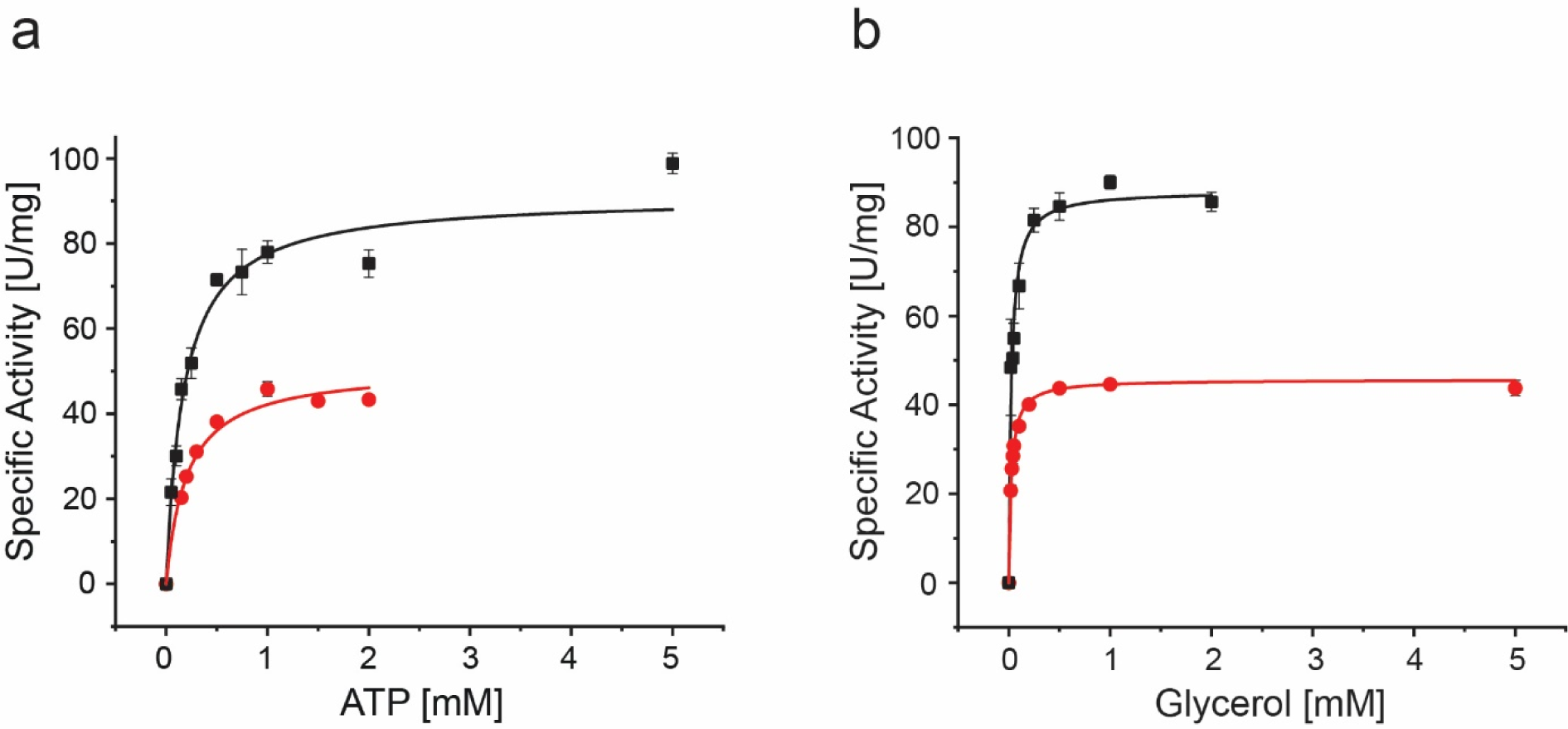
Kinetic characterization of the purified recombinant GK isoenzymes Saci_1117 and Saci_2033. The kinetic properties with ATP (**a**) and glycerol (**b**) for Saci_2033 (black squares) and for Saci_1117 (red circles) were determined in a continuous assay by coupling the glycerol-dependent ADP formation from ATP to NADH oxidation via PK and LDH at 50°C monitored as decrease in absorbance at 340 nm. Experiments were performed in triplicate and error bars indicate the SD of the mean.

**Table 1:**
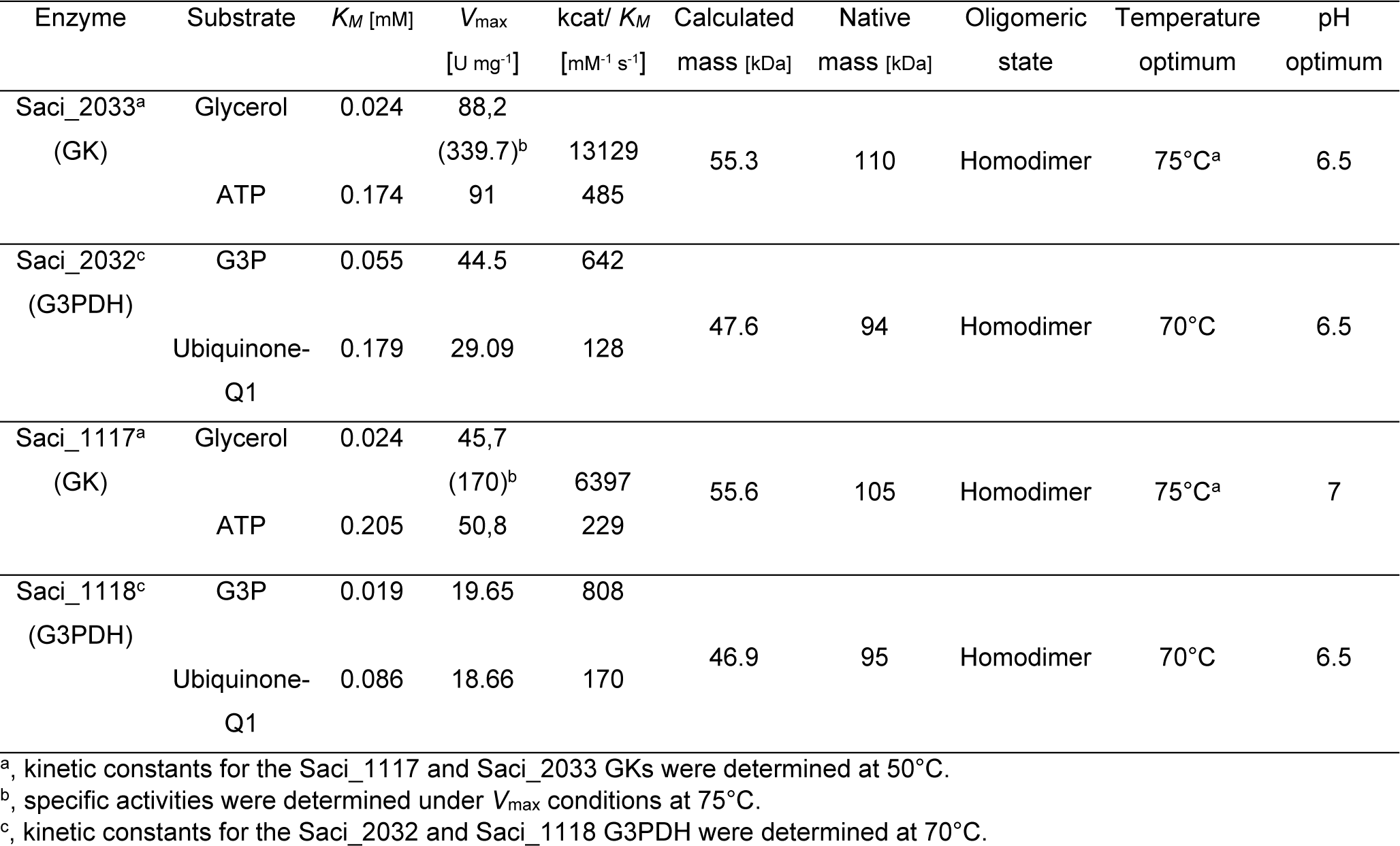
Kinetic characterization of the recombinant GK and G3PDH isoenzymes from *S. acidocaldarius*.

Other substrates like D(+)-glyceric acid, D-glucose, D-sorbitol, D-xylose, xylitol, and erythritol were not accepted as substrate. In addition to ATP also GTP and CTP could serve as phosphoryl donor, however with only 10% and 5% residual activity for Saci_1117 and 20% and 45% residual activity for Saci_2033, respectively (supplementary Figure 4a and b). Fructose-1,6-bisphosphate a known inhibitor of GK activity in e.g. *E. coli* (Applebee et al., 2011) had no effect on either Saci_1117 or Saci_2033 (supplementary Figure 4c and d).

The genes *saci_1118* and *saci_2032* are annotated to encode FAD-dependent oxidoreductases/G3PDHs consisting of 428 and 431 amino acids corresponding to a calculated molecular mass of 46.9 kDa and 47.6 kDa, respectively. Accordingly, heterologous expression in *E. coli* and purification of the N-terminally His-tagged G3PDHs yielded soluble, yellow proteins with subunit sizes of around 50 kDa (SDS-PAGE) and native molecular masses of 95 kDa for both enzymes (supplementary Figure 5) determined by size exclusion chromatography thus representing homodimers. The yellow colour already indicated the presence of a FAD cofactor and by spectrophotometric measurements (maximal absorption at 450 nm, extinction coefficient of 11.300 mM^-1^ cm^-1^, see materials and methods part) a FAD content of two for each of the homodimers was determined (Figure 5a). After reduction with G3P without the addition of an artificial electron acceptor the bound FAD cofactor remained stable in its colourless, reduced state. Furthermore, the direct electron transfer from FAD to oxygen forming hydrogen peroxide was excluded using the 2,2’-azinobis-3-ethylbenzothiazoline-6-sulfonate (ABTS) assay (data not shown). For Saci_2032 a *K_M_* of 0.055 mM for G3P and a *V*_max_ of 44.5 U mg^-1^ with DCPIP as electron acceptor was determined, while Saci_1118 showed a *K_M_* of 0.019 mM and a *V*_max_ of 19.7 U mg^-1^ (Figure 5b, Table 1). No activity could be detected with glycerol-1-phosphate (G1P), glycerol, glyceric acid, glyceraldehyde, glyceraldehyde-3-phosphate (GAP), and phosphoglyceric acid. Also, no activity could be observed with NAD^+^ or NADP^+^ as electron acceptor neither with G3P nor with glycerol, DHAP or glycerol-1-phosphate as substrate. However, ubiquinone-Q1 served as electron acceptor as followed by G3P dependent loss of absorption of the quinone at 280 nm (Figure 5c). *K_M_* and *V*_max_ values of 0.179 mM and 21.1 U mg^-1^ for Saci_2032 and a *K_M_* of 0.086 mM and *V*_max_ of 18.66 U mg^-1^ for Saci_1118 were determined with ubiquinone-Q1 (Figure 5d). Both enzymes showed a pH and temperature optimum of 6.5 and 70°C, respectively (Table 1, supplementary Figure 6a and b) as well as a high thermal stability without any loss in activity upon more than 6 h at 70°C. At 80°C both G3PDHs had a half-life time of around 3 h, and no remaining activity could be observed after 1 h of incubation at 90°C (supplementary Figure 6c and d).

**Figure 5:**
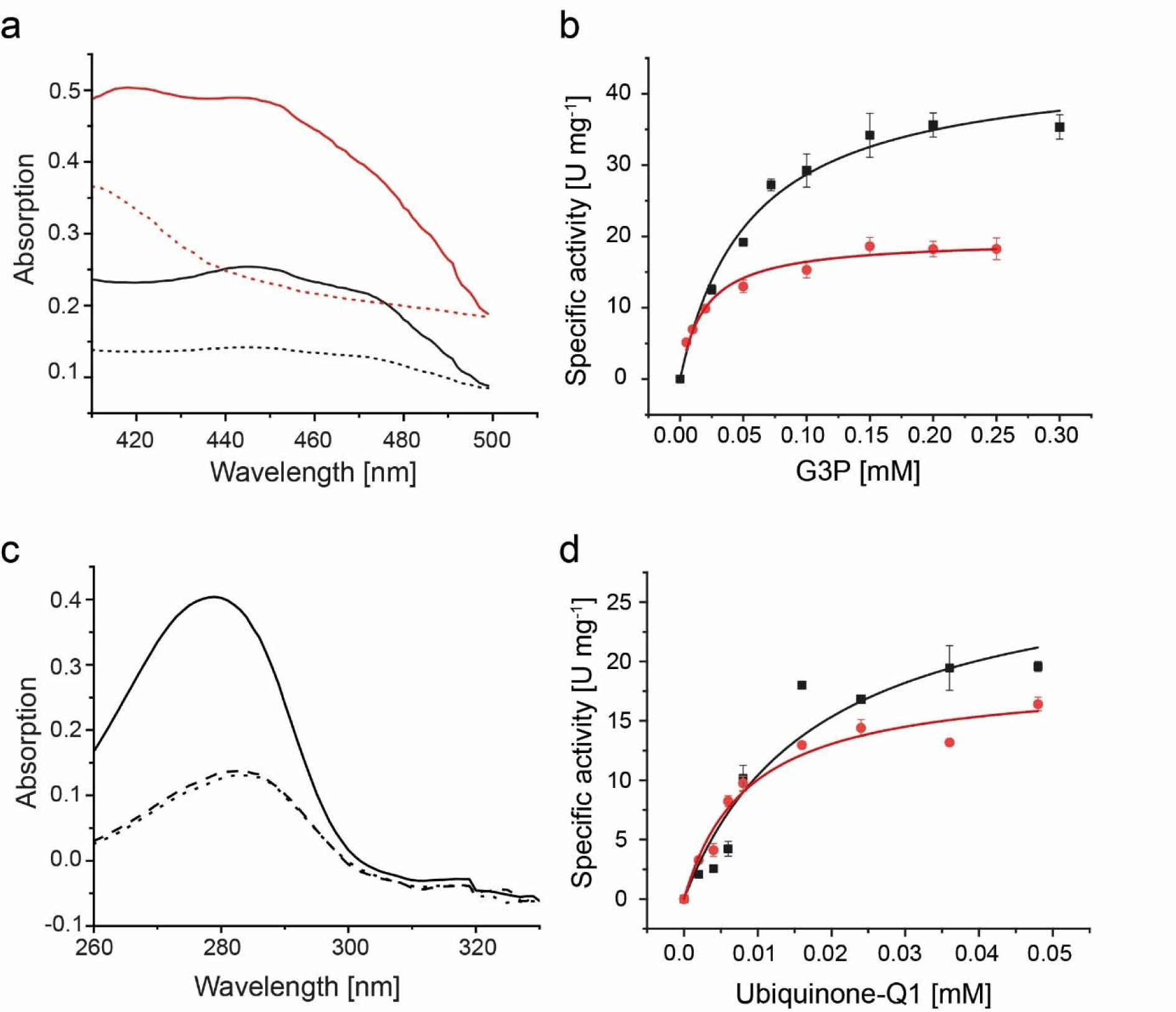
Biochemical and kinetic characterization of the recombinant G3PDH Saci_2032. (**a**) Absorption spectrum (400-500 nm) of Saci_2032 (black solid line) and Saci_1118 (red solid) bound FAD before addition of G3P. XX min after addition, loss of absorption at 450 nm indicates G3P dependent reduction of bound FAD in Saci_2032 (black dashed line) and Saci_1118 (red dashed line). Maximal absorption at 450 nm after denaturation by SDS was used to calculate the amount of bound FAD per protein. (**b**) The kinetic properties of recombinant Saci_2032 (black squares) and Saci_1118 (red circles) with G3P as substrate (0-0.3 mM) were determined in a continuous assay by coupling the oxidation of G3P to the reduction of the artificial redox active dye DCPIP as decrease of absorbance at 600 nm and 70°C. (**c**) Absorption spectra of ubiquinone-Q1 in the absence (solid line) and in the presence of Saci_2032 (dashed line) or Saci_1118 (dotted line) and 200 µM of G3P after 60 seconds. Loss of absorption at 280 nm indicates reduction of ubiquinone-Q1 by G3PDH. (**d**) Kinetic properties of Saci_2032 (black square) and Saci_1118 (red circle) towards ubiquinone-Q1 were determined in a continuous assay following the G3P dependent reduction of ubiquinone-Q1 to ubiquinol-Q1 at 280 nm and 70°C. Experiments were performed in triplicate and error bars indicate the SD of the mean.

### CoxG homologues serve as membrane anchor for *S. acidocaldarius* G3PDH isoenzymes

The ability of the Saci_2032 and Saci_1118 G3PDHs to reduce quinones raised questions about the mechanism of membrane association of both enzymes in *S. acidocaldarius*. As shown by the transcriptomics data, both *saci_2032* and *saci_1118* are cotranscribed and coregulated and thus forming an operon with *saci_2031 and saci_1119*, respectively, suggesting a functional association. However, both Saci_2032 and Saci_1118 G3PDHs were readily active *in vitro* in the absence of their corresponding CoxG-homologue Saci_2031 and Saci_1119, respectively, indicating that CoxG is not essential for G3PDH activity or electron transfer even to quinones (see above). Nevertheless, CoxG homologues have been implicated in membrane binding of few other proteins (Pelzmann et al., 2014; Wang et al., 2020) and therefore, we elucidated a potential role of Saci_2031 and Saci_1119 as membrane anchors for G3PDHs. We heterologously co-overexpressed *saci_2032/saci_2031* and *saci_1118*/*saci_1119* in the pETDuet-1-vector in *E. coli*. As controls, *saci_2032* and *saci_1118* were expressed alone from the same vector. Afterwards membrane and soluble proteins were separated by ultracentrifugation and the localization of His tagged Saci_2032 and Saci_1118 was analyzed via western blotting and immunodetection using anti-His antibodies. Only the co-expression of either *saci_2032* and *saci_2031* or *saci_1118* and *saci_1119* resulted in an enrichment of the respective G3PDH in the membrane fraction. In contrast, the expression of *saci_2032* and *saci_1118* alone in the absence of the corresponding CoxG did not yield any membrane associated G3PDH, instead the protein was exclusively localized in the soluble, cytoplasmic fraction in *E. coli* (Figure 6a and b). The signal detected in the cytoplasmic fraction for the G3PDHs in the cells co-expressing *saci_2032*/*saci_2031* and *saci_1118/saci_1119* might be due to a less efficient *coxG* expression compared to *saci_2032* and *saci_1118*, so that not all G3PDH could be recruited to the membrane.

**Figure 6:**
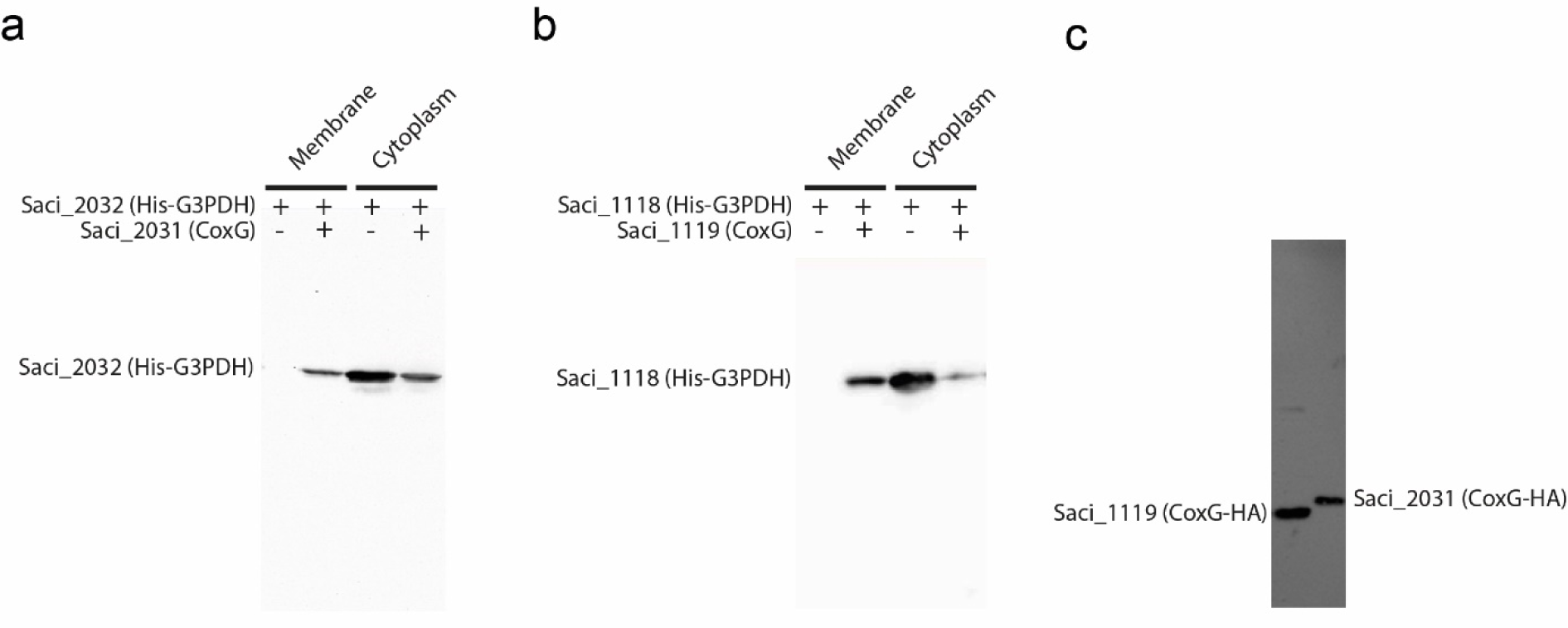
CoxG homologues Saci_2031 and Saci_1119 facilitate membrane binding of Saci_2032 and Saci_1118 G3PDH, respectively, *in vitro*. Effect of heterologous co-expression of CoxGs on the localization of His-tagged G3PDHs in the cytoplasmic and membrane fraction in *E. coli* monitored via western blotting and immunodetection with the anti-His antibody: Only upon co-expression with CoxG (Saci_2031) and G3PDH (Saci_2032) (**a**) as well as CoxG (Saci_1119) and G3PDH (1118) (**b**) the respective G3PDH is found in the membrane fraction. (**c**) Isolated membrane fractions obtained from *S. acidocaldarius* cells after homologous overexpression of HA-tagged CoxG homologues Saci_1119 (left lane) and Saci_2031 (right lane) using the pSVAaraFX-HA vector. Western blotting and immunodetection using the anti-HA antibody clearly show the localization of both CoxG homologues to the cell membrane.

Furthermore, upon homologous production of either Saci_2031 or Saci_1119 in *S. acidocaldarius* MW001, both HA-tagged proteins were found to a high degree in the insoluble/membrane fraction from which they could be solubilized using n-dodecyl β-D-maltoside (DDM). This finding provided further evidence that both CoxG homologues (Saci_2031 and Saci_1119) are membrane-associated in *S. acidocaldarius in vivo* (Figure 6c). To further confirm the interaction between the G3PDHs Saci_2032 and Saci_1118 and the CoxG homologues Saci_2031 and Saci_1119, respectively, coimmunoprecipitation experiments were performed using C-terminally HA-tagged CoxGs. MS analysis of interacting proteins revealed a specific interaction of Saci_2031 with Saci_2032 as well as of Saci_1119 with Saci_1118 (supplementary Table 2). Together these results strongly indicate that the interaction of G3PDHs with the membrane required for electron transfer from G3P to the (caldariella)quinone pool of the respiratory chain is mediated by the CoxG homologues Saci_2031 and Saci_1119, thus representing a novel function of CoxG homologues in Archaea and an unusual mechanism of membrane anchoring of G3PDH in *S. acidocaldarius*.

### Deletion of glycerol kinase genes affect growth on glycerol

To elucidate the significance of the individual glycerol kinase paralogues for glycerol conversion, single deletion mutants Δ*saci_1117* and Δ*saci_2033* as well as a double-deletion mutant Δ*saci_1117* Δ*saci_2033* were constructed in the parental strain MW00G.When grown on glycerol, the Δ*saci_1117* showed a diminished growth rate compared to the MW00G (Figure 7a), a slightly retarded glycerol consumption (Fig. 7b), and an only slightly reduced GK activity in crude extracts. In contrast, growth of Δ*saci_2033* as well as Δ*saci_1117* Δ*saci_2033* was completely abolished (Figure 7a) and the glycerol consumption was totally blocked (Figure 7b), indicating that Saci_2033 is essential for growth on glycerol. Since Δ*saci_2033* as well as Δ*saci_1117* Δ*saci_2033* did not grow on glycerol at all (and thus the effect of the deletion on GK activity in crude extracts could not be addressed), the effect of these GK deletions on the activity in crude extracts was determined in cells grown on NZ-amine. In Δ*saci_1117,* the GK activity was not affected compared to the parental strain MW00G (100 mU mg^-1^), whereas only minor (20 mU mg^-1^) and even no GK activity could be detected in Δ*saci_2033* and Δ*saci_1117* Δ*2033*, respectively, again confirming the major contribution of Saci_2033 to total GK activity (supplementary Figure 7).

**Figure 7:**
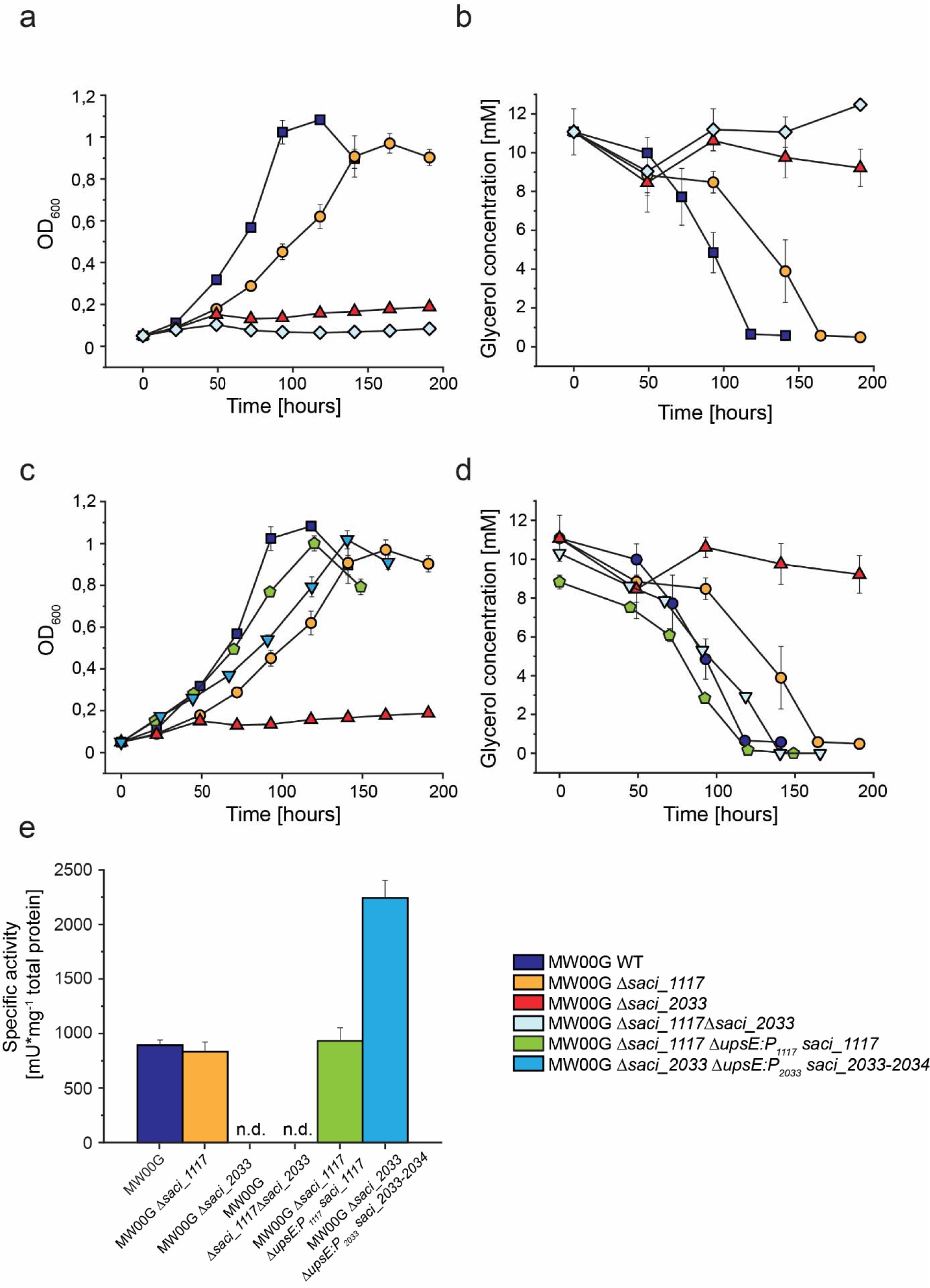
Comparison of growth, glycerol consumption and GK activities in *S. acidocaldarius* parental (MW00G), GK deletion and complementation strains. (**a**) Growth comparison of the parental strain MW00G with the single deletion strains Δ*saci_1117* and *Δsaci_2033* and the double deletion strain *Δsaci_1117 Δsaci_2033*. Cells were grown on Brock’s basal medium containing 10 mM glycerol as sole carbon and energy source and the respective glycerol consumption (**b**) was determined. Growth (**c**) and glycerol consumption (**d**) of the single deletion strains *Δsaci_1117* and *Δsaci_2033* complemented *in trans* by ectopic integration of the wildtype genes *saci_1117* and the *saci_2033/34* operon, respectively, into the *upsE* gene locus (*saci_1494*) under control of the respective native promoter. For comparison growth of mutants and parental strains as in (a) are shown. (**d**) GK activity measured in crude extracts from parental MW00G and deletion mutants as well as the complementation strains grown glycerol; n.d. indicates that activity could not be detected due to growth deficiency on glycerol. In the lower right panel the color code used in the whole Figure is depicted.

To restore the growth on glycerol, both single deletion strains Δ*saci_1117* and Δ*saci_2033* were complemented *in trans* by ectopic integration of the wildtype genes *saci_1117* and the *saci_2033/34* operon, respectively, into the *upsE* gene locus (*saci_1494*) under control of the respective native promoter. For both, Δ*saci_1117* and Δ*saci_2033* complementation could largely restore the growth on glycerol (Figure 7c) as well as substrate uptake to WT levels (Figure 7d) and crude extract activities (Figure 7e) Together with the growth studies, transcriptomics and proteomics analysis, as well as crude extract measurements, these results indicate that Saci_2033 is the primarily expressed and essential glycerol kinase during growth on glycerol whereas Saci_1117 has only minor importance.

## Discussion

*S. acidocaldarius* has previously been shown to cleave short chained triacylglycerols like tributyrin by means of esterases and was also shown to grow with short chain fatty acids like butyrate and hexanoate as sole carbon and energy sources (Zweerink et al., 2017; Wang et al., 2019). Therefore, the question arose whether this crenarchaeal model organism is also able to grow with glycerol, the other product of esterase mediated lipid breakdown. Herein, we demonstrated that (i) *S. acidocaldarius* grows with glycerol as sole carbon and energy source; (ii) The gene cluster *saci_2031-2034*, encoding CoxG, G3PDH, GK, and GUF were strongly up-regulated on transcript and protein level in the presence of glycerol (compared to D-xylose), while another gene cluster *saci_1117-1119* encoding a second set of paralogues of CoxG, G3PDH, GK was only slightly up-regulated; (iii) The activities of GK and G3PDH (with DCPIP as electron acceptor not with NAD) were highly up-regulated on glycerol but only residual (GK) or no activity (G3PDH) at all was measurable on D-xylose grown cells, and also activities of alternative pathways for glycerol utilization appeared to be absent; (iii) The intermediates G3P and also DHAP were much more abundant in glycerol grown cells; (iv) The enzymatic function of G3PDHs (Saci_2032 and Saci_1118) and GKs (Saci_2033 and Saci_1117), respectively, was confirmed by characterization of the recombinant proteins; (v) Only one of the two GKs GK (Saci_2033) was shown to be essential for growth on glycerol; (vi) The G3PDH activity was at least partly associated with the membrane which is mediated by the CoxG homologues Saci_2031 and Saci_1119. These results demonstrate that glycerol degradation in *S. acidocaldarius* proceeds via glycerol phosphorylation catalyzed by a “classical” GK followed by FAD-dependent G3P oxidation with electron transfer to the quinone of the respiratory chain carried out by a structurally unusual G3PDH. The resulting DHAP is then channeled into central metabolism (Figure 8).

**Figure 8:**
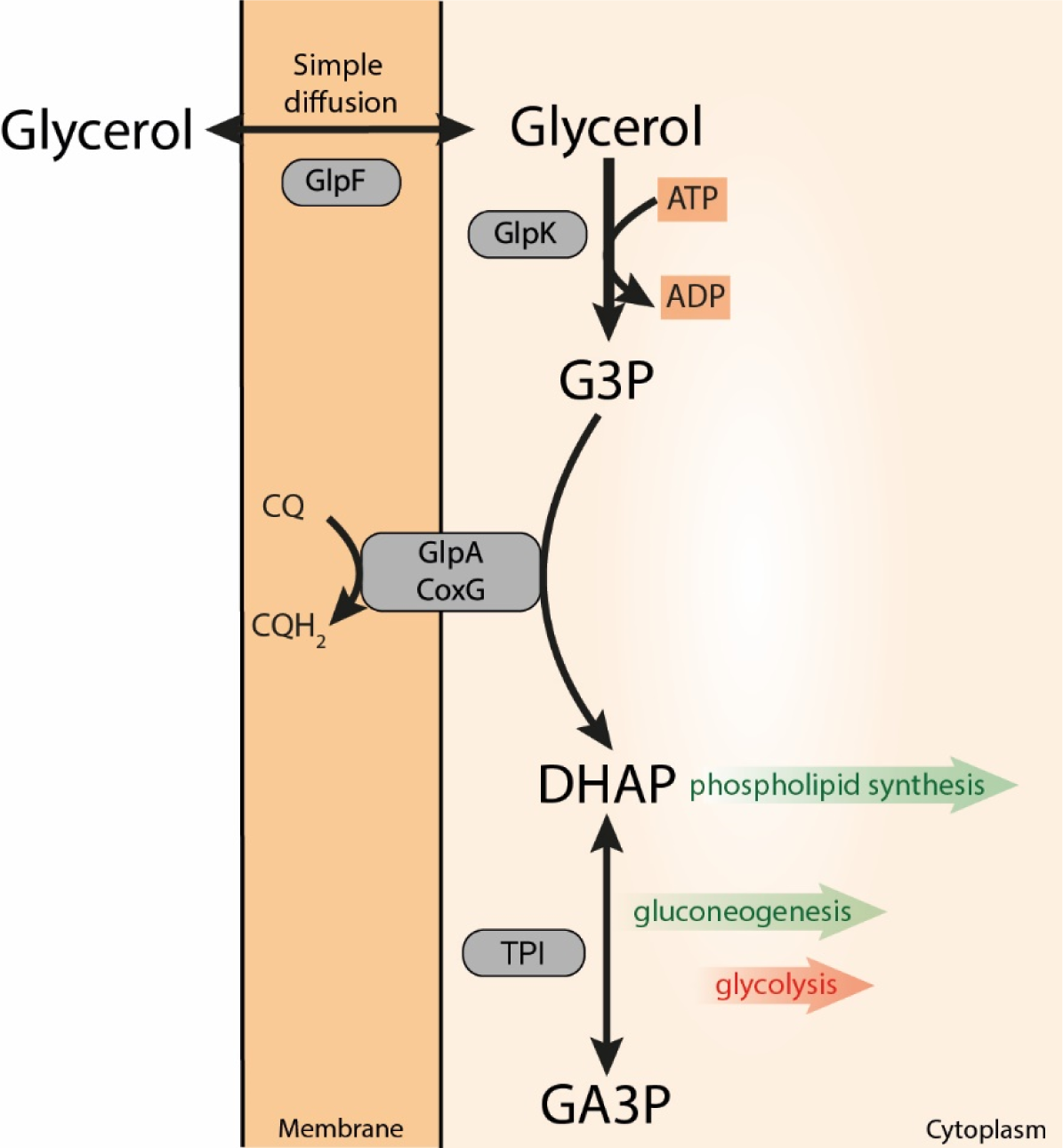
Glycerol metabolism in *S. acidocaldarius.* Glycerol uptake likely involves the GlpF-like glycerol uptake facilitator Saci_2034 and eventually simple diffusion through the cytoplasmic membrane. Additional uptake systems like the ABC transporter (Saci_1762-1765) might also contribute. Intracellularly, glycerol is phosphorylated by GK (Saci_2033), homologous to the bacterial GlpK, forming G3P which is then oxidized by a membrane bound, unusual GlpA-like FAD dependent G3PDH (Saci_2032). Saci_2032 is anchored to the membrane by the CoxG homologue Saci_2031 forming a complex that transfers electrons to the caldariellaquinone (CQ) yielding dihydroxyacetone phosphate (DHAP). DHAP can be used for phospholipid synthesis (via G1P in Archaea), and - with concomitant triosephosphate isomerase (TPI, Saci_0117) mediated conversion to glyceraldehyde-3-phosphate (GA3P) - for gluconeogenesis (green arrows), or glycolysis (red arrow). The paralogues Saci_1117 (GK), Saci_1118 (G3PDH), and Saci_1119 (CoxG) might contribute to glycerol degradation under different growth conditions (for details see discussion section).

Glycerol degradation in Archaea has so far only been analyzed in some detail in the halophile *Haloferax volcanii* which also grows on glycerol minimal media. Similarly, to *S. acidocaldarius*, the growth of *H. volcanii* on glycerol was better than on sugars (e.g. D-xylose) in terms of growth yield per mol of substrate carbon (Johnsen et al., 2009; Rawls et al., 2011). This higher growth yield and hence the higher ATP gain per mol of substrate is on the one hand due to the higher reduction state of glycerol compared to sugars like D-xylose but can likely not account alone for the observed yield differences and might hence hint at a more efficient energetic coupling during growth on glycerol. Accordingly, in the transcriptomics and proteomics data we observed changes in the respiratory chain of *S. acidocaldarius* with the SoxEFGHIM (*saci_2258-saci_2263*), one of three terminal oxidases, being strongly upregulated (supplementary Table 1). This terminal oxidase was shown to have a higher H^+^/e^-^ ratio (Gleißner et al., 1997; Schafer et al., 1999; Komorowski et al., 2002; Bischof et al., 2019) and thus likely additionally contribute to the higher growth yield. In contrast to the results presented herein, *S. acidocaldarius* has been previously reported not to grow on glycerol (Grogan, 1989) and in agreement G3PDH homologues seemed to be absent in Sulfolobales (Villanueva et al., 2017). These findings can likely be explained by the prolonged time the organism required for adaption to glycerol as carbon and energy source observed in this study and at sequence level by the C-terminal truncation/differences of the G3PDH of *S. acidocaldarius* and other Sulfolobales (as well as some Thermoprotei and Thermoplasmatales) (see below). Such an adaptation phase - although much less pronounced – was also reported for *Pseudomonas* spp. and was attributed to the mode of transcriptional regulation with G3P as inducer (Poblete-Castro et al., 2020). Like *S. acidocaldarius*, *H. volcanii* utilizes the GK/G3PDH pathway for glycerol degradation. In *H volcanii* high constitutive activity levels of both GK and G3PDH (∼250 mU mg^-1^ and 28 mU mg^-1^ respectively) were observed in absence of glycerol which were roughly two fold up-regulated in its presence (Nishihara et al., 1999; Sherwood et al., 2009). In *S. acidocaldarius* only residual GK activity (<10%) and no G3PDH activity was observed in the absence of glycerol when D-xylose was the growth substrate. This different expression pattern compared to *H. volcanii* might reflect that glycerol in halophilic habitats is regarded as main and preferred carbon and energy source (see references in (Sherwood et al., 2009)) which is likely different in thermoacidophilic environments.

In accordance with previous findings (Yokobori et al., 2016; Villanueva et al., 2017), bioinformatic/phylogenomic analyses revealed that – in addition to Halobacteriales and Sulfolobales - also other archaeal lineages harbor the GK/G3PDH pathway. However, the GK/G3PDH homologues and hence the potential ability to utilize glycerol is present only in some representatives of these lineages and appears thus only patchily distributed in Archaea (Villanueva et al., 2017).

### Glycerol uptake

For glycerol uptake, *S. acidocaldarius* harbors a GUF homologous to GlpF (Stroud et al., 2003) which is up-regulated in response to glycerol and thus likely involved in glycerol uptake. In Archaea, GlpF homologues seems to be restricted to few *Sulfolobus* and *Saccharolobus* spp. as well as to halophiles (Rawls et al., 2011; Williams et al., 2017) and are thus much less abundant then GlpA homologues in Archaea meaning that by far not all Archaea harboring GK and G3PDH also contain a GlpF homologue. This might indicate that alternative transporters than GlpFs are utilized for glycerol uptake in other Archaea and the gene neighborhoods of the GK/G3PDH encoding genes often also comprise e.g. putative MFS (major facilitator superfamily) transporter encoding genes (e.g. in *Thermococcus* and *Pyrococcus* species, some Thermoplasmatales, *Vulcanisaeta*). Alternative glycerol transporters are known in *Mycoplasma spp., Rhizobium* spp. and other α proteobacteria employing ABC transporters (in addition to a GlpF homologue) with implications for transport efficiencies and pathogenicity (Ding et al., 2012; Blötz and Stülke, 2017; Mahdizadeh et al., 2021). In Eukarya, also H^+^ (Na^+^) symport systems have been described especially in yeast (Wille et al., 1998; Klein et al., 2017). In *S. acidocaldarius,* in addition to the *glpF* gene, also a putative ABC transporter (*saci_1762-1765*) was up-regulated which could thus also be involved in glycerol transport. In addition, glycerol uptake via simple diffusion might also take place in *S. acidocaldarius* and other Archaea. For example, in *E. coli* deletion of the *glpF* gene had no severe impact on glycerol consumption under high concentrations. Nevertheless, below 5 mM glycerol in the medium the growth rate remarkably decreased indicating that the facilitator is especially important at low substrate availability (Richey and Lin, 1972). Further on, besides the concentration gradient a major driving force of the glycerol uptake was shown to be the subsequent ATP-dependent phosphorylation catalyzed by the GK (Voegele et al., 1993; Poblete-Castro et al., 2020). Accordingly, intracellular glycerol was nearly undetectable in *S. acidocaldarius*, whereas the G3P and to a lesser extend also the DHAP concentrations were significantly increased during growth on glycerol compared to D-xylose. Furthermore, the GK and G3PDH activities in crude extracts were clearly induced to 0.89 U mg^-1^ and 0.12 U mg^-1^, respectively. In D-xylose grown cells GK activity was reduced by more than 90% to 0.06 U mg^-1^ and G3PDH activity was not detectable at all. Similar GK activities were reported for *H. volcanii* (0.43 U mg^-1^) (Sherwood et al., 2009) or for *E. coli* (0.61 U mg^-1^) (Richey and Lin, 1972) and for the latter a further activation of the GK activity through interaction with the GlpF was suggested (Voegele et al., 1993). The GK and G3PDH activities agree with the much more pronounced accumulation of G3P compared to DHAP in *S. acidocaldarius* and also show that both activities are not rate limiting. The transcriptomics and proteomics analyses revealed that candidates for further DHAP conversion via glycolysis and gluconeogenesis as well as the TCA cycle are not regulated in response to glycerol which thus likely accounts for the accumulation of G3P and DHAP.

### Properties of the glycerol kinase isoenzymes Saci_1117 and Saci_2033

The *S. acidocaldarius* GKs like all other GKs characterized so far belong to the FGGY family within the sugar kinase/HSP70/actin superfamily (Lu et al., 2020) and show a high degree of sequence conservation not only to other archaeal homologues but also to bacterial (>50% sequence identity) and even eukaryotic (∼44%) enzymes and thus across the domains of life. The homodimeric structure of Saci_1117 and Saci_2033 is well in accordance with some other GKs from eukaryotes (e.g. *Trypanosoma brucei gambiense*, *Plasmodium falciparum*, *Chaetomium thermophilum* (Schnick et al., 2009; Balogun et al., 2013; Wilk et al., 2020)) and bacteria (e.g. *Cellulomonas* sp. (Fukuda et al., 2016)). Also, homotetrames (for some Bacteria like *Thermus thermophilus*, *Elizabethkingia meningoseptica*, and also some Eukaryotes like *Saccharomyces cerevisiae* (Thorner and Paulus, 1973; Huang et al., 1997; SAKASEGAWA et al., 1998)) and even homohexamers (*T. kodakarensis* (Hokao et al., 2020)) have been reported. For *E. coli* a dimer-tetramer equilibrium in solution was described (Applebee et al., 2011) and also for *Haemophilus influenzae* a higher (>4mer) and a lower (<4mer) oligomerization state was reported (Pawlyk and Pettigrew, 2001). Two different oligomeric species were clearly not observed for Saci_1117 and Saci_2033 which eluted solely as homodimer in size exclusion chromatography experiments (supplementary Figure 2). The tetramer formation observed for the *E. coli* GK appears as prerequisite for allosteric regulation by fructose-1,6-bisphosphate (FBP) (Applebee et al., 2011). Thus, the homodimeric structure of both *S. acidocaldarius* GKs appears in accordance with the missing FBP regulation. Also, for the dimeric enzymes from *P. falciparum* and *T. brucei gambiense* no allosteric control with FBP was observed (Schnick et al., 2009; Balogun et al., 2014). Similarly, the *T. kodakarensis* GK initially described as a homodimer did not respond to FBP as allosteric effector. Later on, it has been shown that in presence of glycerol the *Thermococcus* enzyme assembles into hexamers but the subunit interactions appear different than observed in the tetramer interactions in *E. coli* essential for FBP binding and seems thus also in agreement with the missing FBP regulation (Koga et al., 2008; Hokao et al., 2020). The glycerol mediated hexamer formation of the *Thermococcus* GK increases the ATP affinity of the enzyme tenfold. Under the same glycerol concentration, however, the Saci_1117 and Saci_2033 GKs did not change their oligomeric structure (supplementary Figure 2) and the lysine 271 (K271) in *T. kodakarensis* enzyme essential for hexamer formation is not conserved in Saci_1117 and Saci_2033. Additionally, from *Enterobacteriaceae* (e.g. *E. coli*) and *Firmicutes* (e.g. *Enterococcus, Streptococcus, Bacillus*), two different GK regulation mechanisms involving components of the PEP-dependent phosphotransferase system (PTS) for sugar transport have been described (for literature see (Yeh et al., 2009)). In *E. coli* the unphosphorylated Enzyme IIA^Glc^ acts as allosteric GK inhibitor and in firmicutes GK is phosphorylated by HPr∼P at a conserved histidine residue (H232 in *Enterococcus casseliflavus)* located in the loop region where also FBP is bound (for literature see (Yeh et al., 2009)). This phosphorylation in the absence of PTS sugars stimulates the enzyme activity 10-15-fold. Similar regulatory mechanisms involving components of the sugar transport PTS systems are unlikely to occur in *S. acidocaldarius* since PTS systems in this organism and in Archaea in general are absent despite in some halophiles (Pickl et al., 2012). Interestingly, in *H. volcanii* one PTS component protein, a HPr homologue encoded by the ptsH2 gene (HVO_1543), is part of the chromosomal glycerol gene cluster (Sherwood et al., 2009; Rawls et al., 2011). However, the phosphorylated H232 in the *E. casseliflavus* enzyme is not conserved in the *H. volcanii* GK (and also not in Saci_1117 or Saci_2033).

The kinetic constants of the Saci_1117 and Saci_2033 GKs (Table 1) are well in the (rather broad) range of those determined for other GKs. The *V*_max_ of the Saci_1117 and Saci_2033 GK at a more physiological temperature of 75°C was determined at 170 U mg^-1^ and 340 U mg^-1^, respectively, which roughly approximates a similar range reported for other GKs (for the kinetic constant of selected GKs see supplementary Table 5). A relaxed substrate specificity as observed in both *S. acidocaldarius* GKs converting also DHA and GA although with much lower catalytic efficiencies (<2%) (supplementary Figure 3) was also reported e.g. for the *E. coli* enzyme (Hayashi and Lin, 1967). The temperature optimum (75°C) and the thermal stability with half-life times (t_1/2_) of 45 h and 12 h (70°C) and 42 h and nearly 2 h at (80°) determined for Saci_1117 and Saci_2033, respectively. fit well with the growth conditions of *S. acidocaldarius*. Nevertheless, in accordance with the growth optimum of *T. kodakarensis* of 85°C the thermal properties of the GK from this euryarchaeon are more pronounced (T_opt_ 80°C, t_1/2_ 30 min (100°C)) (Koga et al., 1998; Atomi et al., 2004). The GK from the thermophilic bacterium *Thermus thermophilus* (optimal growth temperature around 68°C) on the other hand showed slightly lower T_opt_ between 50°C and 70°C and lost ∼75% activity upon 30 min incubation at 70°C (Huang et al., 1997).

### Only Saci_2033 GK is essential for glycerol degradation

The *S. acidocaldarius* genome encodes two gene clusters for glycerol metabolism, *saci_2031-2034* (encoding CoxG homologue, G3PDH, GK and GUF, respectively) and *saci_1117-1119* which differ in the presence of the GUF encoding gene *saci_2034* which is missing the *saci_1117-1119* gene cluster. Beside the missing GUF gene both glycerol gene clusters show a similar organization (Figure 3). However, only the *saci_2031-2034* gene cluster was shown to be significantly up-regulated in response to glycerol on transcript and protein level whereas *saci_1117-1119* is only slightly induced if at all. Furthermore, only the GK Saci_2033 was shown to be essential for growth on glycerol whereas the Saci_1117 GK is dispensable suggesting that the *saci_2031-2034* gene cluster is mainly involved in glycerol degradation at least under the tested conditions.

The presence of two GK/G3PDH couples appears scarce in Archaea and also in Bacteria as revealed by database searches. The presence of two G3PDHs but only one GK seems more widespread, either of two different G3PDHs, like GlpABC and GlpD in e.g. *E. coli* and other facultatively anaerobic bacteria (Lin, 1976; Schryvers et al., 1978; Schryvers, 1981; Austin and Larson, 1991; Yeh et al., 2008; Poblete-Castro et al., 2020), or two GlpABC isoenzymes like reported for e.g. *H. volcanii* (and some other haloarchaea) (Rawls et al., 2011; Williams et al., 2017). The presence of two GK paralogues appears even more unusual and has been described so far for *Lactobacillus rhamnosus* and *Pseudomonas aeruginosa* (de Fátima Alvarez et al., 2004; Tang et al., 2023). In the archaeal domain, also few haloarchaea were found to harbor two GK copies (Williams et al., 2017). As observed herein for the *saci_2033* encoded GK in *S. acidocaldarius*, in studies on the role of two GK isoenzymes in *L. rhamnosus* as well as on the function of two G3PDH (GlpABC) isoenzymes in H. volcanii, only one of the paralogues was shown to be essential and up-regulated during growth on glycerol whereas the other is dispensable at least under the tested conditions, although the respective activity has been confirmed for both (de Fátima Alvarez et al., 2004; Rawls et al., 2011). In contrast to the essential paralogue in both cases cooccurring with other glycerol related genes in the genome encoding G3PDH or GK, respectively, as well as with the GUF, the second paralogue is not colocalized with glycerol genes.

For the two G3PDHs, GlpD and GlpABC, e.g. in *E. coli* it is well known that they are primarily expressed under different environmental settings, i.e. under aerobic and anaerobic conditions, respectively (see above). Also, for *P. aeruginosa* it has been reported that the two GK paralogues belong to different regulons (Tang et al., 2023). Here, the enzyme non-essential for glycerol catabolism has also significantly lower GK activity and might thus be either required for glycerol breakdown under different conditions or might have even acquired a different metabolic function reflected by different evolutionary history, which has in turn be discussed to improve the fitness of the organism (Tang et al., 2023).

Since the two G3PDH and GK paralogues in *S. acidocaldarius* show very similar functional parameters it appears reasonable to speculate that both might be responsible for glycerol degradation under different growth conditions. In this respect, it is interesting to note that *saci_1117-1119* gene cluster is located in a larger array of genes (*saci_1103-1126*) containing mainly fatty acid β oxidation related genes as well as two esterases which were shown to be regulated by a newly identified TetR-family transcription factor (Wang et al., 2019). Thus, a role of the *saci_1117-1119* gene cluster in glycerol utilization during lipid breakdown might be envisaged whereas the *saci_2031-2024* gene cluster appears responsive to free glycerol (or to lipids containing FAs of different chain length). Interestingly, the regulator of the *saci_1103-1126* gene region was shown to bind to fatty acyl-CoA esters with increasing binding affinity to longer chain acyl-CoAs (Wang et al., 2019). However, the different functions of both glycerol gene clusters in *S. acidocaldarius* remains to be further analyzed.

### Properties of the G3PDHs Saci_2032 and Saci_1118 and an unusual mode of membrane anchoring by CoxG (Saci_2031 and Saci_1119)

In contrast to the *S. acidocaldarius* GK/G3PDH pathway, the route in *H. volcanii* involves a G3PDH composed of all three GlpABC subunits homologous to the “anaerobic” route known from many especially facultatively anaerobic bacteria like *E. coli* (Lin, 1976; Cole et al., 1988; Varga and Weiner, 1995; Rawls et al., 2011; Poblete-Castro et al., 2020). In Bacteria and haloarchaea, the GlpC and GlpB subunits are supposed to permit membrane anchoring and electron transfer, respectively, from the catalytic subunit GlpA finally to the (mena)quinone pool in *E. coli* (Cole et al., 1988; Varga and Weiner, 1995). However, except for most haloarchaea the GlpB and C subunits are absent from nearly all other Archaea and also GlpD and GlpO homologues were not identified. These organisms in addition to GlpK (and frequently also GlpF) only contain GlpA homologues which were however demonstrated to be active without additional subunits herein for *S. acidocaldarius* and also previously for *T. kodakarensis* (Koga et al., 2019). Compared to the *S. acidocaldarius* G3PDHs, the TK1393 G3PDH catalytic subunit from *T. kodakarensis* alone showed a 4-5 fold (Saci_1118) and a 7-8-fold (Saci_2032) lower *V*_max_ of 5-6 U mg^-1^ and a 50-to more than 100-fold higher *K_M_* value for G3P (2.6 mM vs. 0.055 mM (Saci_2032) and 0.019 mM (Saci_1118)). However, the G3PDH encoding genes in Archaea are often downstream co-localized with different likely functionally related genes (for detailed discussion see below) which suggest – together with the absence of the GlpB and C subunits - that electron transfer in these archaeal species might be different. We herein forwarded evidence that the *S. acidocaldarius* G3PDHs form a catalytically active dimer which can directly reduce the (ubi)quinone analogue Q1, two features that are similar to characterized bacterial and eukaryotic GlpDs (Unemoto et al., 1981; Rauchová et al., 1997; Yeh et al., 2008) which directly transfer the electrons to the quinone pool. The use of quinones (E^0^’ ∼+110 mV (ubiquinone) and −75mV (menaquinone)) as acceptor for the relatively high potential electrons of G3P (∼-200 mV) oxidation is advantageous since NAD(P)^+^ (−320 mV) would render this reaction endergonic (Thauer et al., 1977). Also, the FAD content of one per monomer is more similar to the GlpDs. In contrast, the catalytic heterodimer from *E. coli* (GlpAB) was described to contain only one FAD and also an FeS cluster both likely located in the GlpA subunit. How the GlpB subunit acts in the electron transport is not fully understood yet. Sequence analyses suggest that the GlpB subunit, which is homologous to several flavoproteins like the FAD subunit of the succinate dehydrogenase, might contain a further flavin cofactor involved in electron transport to the FeS cluster(s) in GlpC (Cole et al., 1988).

The *S. acidocaldarius* G3PDHs showed similar specific activities with DCPIP as artificial electron acceptor as described for the GlpD from *Vibrio alginolyticus* with the artificial redox carriers PMS-MTT or ferricyanide (see supplementary Table 6). For the GlpDs from *E. coli* a roughly ten-fold lower activity with DCPIP has been reported which was even lower for the pig brain mitochondrial enzyme (0.8 U mg^-1^). However, for the pig brain enzyme also the kinetic constants for the more native substrates, the ubiquinones-Q0 and -Q1, have been determined. For the ubiquinone-Q1 both the *K_M_* and *V*_max_ values were more than 10-fold lower (*V*_max_ 2.6 U mg^-1^, *K_M_* 0.013 mM) (Cottingham and Ragan, 1980) compared to the *S. acidocaldarius* enzymes (*V*_max_ 21 U mg^-1^, *K_M_* 0.18 mM (Saci_2032); *V*_max_ 18.7 U mg^-1^, *K_M_* 0.086 mM (Saci_1118)) resulting in a similar catalytic efficiency. The high substrate specificity observed for the *S. acidocaldarius* G3PDHs not converting G1P, glycerol, glyceric acid, GAP, and phosphoglyceric acid has also been reported for two isoenzymes from *Acidiphilium* sp. (Hatta et al., 1989).

However, as further detailed below the *S. acidocaldarius* G3PDHs are more similar to GlpAs. So far, the GlpAB active heterodimer from *E. coli* is the only bacterial GlpA-like enzyme which has been kinetically characterized in some detail. The *V*_max_ value reported for the *E. coli* enzyme (34 U mg^-1^) is in a similar range as the *S. acidocaldarius* G3PDHs homodimers (44.5 U mg^-1^ and 19.6 U mg^-1^) although the *K_M_* for G3P is 6-fold and 15-fold lower for *S. acidocaldarius* G3PDHs (0.055 mM (Saci_2032) and 0.019 mM (Saci_1118) vs. 0.34 mM for the *E. coli* GlpAB heterodimer) (Schryvers and Weiner, 1981). Conversely, the direct reduction of quinone analogues was not described for the *E. coli* enzyme likely because it relies on the GlpC subunit for both, membrane anchoring and quinone reduction (electron transfer) (also underscored by the presence of a FeS cluster(s) in the C subunit) (Varga and Weiner, 1995). In *S. acidocaldarius* the gene couples encoding G3PDHs and the CoxG homologues, i.e. saci_2032/2031 and saci_1118/1119, were shown to form operons and the proteins encoded by each of the two operons were shown to interact with each other. The CoxG homologue recruits the G3PDH active dimer to the membrane, but as demonstrated by the quinone reactivity of bothG3PDH dimers alone, the CoxG (subunit) is not involved in electron transfer and thus only anchors the protein to the membrane. The only functionally analyzed CoxG homologue so far is the protein from *O. carboxydovorans* which recruits the carbon monoxide dehydrogenase (from the aerobic xanthin oxidase type) to the membrane but does not seem to have any enzymatic implications (Pelzmann et al., 2014). A similar function was discussed for the 3-hydroxypyridine dehydrogenase of *Ensifer adhaerens* HP1 where a CoxG homologue was proposed to facilitate membrane binding (Wang et al., 2020).

In contrast, the homodimeric “aerobic” G3PDH (GlpD), e.g. from *E. coli*, does not require any membrane anchoring and/or additional electron transferring subunits but as a monotopic membrane protein directly interacts with the membrane and transfers electrons to the ubiquinone pool. Furthermore, these GlpDs were also shown to require membrane interaction, or phospholipids or nondenaturing detergents for conformational integrity and thus full activity ((Yeh et al., 2008) and the literature cited therein). This was not the case for the *S. acidocaldarius* G3PDHs and might further reflect the different mode of membrane association (supplementary Figure 8).

### Sequence and structural comparisons of G3PDHs

As indicated by sequence comparisons (exemplarily shown for Saci_2032 in Figure 9), conserved domain analyses, and structural comparison including modeling using the “ColabFold: AlphaFold2 using MMseqs2” online tool (Mirdita et al., 2022) (supplementary Figure 9), the Saci_G3PDH and other archaeal homologues mentioned above are homologous in their first N-terminal ∼360 amino acids with GlpDs, GlpAs and also with GlpOs. This part of the proteins constitutes the FAD-dependent oxidoreductase domain (pfam01266) which belong to the D-amino acid oxidase (DAAO) superfamily (including e.g. also the glycine oxidase from *Bacillus subtilis* (1ng3) (Settembre et al., 2003)). These proteins are composed of a ‘glutathione-reductase-2’ type FAD-binding domain and an antiparallel β-sheet based substrate-binding domain (for details and literature see (Elkhal et al., 2015)). Accordingly, the FAD binding residues and also the G3P binding site are quite conserved (Figure 9). However, GlpD, GlpO, and GlpA proteins as well as the archaeal homologues differ in length and domain organization of their C-terminus. Glycine oxidases just constitute the DAAO fold without any C-terminal extensions. However, as revealed by the crystal structures, the C-terminus in GlpD from *E. coli* (2qcu) forms a truncated G3P oxidase domain (pfam 16901, composed of roughly 120 amino acids) shortened by a twenty amino acids comprising helix in its very C-terminal end compared to the complete G3P oxidase domain (140 aa) present in GlpO from *Streptococcus* sp. (2rgo), (supplementary Figure 9 as well as supplementary Figure 10a and b). In addition to the complete C-terminal G3P oxidase domain, GlpOs differ from GlpDs by the presence of a 50 amino acid insertion in the C-terminal part of the DAAO fold (Figure 9). In contrast to GlpDs and GlpOs, the C-terminus of GlpAs is even larger (160 aa) containing a bfd-like domain with an FeS cluster (pfam04324) (Marchler-Bauer and Bryant, 2004; Yao et al., 2012; Wijerathne et al., 2018; Lu et al., 2020) and the four highly conserved cysteine residues likely involved in FeS cluster binding are clearly visible in the C-terminus of GlpA sequences but missing in GlpDs and GlpOs (Figure 9, supplementary Figure 11). The presence of such an FeS cluster is in accordance with the presence of two non-heme irons per catalytically active GlpAB subunit early reported (Schryvers and Weiner, 1981). As indicated by structural predictions the C-terminal domain of GlpA resembles in its first part the C-terminal domain of GlpOs (and thus also of gGlpDs) but is even further extended by an additional α helix in its very C-terminal end compared to GlpO (supplementary Figure 9 and 10).

**Figure 9:**
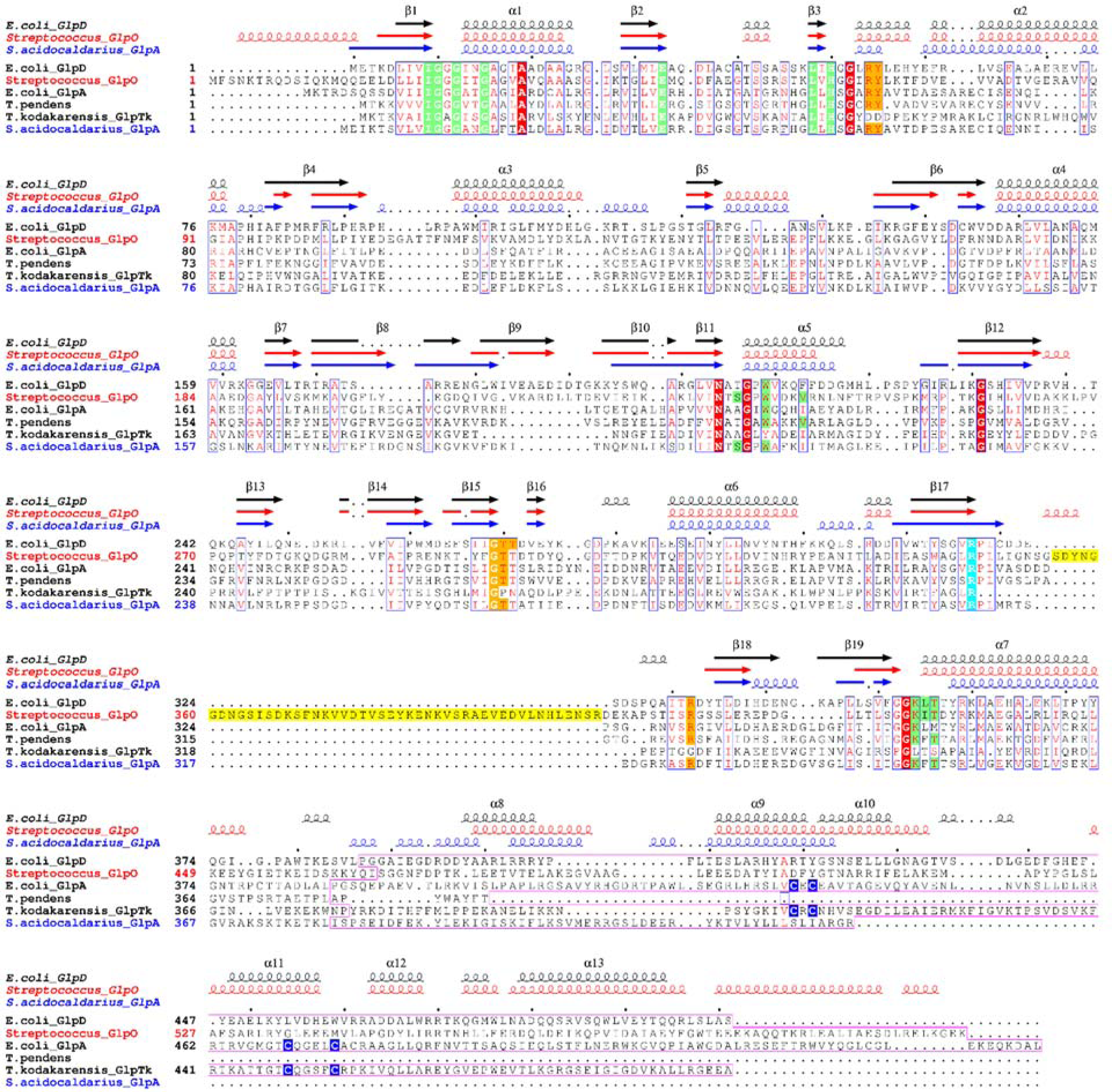
Sequence alignment of *S. acidocaldarius* G3PDH (Saci_2032) with *E. coli* GlpD (2qcu, P13035) and GlpA (P0A9C0), GlpO from *Streptococcus* sp. (2rgo, D0VWP7) as well as archaeal homologues from *T. pendens* (A1RZ95, Tpen_1127) and *T. kodakarensis* (Q5JGZ7, TK1393). The secondary structural elements from the available crystal structures of GlpD and GlpO as well as the predicted structure for *S. acidocaldarius* G3PDH by alphaFold based modelling are depicted above of the sequences. The alignment illustrates the sequence and structural conservation in the N-terminal 370-390 aa residues. The residues shown to be involved in FAD cofactor binding and substrate (G3P/DHAP) binding in the GlpD and GlpO sturctures as well as their conservation in the other sequences are shaded green and orange, respectively. The arginine residue supposed to act as general base in catalysis is highlighted in cyan. The sequence insertion distinctive for GlpOs is shaded yellow. The amino acid residues comprising the C-terminal domains are boxed in pink with the cysteine residues forming the FeS clusters in GlpA and the *Thermococcus* G3PDH marked by blue shading. The figure was generated using ESPript 3.0 (Robert and Gouet, 2014).

Notably, all the archaeal G3PDH homologues differ from GlpD, GlpO, and GlpA enzymes in their C-terminus: The *Thermofilum pendens* related homologues nearly completely lack a C-terminal extension and the DAAO fold is only extended by a very short α helix of 5-6 amino acids (if at all) (supplementary Figure 11a). In the *S. acidocaldarius*-like archaeal G3PDH homologues the C-terminal tail comprises roughly 50 amino acids which does not form any known or annotated domain comprising only three α helices (and shows also no similarities to GlpA, GlpO, or GlpD) (supplementary Figure 11b). The Thermococcales G3PDHs as indicated by model predictions seem to contain the bfd-fold (pfam04324) like in GlpA with four cysteine residues present (and align well with the GlpAs, Figure 9), although the overall fold of the C-terminal domain appears to be different and somewhat shorter (120 aa) (supplementary Figure 11c and d).

The structural differences in C-terminal domains of GlpDs, GlpOs and GlpAs seem to be linked to their different functions. In GlpD, the C-terminus was demonstrated to be involved in dimer formation whereas in non-membrane associated, cytoplasmic dimeric GlpOs that are reactive with oxygen (in contrast to glpDs) the C-terminus is not involved in subunit interaction (Colussi et al., 2008; Yeh et al., 2008). For GlpA - although no crystal structure is available - the presence of an FeS clusters in the C-terminus suggest that this protein moiety is involved in electron transfer to GlpB which might necessitate their physical interaction, which in turn is supported by the fact that the GlpAB forms the catalytically active unit. The ColabFold prediction of the Saci_G3PDH complex (based on the finding that Saci_2032 alone forms a soluble dimer) (see supplementary Figure 12) suggests that also in the Saci_G3PDH the (dissimilar) C-terminus might be involved in dimer formation, which, however, leads to different spatial orientation of the DAAO domains to each other than observed in GlpD. This different spatial orientation of the dimer together with CoxG interaction then likely enables optimal transfer of electrons to the quinone pool in the cytoplasmic membrane in *S. acidocaldarius*.

### Distribution, sequence similarities and phylogenetic affiliation of GK and G3PDH isoenzymes from *S. acidocaldarius*

Previous studies have already addressed the question of distribution and phylogenetic affiliation of glycerol degrading genes/enzymes in Archaea (Elkhal et al., 2015; Yokobori et al., 2016; Villanueva et al., 2017). Complete sets of glycerol degrading enzymes and thus the potential ability to utilize glycerol as sole carbon and energy source were mainly found in the euryarchaeal orders of Thermococcales, Halobacteriales, Archaeoglobales and Thermoplasmatales and in the crenarchaeal lineages of Thermoproteales, Thermofilales, Desulfurococcales, and Sulfolobales (Yokobori et al., 2016; Villanueva et al., 2017). However, the GK/G3PDH homologues are present only in some representatives (not in all or at least not in the majority) of these lineages and thus only patchily distributed in Archaea. It was also indicated that the evolutionary history of GK and G3PDH genes in Archaea is different with the GK acquired by horizontal gene transfer from bacteria whereas the G3PDH might have been already present in the universal common ancestor as well as the bacterial and archaeal common ancestors (Yokobori et al., 2016). This is also reflected by the higher overall sequence identity among the archaeal GKs (40-70% identity) compared to archaeal G3PDHs (20-50% identity) as well as by the fact that the closest homologues to GK and G3PDH from *S. acidocaldarius* were identified in different archaeal lineages, i.e. in Thermococcales (61-68% identity) and Thermoproteales (45-48% identity), respectively. The different evolutionary history of GKs and G3PDHs points towards the conserved function of GK in all the organisms whereas the G3PDH shows more functional variability with respect to interaction partners, membrane anchoring, and electron acceptors (see below).

Phylogenetic analyses of the G3PDH sequences from all domains resulted in similar tree topologies as previously reported (Rawls et al., 2011; Yokobori et al., 2016; Williams et al., 2017), however, closer inspection of structural and sequence features including gene neighborhood analysis provided deeper functional insights as illustrated in Figure 10.

**Figure 10:**
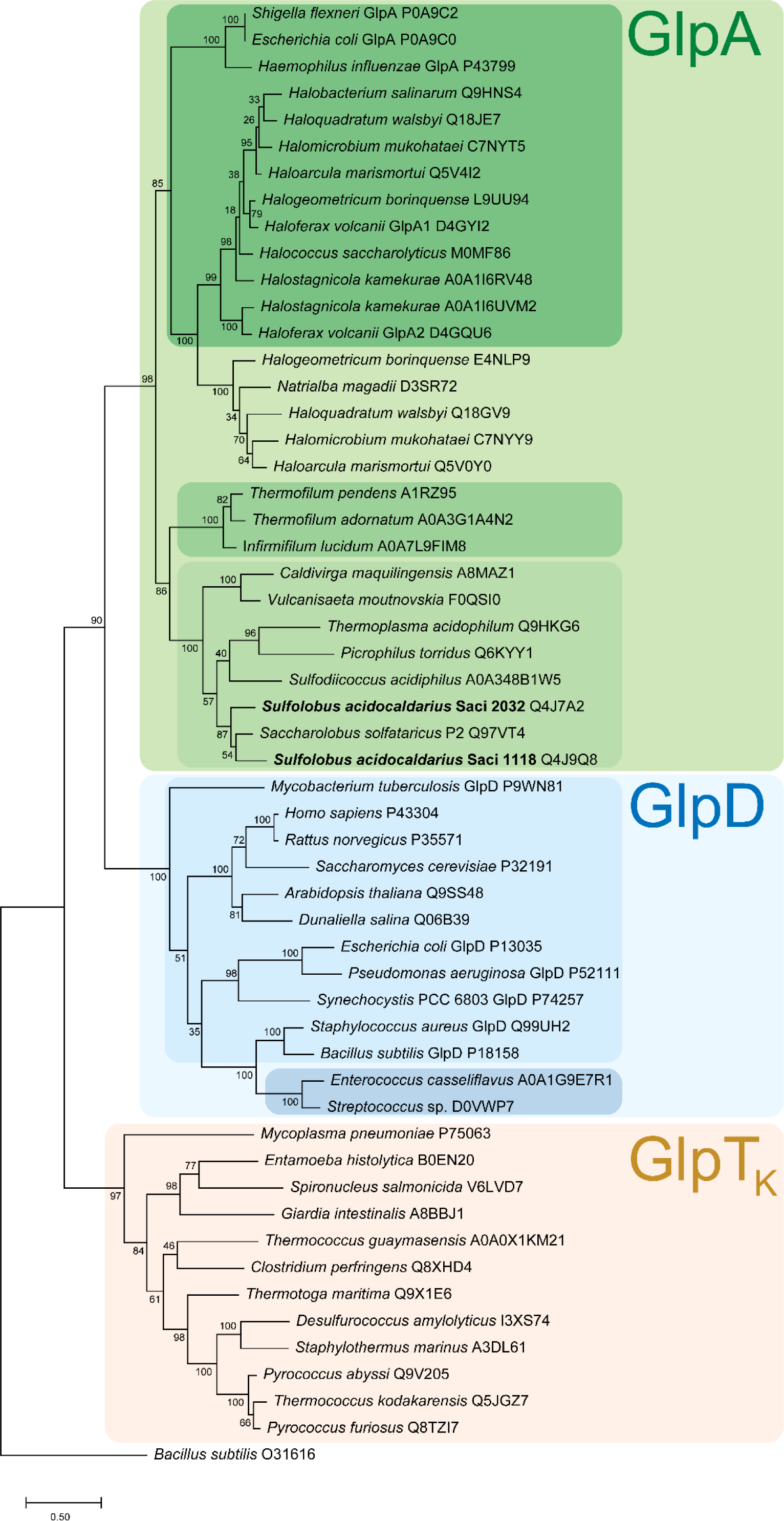
Phylogenetic affiliation of the *S. acidocaldarius* G3PDH isoenzymes Saci_2032 and Saci_1118 with selected GlpDs, GlpOs, and GlpAs from other Archaea, Bacteria and Eukarya. (organism names and uniprot accession numbers are given). GlpA homologous are shaded green, with the canonical bacterial and haloarchaeal GlpAs shown in dark green, the *Thermophilum* homologues in green and the *S. acidocaldarius* homologues in light green. The GlpD cluster shaded blue comprises the GlpDs and GlpOs highlighted in light and dark blue, respectively. The Thermococcus-like G3PDHs, designated as GlpTk are shown in orange. The evolutionary history was inferred by using the Maximum Likelihood method and JTT matrix-based model (Jones et al., 1992). The tree with the highest log likelihood (−22770.18) is shown. The percentage of trees in which the associated taxa clustered together is shown next to the branches. Initial tree(s) for the heuristic search were obtained automatically by applying Neighbor-Join and BioNJ algorithms to a matrix of pairwise distances estimated using a JTT model, and then selecting the topology with superior log likelihood value. The tree is drawn to scale, with branch lengths measured in the number of substitutions per site. This analysis involved 55 amino acid sequences. All positions containing gaps and missing data were eliminated (complete deletion option) resulting in a total of 280 positions in the final dataset. Evolutionary analyses were conducted in MEGA 11 (Tamura et al., 2021).

The FAD dependent G3PDHs cluster according to their different C-terminal extensions: GlpAs, GlpDs, and GlpOs form distinct subgroups within one main cluster within the DAAO superfamily with GlpDs and GlpOs more closely related to each other than to GlpAs. In contrast, the mycoplasma GlpOs appear only distantly related to glpDs/As/and Os (Elkhal et al., 2015). The G3PDH from Sulfolobales and Thermoplasmatales additionally including the *Caldivirga maquilingensis* and *Vulcanisaeta moutnovskia* homologues as well as those from Thermofilales form distinct subclusters separated from canonical bacterial and haloarchaeal sequences within the GlpA subgroup, justifying their annotation as GlpAs. Based on these analyses, no archaeal GlpDs could be identified. The Thermococcales sequences constitute a cluster clearly separated from *glpA/D* sequences and might thus represent a distinct family within the DAAO superfamily designated herein as GlpTk. This cluster also comprises sequences from anaerobic Crenarchaea (Desulfurococcales) as well as anaerobic Bacteria (e.g. *Thermotoga maritima* and *Clostridium perfringens*) and amitochondriate protists (*Giardia intestinalis*, *Entamoeba histolytica*, and *Spironucleus salmonicida*).

The phylogenetic affiliation does not only reflect the structural differences of FAD-dependent G3PDHs, it is also accompanied by a conserved gene organization (Figure 11): (i) The canonical *glpA* genes always cooccur in operons with downstream located *glpB* and *C* genes (Figure 11a). (ii) In the *Thermofilum* related species three genes are located downstream of the *glpA* gene homologue encoding putative counterparts of the succinate dehydrogenase b, c and d subunits (also involved in membrane anchoring and electron transfer (Unden et al., 2008)) (Figure 11b). (iii) All genes of the *S. acidocaldarius* related *glpA* subgroup show the *coxG* gene directly downstream of the *glpA* gene homologue (Figure 11c). (iv) The *glpD* and *glpO* genes don’t show any obvious operon like structures with other genes which coincides with their function as homodimers irrespective of their membrane or cytoplasmic localization, respectively (Figure 11d and e). (v) The Thermococcus related *glpTk* genes including the bacterial sequences from *T. maritima* and *C. perfringens* (see above) are all organized in a cluster with two genes downstream annotated to encode a pyridine nucleotide disulfide oxidoreductase/FAD-dependent oxidoreductase/ NADH oxidase (NOX) and a DUF1667 domain protein/molybdopterin oxidoreductase Fe_4_S_4_ domain (MOX) shown to from a soluble complex upon *in vitro* reconstitution (Koga et al., 2019) (Figure 11f). Interestingly, the GlpTk-like G3PDH enzymes from protists such as *Giardia intestinalis* (Lalle et al., 2015), *Entamoeba histolytica* and *Spironucleus salmonicida* represents a fusion protein with a GlpTk, NOX, and a DUF1667 domain. Thus, the observed clustering in the phylogenetic tree is accompanied by the formation of different complexes with proteins encoded downstream of the respective G3PDH genes in operon like structures for membrane association and electron transfer to the respective acceptors. Therefore, the acquisition of glycerol utilization necessitated the concomitant evolution of tailored membrane anchoring and/or electron transfer systems to fit the requirements of the organism’s lifestyle, metabolism and terminal electron acceptors.

**Figure 11:**
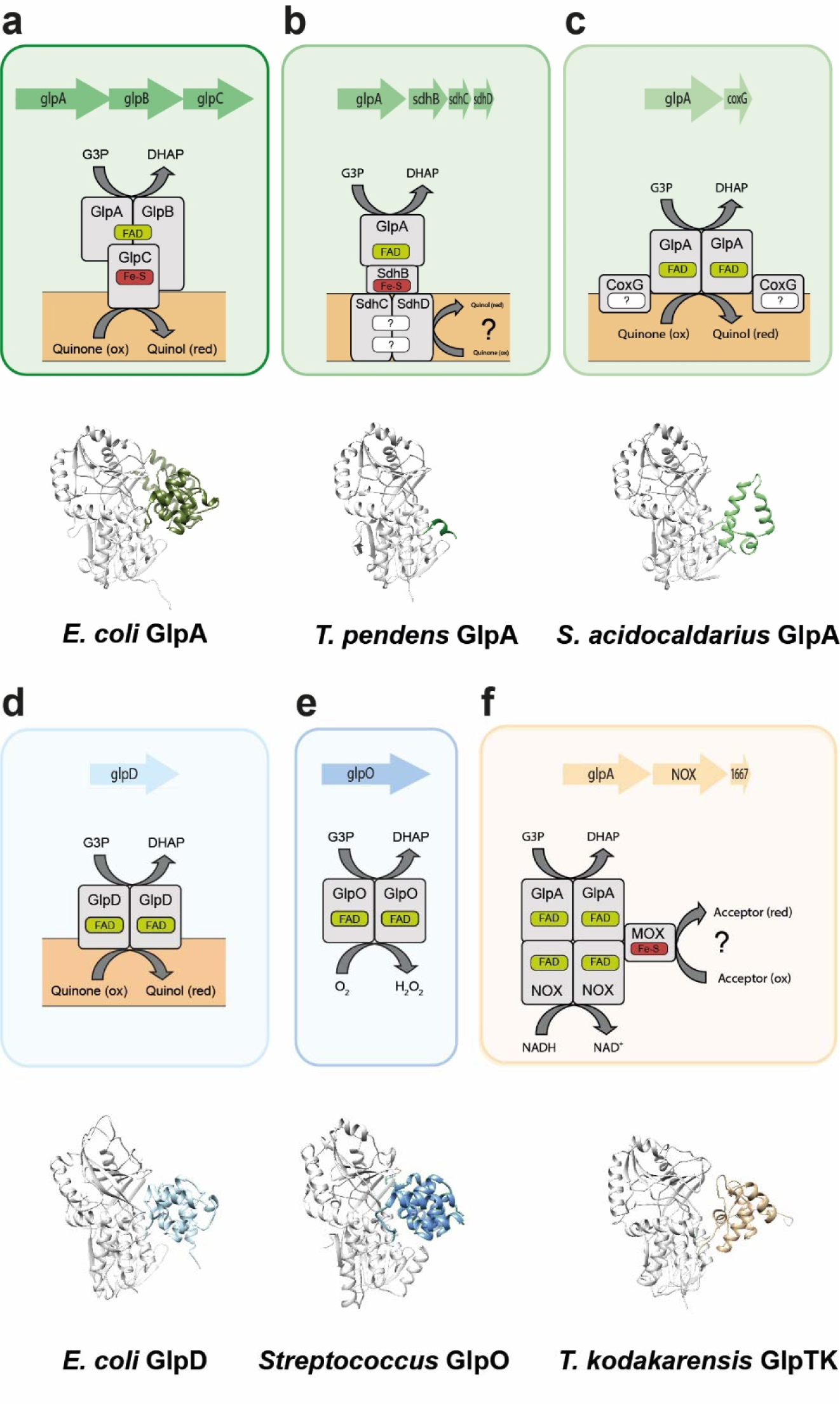
Comparison of different G3PDHs from Bacteria and Archaea. The respective genome neighbourhood, the derived reconstructed protein complexes with potential interaction partners, membrane or cytoplasmic localization and cofactor content, as well as the catalytic oxidoreductase-monomer 3D-structure of the G3PDHs GlpABC (*E. coli* (**a**), GlpA-SdhBCD (*T. pendens)* (**b**), GlpA-CoxG (*S. acidocaldarius)* (**c**), GlpD (*E. coli)* (**d**) GlpO (*Streptocoocus)* (**e**) and GlpTk-NOX-MOX (*T. kodakarensis)* (**f**) are shown. For colour code see Figure 9. For all 3D structures the principal FAD-binding G3P-oxidoreductase domain is coloured in grey while the highly divergent C-terminus is coloured according to Figure 9.

In summary, we herein unraveled the glycerol degradation in *S. acidocaldarius* to proceed via the GK-G3PDH pathway involving “classical” bacterial GlpK-like GKs from the FGGY family. However, with respect to the G3PDHs the pathway differs remarkably from those known from bacteria and haloarchaea. Although sequence comparison clearly identified the G3PDH as GlpA homologue, the enzyme shows an unusual subunit composition lacking the B and C subunits, a different domain structure in the C-terminus of the catalytic GlpA subunit, and a different mode of membrane anchoring via a CoxG homologue instead of a GlpC. The *S. acidocaldarius* genome harbors two paralogous copies of each GK and G3PDH. However, only one of these GK-G3PDH couples (Saci_2031-2033) is highly upregulated on glycerol and essential for glycerol degradation. Furthermore, although the physiological significance remains to be studied, sequence analyses suggest that glycerol metabolism in Archaea is more versatile with respect to the G3PDHs their interaction partners, membrane association, and electron transfer mechanisms.

