## Supplementary Tables and Figures for "Glycerol degradation in the thermoacidophilic crenarchaeon *Sulfolobus acidocaldarius* involves an unusual glycerol-3-phosphate dehydrogenase"

### Supplementary information

**Supplementary Table 1: Differentially expressed genes and proteins given as log2-fold-changes in *S. acidocaldarius* MW00G after growth on glycerol and 0.1 % /w/v) N-Z-Amine (NZA) compared to D-xylose identified via RNASeq (transcriptome) and label free quantification (LFQ) (proteome) analyses. For growth on NZA only log2-fold changes higher than 2 are given**

| | | Transcriptome (log2-fold changes) | | | | Proteome (log2-fold changes) | | | |
|---|---|---|---|---|---|---|---|---|---|
| | Glycerol concentration | 10 mM | 20 mM | 40 mM | NZA | 10 mM | 20 mM | 40 mM | NZA |
| **Locus tag** | **Function** | | | | | | | | |
| **Saci_0451** | ESCRT-III | 3.75 | 4.08 | 3.27 | 4.93 | 6.86 | 7.15 | 6.32 | 8.92 |
| **Saci_0942** | CopG family transcriptional regulator | 2.10 | 2.61 | 2.19 | 2.82 | 3.51 | 3.72 | 3.68 | 3.34 |
| **Saci_1050** | ATPase, ParA family | 2.96 | 3.00 | 2.85 | - | 2.61 | 2.56 | 2.71 | - |
| **Saci_1052** | GYD domain-containing secretory protein | 3.87 | 3.82 | 3.81 | - | 5.32 | 5.38 | 5.20 | - |
| **Saci_1054** | 3-(methylthio)propionyl-CoA ligase | 3.39 | 2.44 | 2.20 | - | 4.63 | 4.52 | 4.68 | - |
| **Saci_1058** | Xylulokinase | 3.12 | 2.99 | 2.94 | - | 2.62 | 2.58 | 2.71 | - |
| **Saci_1099** | Betaine-aldehyde dehydrogenase | 3.00 | 2.32 | 2.17 | - | 5.40 | 5.58 | 5.50 | - |
| **Saci_1228** | Nucleoside triphosphate hydrolase | 3.37 | 3.10 | 2.56 | 5.06 | 3.84 | 4.00 | 3.39 | 5.47 |
| **Saci_1372** | Vesicle-fusing ATPase | 3.04 | 2.25 | 2.00 | 4.76 | 4.05 | 3.83 | 4.02 | 6.80 |
| **Saci_1416** | ESCRT-III | 3.91 | 3.99 | 3.25 | 5.60 | 4.42 | 4.45 | 3.85 | 6.61 |
| **Saci_1616** | Antitoxin (DNA-binding domain) | 2.08 | 2.54 | 2.76 | - | 3.29 | 3.17 | 3.20 | 3.01 |
| **Saci_1762** | ABC-type | 3.33 | 3.11 | 3.24 | 3.03 | 3.17 | 3.05 | 2.98 | 2.42 |
| **Saci_1763** | ABC-type | 3.54 | 2.91 | 3.15 | - | 3.75 | 3.82 | 3.36 | 3.01 |
| **Saci_1764** | ABC-type | 3.53 | 3.12 | 3.49 | 3.41 | 2.87 | 2.94 | 2.89 | 2.26 |
| **Saci_1765** | ABC-type | 3.39 | 3.17 | 3.38 | 3.34 | 2.89 | 2.98 | 2.89 | 2.24 |
| **Saci_1804** | DUF2173 domain-containing protein | 2.39 | 2.19 | 2.74 | - | 4.85 | 4.94 | 5.04 | - |
| **Saci_1810** | Energy-coupling factor cobalt transporter, ATP-binding protein | 2.83 | 2.37 | 2.35 | - | 3.51 | 3.53 | 3.58 | - |
| **Saci_1855** | DUF973 domain-containing membrane protein | 7.66 | 6.92 | 6.98 | - | 5.81 | 5.90 | 5.88 | 3.13 |
| **Saci_1856** | DUF973 domain-containing membrane protein | 3.74 | 3.88 | 3.90 | - | 4.13 | 4.15 | 4.07 | - |
| **Saci_2032** | Glycerol 3-phosphate dehydrogenase | 3.92 | 3.31 | 2.97 | 2.90 | 3.01 | 2.98 | 3.10 | - |
| **Saci_2033** | Glycerol kinase | 3.96 | 3.51 | 3.19 | 2.37 | 2.47 | 2.47 | 2.24 | - |
| **Saci_2034** | Glycerol uptake facilitator | 3.85 | 3.52 | 3.20 | 2.22 | 4.84 | 3.59 | 4.66 | - |
| **Saci_2139** | CBS domain containing protein | 4.62 | 5.54 | 5.81 | 2.41 | 3.51 | 3.38 | 3.05 | - |
| **Saci_2188** | Ribonucleoside-diphosphate reductase, subunit beta | 3.42 | 2.99 | 3.07 | - | 2.93 | 2.61 | 2.36 | - |
| **Saci_2206** | Lipoate-protein ligase A | 4.77 | 4.16 | 4.13 | - | 3.68 | 3.69 | 3.59 | - |
| **Saci_2208** | 3-hydroxyacyl-CoA dehydrogenase | 5.30 | 4.72 | 4.56 | - | 9.34 | 9.38 | 9.45 | - |
| **Saci_2209** | Acetyl-CoA C-acetyltransferase | 4.26 | 3.63 | 3.45 | - | 2.75 | 2.76 | 2.54 | - |
| **Saci_2210** | DUF35 OB-fold domain containing protein | 5.15 | 4.12 | 4.16 | - | 6.93 | 6.76 | 7.02 | - |
| **Saci_2211** | 3-(methylthio)propionyl-CoA ligase | 2.79 | 2.58 | 2.54 | - | 3.00 | 2.98 | 2.87 | - |
| **Saci_2212** | Ribonucleoside-diphosphate reductase, subunit beta | 6.32 | 5.23 | 5.38 | - | 3.91 | 3.95 | 3.37 | - |
| **Saci_2230** | DUF35 OB-fold domain containing protein | 2.94 | 2.42 | 2.68 | 2.94 | 2.80 | 2.37 | 2.74 | - |

| | | | | | | | | | |
|---|---|---|---|---|---|---|---|---|---|
| **Saci_2231** | DUF35 OB-fold domain containing protein | 3.29 | 2.63 | 2.93 | 3.31 | 2.95 | 2.85 | 2.96 | - |
| **Saci_2232** | Acetyl-CoA C-acetyltransferase | 3.39 | 2.62 | 2.86 | 3.31 | 2.77 | 2.65 | 2.61 | - |
| **Saci_2233** | Acetyl-CoA C-acetyltransferase | 3.24 | 2.81 | 3.07 | 3.08 | 2.80 | 2.73 | 2.71 | - |
| **Saci_2235** | 3-hydroxypropionyl-CoA synthase | 5.67 | 5.07 | 5.30 | - | 8.19 | 8.39 | 7.99 | - |
| **Saci_2258** | Membrane protein, terminal oxidase function | 2.25 | 2.60 | 2.84 | - | 2.48 | 2.43 | 2.39 | - |
| **Saci_2259** | Cytochrome B558 subunit B | 2.61 | 2.71 | 3.02 | - | 4.61 | 4.10 | 4.54 | - |
| **Saci_2261** | Rieske Fe-S protein | 2.49 | 3.23 | 3.35 | - | 5.08 | 4.87 | 5.16 | - |
| **Saci_2263** | Quinol oxidase subunit 1/3 | 2.97 | 3.05 | 3.13 | - | 5.85 | 5.46 | 5.73 | - |
| **Locus tag** | **Function** | | | | | | | | |
| **Saci_0137** | Phosphomethylpyrimidine synthase | -4.55 | -4.23 | -4.57 | - | -2.86 | -2.93 | -2.80 | - |
| **Saci_0314** | Hypothetical protein | -5.43 | -4.33 | -4.15 | -4.49 | -2.36 | -3.06 | -2.10 | - |
| **Saci_0526** | Thiazole/oxazole-forming peptide maturase | -2.23 | -2.32 | -2.03 | - | -3.01 | -3.33 | -3.20 | -3.43 |
| **Saci_0531** | ABC-type Mn/Zn transport system | -3.54 | -2.59 | -2.54 | - | -3.76 | -3.33 | -3.39 | -2.89 |
| **Saci_1700** | 2,5-dioxovalerate dehydrogenase | -6.09 | -5.54 | -6.06 | -6.07 | -7.18 | -7.32 | -7.08 | -6.61 |
| **Saci_1707** | ABC-type molybdate transport system | -5.53 | -5.93 | -5.31 | -5.58 | -4.34 | -4.79 | -5.08 | -4.69 |
| **Saci_1830** | Alpha/beta hydrolase | -4.95 | -4.19 | -4.07 | -2.14 | -4.16 | -4.49 | -3.60 | -2.28 |
| **Saci_1831** | Transciptional regulator | -4.13 | -3.81 | -3.81 | - | -2.34 | -2.04 | -2.21 | - |
| **Saci_1938** | 2,5-dioxovalerate dehydrogenase | -4.08 | -4.01 | -4.17 | -3.63 | -6.83 | -6.95 | -6.73 | -6.70 |
| **Saci_1939** | 2-dehydro-3-deoxy-D-arabinonate dehydratase | -3.89 | -3.44 | -3.77 | -3.65 | -5.70 | -5.50 | -5.48 | -5.43 |
| **Saci_2090** | Rieske Fe-S protein | -3.12 | -2.86 | -2.62 | - | -3.33 | -3.93 | -3.91 | - |
| **Saci_2122** | Xylose/arabinose ABC-type transport system | -2.35 | -2.43 | -2.41 | -3.49 | -4.07 | -4.10 | -4.24 | -6.56 |
| **Saci_2203** | Sulfate adenylyltransferase | -3.61 | -4.16 | -4.27 | - | -2.16 | -2.49 | -2.75 | -8.21 |
| **Saci_2204** | Hypothetical protein | -3.73 | -4.08 | -4.25 | -7.77 | -2.89 | -3.60 | -3.46 | -6.39 |

**Supplementary Table 2: Enriched proteins via co-immunoprecipitation anti-HA antibodies coupled to magnetic beads using *S. acidocaldarius* MW00G expressing HA-*saci_2031* or HA-*saci_1119* given as log2-fold-changes.**

| majority protein ID (HA-Saci_2031) | gene name ID | log2-fold changes |
|---|---|---|
| **Q4J7A4** | *saci_2031* | 11.06 |
| **Q4J7Q9** | *saci_1865* | 6.00 |
| **Q4JBM7** | *saci_0386* | 4.19 |
| **Q4JB23** | *saci_0614* | 2.91 |
| **Q4JAW2** | *saci_0687* | 2.83 |
| **Q4JAP0** | *saci_0766* | 2.57 |
| **Q4J7A2** | *saci_2032* | 2.43 |
| **P13123** | *saci_0064* | 2.42 |
| **Q4J6G8** | *saci_2329* | 2.15 |
| **Q4J9Q8** | *saci_1118* | 2.13 |
| **majority protein ID (HA-Saci_1119)** | **gene name ID** | **log2-fold changes** |
| **Q4J9R0** | *saci_1119* | 14.72 |
| **Q4J9A5** | *saci_1283* | 6.76 |
| **Q4JBM7** | *saci_0386* | 3.71 |
| **Q4J9Q8** | *saci_1118* | 3.45 |
| **Q4J953** | *saci_1337* | 3.37 |
| **Q4JB63** | *saci_0574* | 2.79 |
| **Q4JAK6** | *saci_0803* | 2.21 |

**Supplementary Table 3:** Strains and Plasmids used in this study

| Designation | Genotype or description | Reference/Source |
|---|---|---|
| **Strains** | | |
| MW001 | *Sulfolobus acidocaldarius* strain; For growth or deletion mutant construction | (Wagner et al., 2012) |
| DH5α | *Escherichia coli* strain; Plasmid construction | Hanahan, USA |
| Rosetta (DE3) | *Escherichia coli* strain; Gene expression; pRARE (chloramphenicol resistance) | Stratagene, USA |
| MW00G | *Sulfolobus acidocaldarius* strain; glycerol adapted strain | This study |
| MW1257 | *S. acidocaldarius* MW00G Δ*saci_1117* | This study |
| MW1258 | *S. acidocaldarius* MW00G Δ*saci_2033* | This study |
| MW1259 | *S. acidocaldarius* MW00G Δ*saci_1117* Δ*saci_2033* | This study |
| MW1257 Δ*saci_1494:P*$_{saci\_1117}$ *saci_1117 CtSS* | *S. acidocaldarius* MW00G Δ*saci_1117* Δ*saci_1494:P*$_{saci\_1117}$ *saci_1117 CtSS* | This study |
| MW1258 Δ*saci_1494:P*$_{saci\_2033}$ *saci_2033-2034* | *S. acidocaldarius* MW00G Δ*saci_2033* Δ*saci_1494:P*$_{saci\_2033}$ *saci_2033-2034* | This study |
| **Plasmids** | | |
| pET15b | *E. coli* expression plasmid carrying an N-terminal His tag (ampicillin resistance) | Novagen, USA |
| pETDuet-1 | *E. coli* coexpression plasmid carrying an N-terminal His tag (multiple cloning site 1) or S-tag (multiple cloning site 2) (ampicillin resistance) | Novagen, USA |
| pBS-Ara-albaUTR-FX | Expression plasmid for *S. acidocaldarius* containing an arabinose inducible promotor and C-terminal strep tag (ampicillin resistance) | unpublished |
| pSVAmalFX-SH10 | Expression plasmid for *S. acidocaldarius* containing a maltose inducible promotor and C-terminal 10xHis tag (ampicillin resistance) | (Wagner et al., 2014) |
| pSVAmalFX-*saci_1117*-SH10 | Expression of *saci_1117* in *S. acidocaldarius* | This study |
| pET15b-*saci_1118* | Expression of *saci_1118* in *E. coli* | This study |
| pETduet1-saci_*1118* | Expression of *saci_1118* in *E. coli* | This study |
| pETduet1-*saci_1118-saci_1119* | Coexpression of *saci_1118, saci_1119* in *E. coli* | This study |
| pET15b-saci_*2032* | Expression of *saci_2032* in *E. coli* | This study |
| pETduet1-saci_*2032* | Expression of *saci_2032* in *E. coli* | This study |
| pETduet1-*saci_2032-saci_2031* | Coexpression of *saci_2031, saci_2032* in *E. coli* | This study |
| pBS-Ara-albaUTR-FX-*saci_2033* | Expression of *saci_2033* in *S. acidocaldarius* | This study |
| pSVA13204 | Expression of *saci_2031* in *S. acidocaldarius,* based on pSVAaraFX-HA | This study, (Van der Kolk et al., 2020) |
| pSVA407 | Gene targeting plasmid for knockout generation | (Wagner et al., 2012) |
| KO plasmid 1117 | Gene knock-out of *saci_1117* | This study |
| KO plasmid 2033 | Gene knock-out of *saci_2033* | This study |
| pSVA407-Δ*saci_1494:P*$_{saci\_1117}$ *saci_1117 CtSS* | In trans complementation of MW1257, insertion of *saci_1117* with the native promoter into *saci_1494* | This study |
| pSVA407-Δ*saci_1494:P*$_{saci\_2033}$ *saci_2033-2034* | In trans complementation of MW1258, insertion of the *saci_2033-2034* operon with the native promoter into *saci_1494* | This study |

**Supplementary Table 4:** Sequences of oligonucleotide used in this study

| ORFs | Name | Sequences (5´→3´) (restriction sites are marked in red) |
|---|---|---|
| **Oligonucleotides for cloning** | | |
| *Saci_1117* | *Fw (pSVAmalFX-SH10)-NcoI* | CAAGATCCATGGTGTCCAAATACATATTGGC |
| | *Rev (pSVAmalFX-SH10)-XhoI* | CTGCTTCTCGAGTTCCATAGTCTCCACTTC |
| | *Fw (pSVA407) upstream* | CCCGGATCCGCTTACCCTCTAGGCTTTCC |
| | *Rev (pSVA407) upstream* | CAAATACATAATGGAATAGAGAAATGCCAAAAG |
| | *Fw (pSVA407) downstream* | TCTATTCCATTATGTATTTGGACACATTAACTATTTT |
| | *Rev (pSVA407) downstream* | AGCCGGGCCCCTCAAGACCCTCTTCACTTC |
| | *Fw (pSVA407comp) GA* | GCAGAGTATTTTGTAGGTCCAGTGACCTCTTTCTACTAAA GTAACG |
| | *Rev (pSVA407comp) GA* | CTGTCTCGAATTCCGTTCTCCTTACTTTTCGAACTGTGGG TGACTC |
| *Saci_1118* | *Fw (pET15b)-NdeI* | GTCCGACATATGATGAAGCAAAACAGTTCAG |
| | *Rev (pET15b)-BamHI* | AATTACGGATCCTTATTCCCCCCTCGCCAG |
| | *Fw (pETDuet-1)-BamHI* | GAGGAGGGATCCATGAAGCAAAACAGTTCAG |
| | *Rev (pETDuet-1)-HindIII* | GAGGAGAAGCTTTTATTCCCCCCTCGCCAG |
| *Saci_1119* | *Fw (pETDuet-1)-NdeI* | GAGGAGCATATGATGGAGATGAGTGGGAAG |
| | *Rev (pETDuet1)-XhoI* | GAGGAGCTCGAGTTAAACTCTTGAAAGACAGGAGA |
| | *Fw (pSVaraFX-HA)-Nco*I | CATCCATGGAGATGAGTGGGAAGTT |
| | *Rev (pSVaraFX-HA)-Xho*I | CATCTCGAGAACTCTTGAAAGACAGGAGA |
| *Saci_2031* | *Fw (pETDuet-1)-NdeI* | GAGGAGCATATGATGGAGATCAGCGGAGAG |
| | *Rev (pETDuet1)-XhoI* | GAGGAGCTCGAGCTAGGTAACTCTAGAAC |
| | *Fw (pSVaraFX-HA)-Nco*I | CATCCATGGAGATCAGCGGAGAGTTTG |
| | *Rev (pSVaraFX-HA)-Xho*I | CATCTCGAGGGTAACTCTAGAACGTATACAG |
| *Saci_2032* | *Fw (pET15b)-NdeI* | GTCCGACATATGATGGAAATAAAAACAAGCG |
| | *Rev (pET15b)-BamHI* | AATTACGGATCCTCATCTCCCTCTTGCAATTAATG |
| | *Fw (pETDuet-1)-BamHI* | GAGAGGGATCCATGGAAATAAAAACAAGCG |
| | *Rev (pETDuet-1)-HindIII* | GAGAGAAGCTTTCATCTCCCTCTTGCAATTAATG |
| *Saci_2033* | *Fw (pBS-Ara-albaUTR-FX)-NcoI* | GAACCATGGTGGCTGAAAAATACGTGATAG |
| | *Rev (pBS-Ara-albaUTR-FX)-XhoI* | GGTCTCGAGACCCCCAATAGTCTTAGC |
| | *Fw (pSVA407) upstream* | GCATCTCGAGCCTGTATCTCTTATAGCGTG |
| | *Rev (pSVA407) upstream* | TACAACTTCCGATGCCATAGCATTTTATCCATTTCTATTT |
| | *Fw (pSVA407) downstream* | ATAGAAATGGATAAAATGCTATGGCATCGGAAGTTGTAAG |
| | *Rev (pSVA407) downstream* | GCATGGGCCCTGATTAACTTCAGAAAACAGTG |
| | *Fw (pSVA407comp) GA* | GCAGAGTATTTTGTAGGTCCAGTGAGCTATGTTTATTGTT TCCACTTTTC |
| | *Rev (pSVA407comp) GA* | TTCACTGTCTCGAATTCCGTTCTCCCTTTAGTGGAGACAC CTGTTGGAG |
| *Saci_1494* up | *Fw (pSVA407comp) GA* | ACGGCCAGTGAATTGTAATACGACTCACTATAGGGCGAAT TGGGCCGTGTATAATGATGACCTATTTAGCTG |
| | *Rev (pSVA407comp1117) GA* | GTTACTTTAGTAGAAAGAGGTCACTGGACCTACAAAATAC TCTG |
| | *Rev (pSVA407comp2033) GA* | GAAAAGTGGAAACAATAAACATAGCTCACTGGACCTACAA AATACTCTG |
| *Saci_1494* down | *Fw (pSVA407comp1117) GA* | GTCACCCACAGTTCGAAAAGTAAGGAGAACGGAATTCGA GACAGTG |
| | *Fw (pSVA407comp2033) GA* | ACTCCAACAGGTGTCTCCACTAAAGGGAGAACGGAATTC GAGACAGTGA |

| | |
|---|---|
| *Rev (pSVA407comp) GA* | AGGCGGCCGCGAATTCACTAGTGATATCGAATTCCCGCC CTCTGATAATTCAGTGGCTTTATTC |

*Red underlined indicates restriction sites.*

**Supplementary Table 5: Kinetic parameters, oligomeric state, and cosubstrate spectrum of different characterized GKs from selected Bacteria and Archaea.**

| Enzyme | Organism | Cofactors | Oligomeric state | Substrate | $V_{max}$ [U/mg] | $K_M$ (for glycerol) [mM] | $K_M$ (ATP) [mM] | Literature |
|---|---|---|---|---|---|---|---|---|
| **Saci_1117** | *S. acidocaldarius* | $Mg^{2+}$ | homodimer | Glycerol, ATP | 170 (at 75°C) | 0.023 | 0.179 | This work |
| **Saci_2033** | *S. acidocaldarius* | $Mg^{2+}$ | homodimer | Glycerol, ATP | 339.7 (at 75°C) | 0.024 | 0.17 | This work |
| **GlpK** | *E. coli* | $Mg^{2+}$ | homodimer/homotetramer | Glycerol, ATP | 18.1-100 (at 25°C) | 0.0013 | 0.0078 | (Hayashi and Lin, 1967; Applebee et al., 2011) |
| **GlpK** | *Thermus thermophilus* | $Mg^{2+}$ | homotetramer | Glycerol, ATP | 56.7 (at 37°C) | 0.038 | 0.162 | (HUA-SHAN HUANG, 1997) |
| **GlpK** | *Trypanosoma brucei* | $Mg^{2+}$ | Homodimer | Glycerol, ATP | 226 (at 25°C) | 0.44 | 0.24 | (Králová et al., 2000; Balogun et al., 2014) |
| **GlpK** | *Thermococcus kodakarensis* | $Co^{2+}$ | homodimer/homohexamer | Glycerol, ATP | 1500 (at 80°C) | 0.111 | 0.059 | (Yuichi Koga et al., 1998; Hokao et al., 2020) |

**Supplementary Table 6: Kinetic parameters, oligomeric state, and cosubstrate spectrum of different characterized G3PDHs from selected organisms from all three domains of life.** DCPIP: 2,6-Dichlorphenolindophenol; FAD: flavin adenine dinucleotide; G3P: Glycerol-3-phosphate; MTT: (4,5-Dimethylthiazol-2-yl)-2,5-diphenyltetrazoliumbromid; PMS: Phenazine methosulfate; UQ: Ubiquinone.

| Enzyme | Organism | Cofactors per monomer | Oligomeric state | Substrate | $V_{max}$ [U/mg] | $K_M$ (for G3P) [mM] | $K_M$ (electron acceptor) [mM] | Literature |
|---|---|---|---|---|---|---|---|---|
| **Saci_1118** | *S. acidocaldarius* | 1 FAD | homodimer | G3P | 19.7 (DCPIP) | 0.019 (DCPIP) | 0.086 (UQ -1) | This work |
| **Saci_2032** | *S. acidocaldarius* | 1 FAD | homodimer | G3P | 44.5 (DCPIP) | 0.055 (DCPIP) | 0.179 (UQ -1) | This work |
| **GlpAB** | *E. coli* | 1 FAD<br>2 non-heme iron | heterodimer | G3P | 34.4 (PMS-MTT) | 0.339 (PMS-MTT) | | (Schryvers and Weiner, 1981) |
| **GlpA** | *Thermococcus kodakarensis* KOD1 | FAD | monomer | G3P | 5.63 (MTT) | 2.68 (MTT) | | (Koga et al., 2019) |
| **GlpD** | *E. coli* | 2 FAD | homodimer | G3P | 5.8 (DCPIP) | 0.8 (DCPIP) | | (Weiner and Heppel, 1972; Schryvers et al., 1978; Yeh et al., 2008a) |
| **GlpD** | Pig brain mitochondria | FAD | homodimer | G3P | 2.63 (UQ-1<br>0.8 - 2.9 (DCPIP) | 6.2 (U-Q10)<br>10 (DCPIP)<br>10 (UQ-0/1/2/6) | 0.013 (UQ-1)<br>0.076 (DCPIP) | (Dawson and Thorne, 1969; Cottingham and Ragan, 1980) |
| **GlpD** | *Vibrio alginolyticus* | FAD | unknown | G3P | 39.2 (PMS)<br>85.8 (ferricyanide) | 4.02 (PMS)<br>3.7 (ferricyanide) | 0.13 (PMS)<br>0.55 (ferricyanide) | (Unemoto et al., 1981) |
| **GlpD** | *Acidiphilium* sp. 63 | 0.58 FAD | homodimer | G3P | 32 (PMS-MTT) | - | - | (Hatta et al., 1989) |

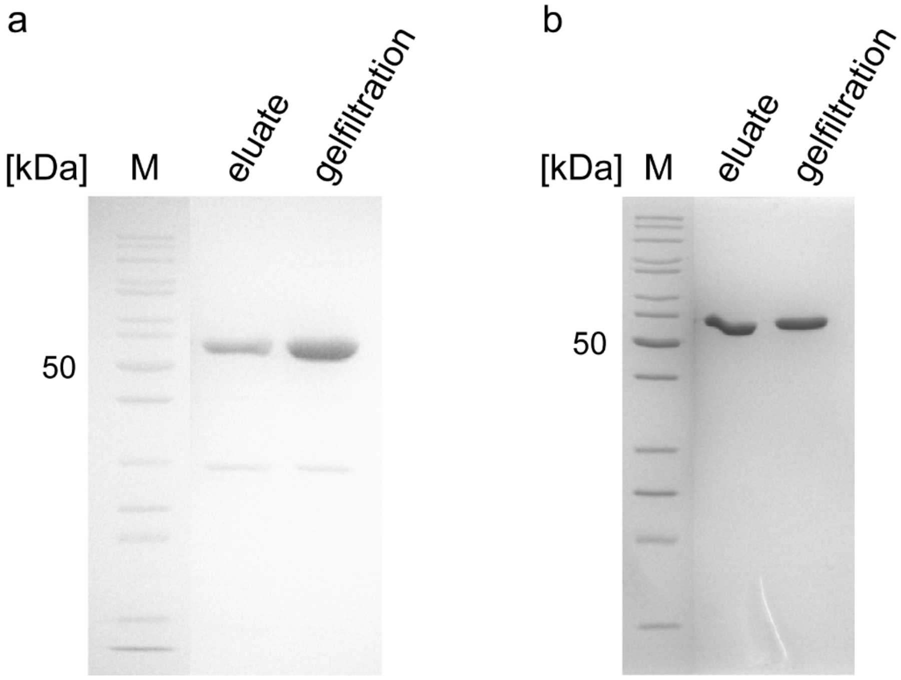


**Supplementary Fig. 1: Purification of recombinant glycerol kinase isoenzymes Saci_1117 and Saci_2033.** The glycerol kinases (GK) Saci_1117 (**a**) and Saci_2033 (**b**) were homologously produced in *S. acidocaldarius* MW001 using the pSVAmalFX-SH10 and pBS-Ara-albaUTR-FX expression vector, respectively. Proteins were purified via His-tag affinity and Strep-tag affinity chromatography, respectively, and size exclusion chromatography. Proteins (2 µg) were separated via SDS-PAGE and stained via Coomassie Blue. Marker (M): prestained PageRuler™ (Thermo Fischer scientific, USA).

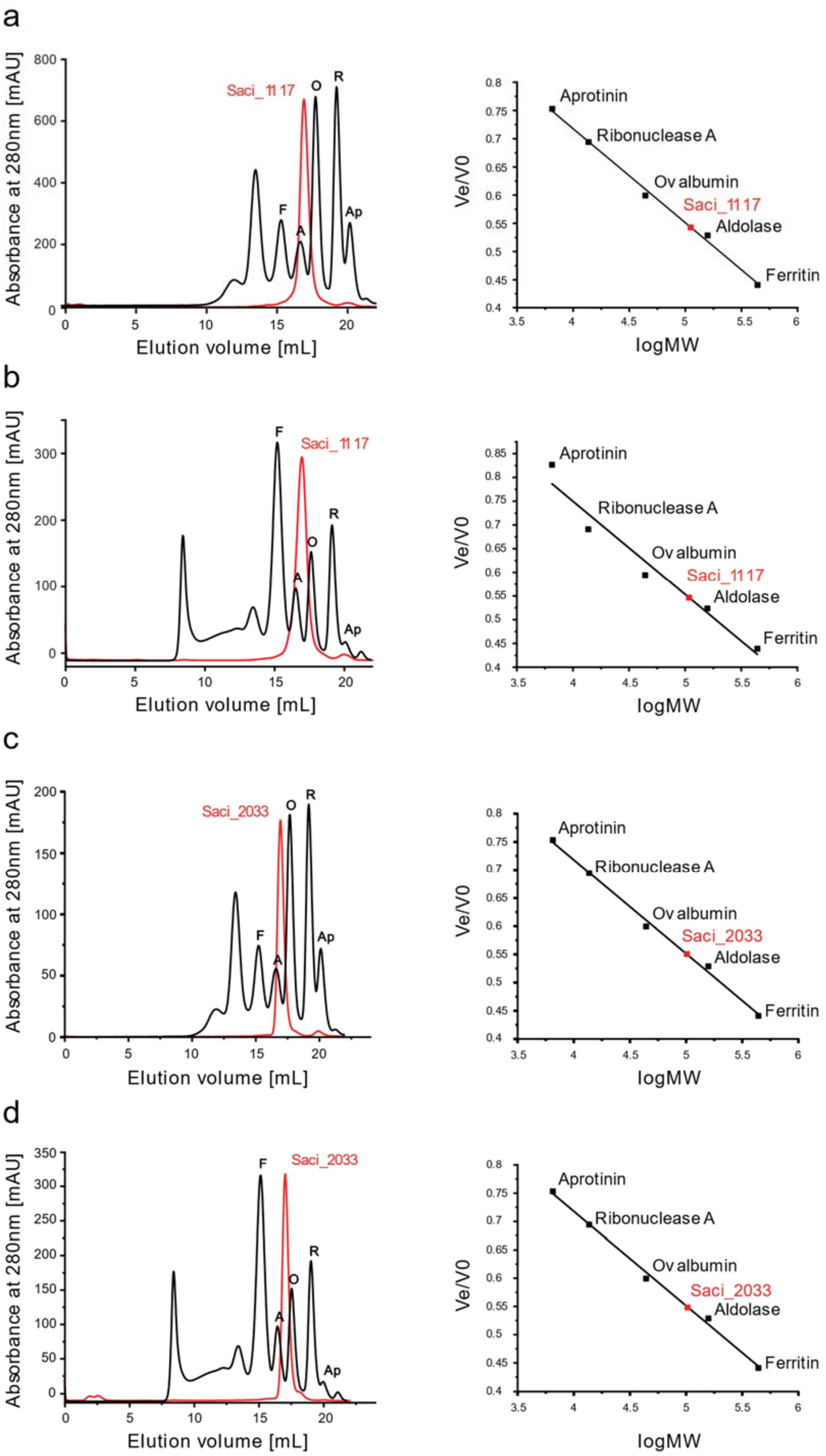
a
b
c
d
Absorbance at 280nm [mAU]
Elution volume [mL]
Saci_1117
Saci_2033
F
A
O
R
Ap
Ve/V0
logMW
Aprotinin
Ribonuclease A
Ovalbumin
Aldolase
Ferritin

**Supplementary Fig. 2: Size exclusion chromatography of the recombinant GK isoenzymes Saci_1117 and Saci_2033 from *S. acidocaldarius.*** Recombinant proteins were separated on a Superose 6 10/300 column in both, the absence (**a**, Saci_1117 and **c**, Saci_2033) and presence of 10 mM glycerol (**b**, Saci_1117 and **d**, Saci_2033). In the elution profiles (left panels) the GKs Saci_1117 and Saci_2033 from *S. acidocaldarius* are shown in red and the standard proteins for calibration in black. In the right panels the calibration curves are shown.

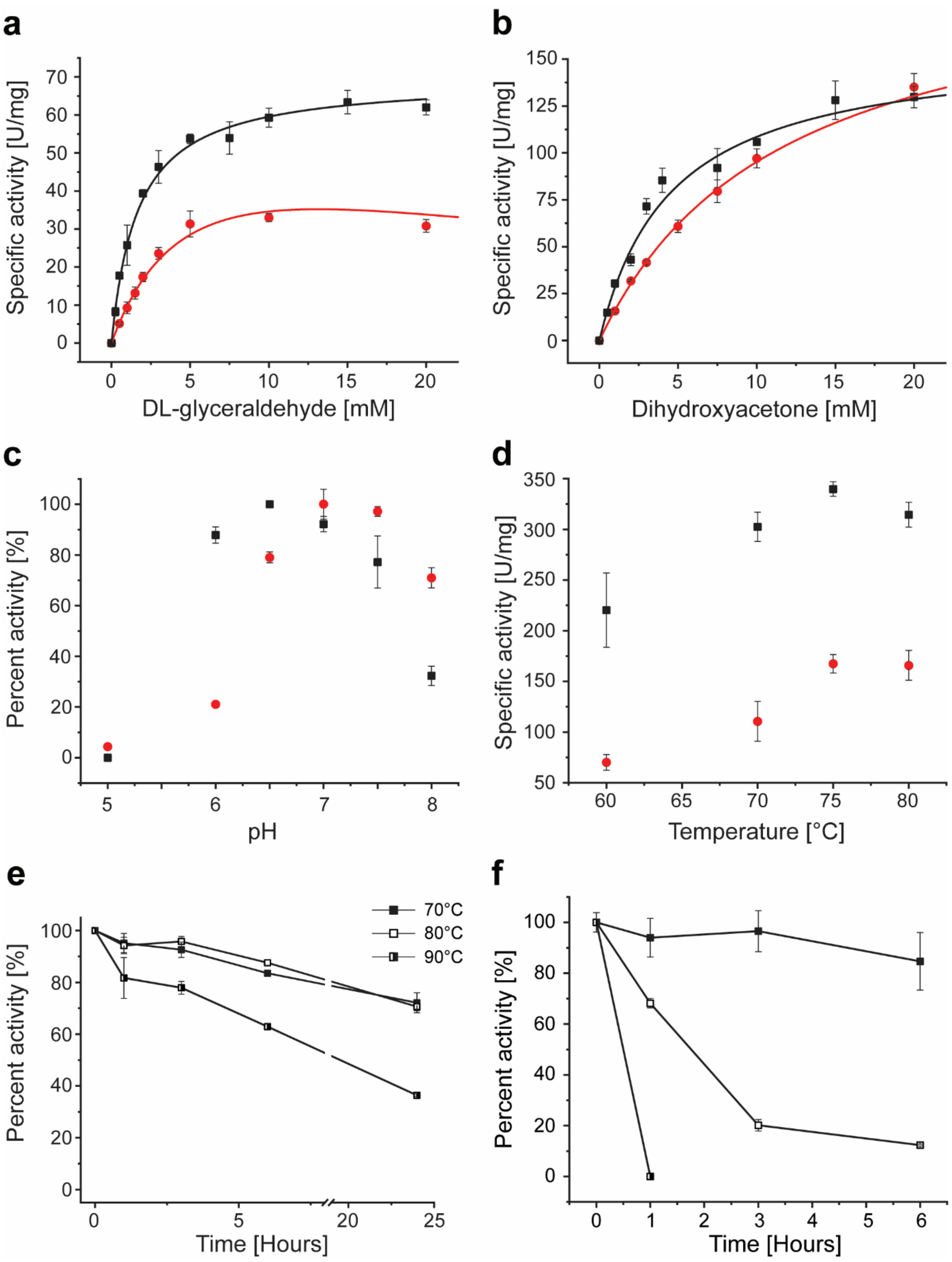


**Supplementary Fig. 3: Biochemical characterization of recombinant GK Saci_1117 and Saci_2033.** The kinetic properties of Saci_1117 (red circle) and Saci_2033 (black square) with (**a**) DL-glyceraldehyde and dihydroxyacetone (**b**) as substrate were determined in the coupled assay via PK and LDH at 50°C in 0.1 M TRIS-HCl pH 7 for Saci_1117 and 0.1 M MOPS pH 6.5 for Saci_2033, respectively. (**c**) The pH dependence of the Saci_1117 (red circle) and Saci_2033 (black square) activity was determined in the range of pH 5.0-8.0 using a mixed buffer system (0.05 M MES, 0.05 M HEPES, 0.05 M TRIS) at 50°C. Activity was determined in a continuous assay coupling the formation of ADP from ATP to the oxidation of NADH via pyruvate kinase (PK) and lactate dehydrogenase (LDH) following the decrease in absorbance at 340 nm. (**d**) The optimum temperature of Saci_1117 (red circle) and Saci_2033 (black square) was determined between 60°C to 80°C using a continuous assay with G3PDH (Saci_2032) as auxiliary enzyme and 0.1 mM DCPIP as electron acceptor. The thermal stability of the GK Saci_1117 (**e**) and Saci_2033 (**f**) was determined by monitoring the residual activity upon incubation at different temperatures (70, 80 and 90°C) over time (1, 3, 6, 24h). Saci_1117 was incubated in 100 mM TRIS-HCl, pH 7 (temperature adjusted) and Saci_2033 in 100 mM MOPS-KOH, pH 6.5 (temperature adjusted) at a protein concentration of 0.05 mg $ml^{-1}$. After the respective incubation period the residual activity was determined in the coupled assay via PK and LDH at 50°C. Experiments were performed in triplicate and error bars indicate the SD of the mean.

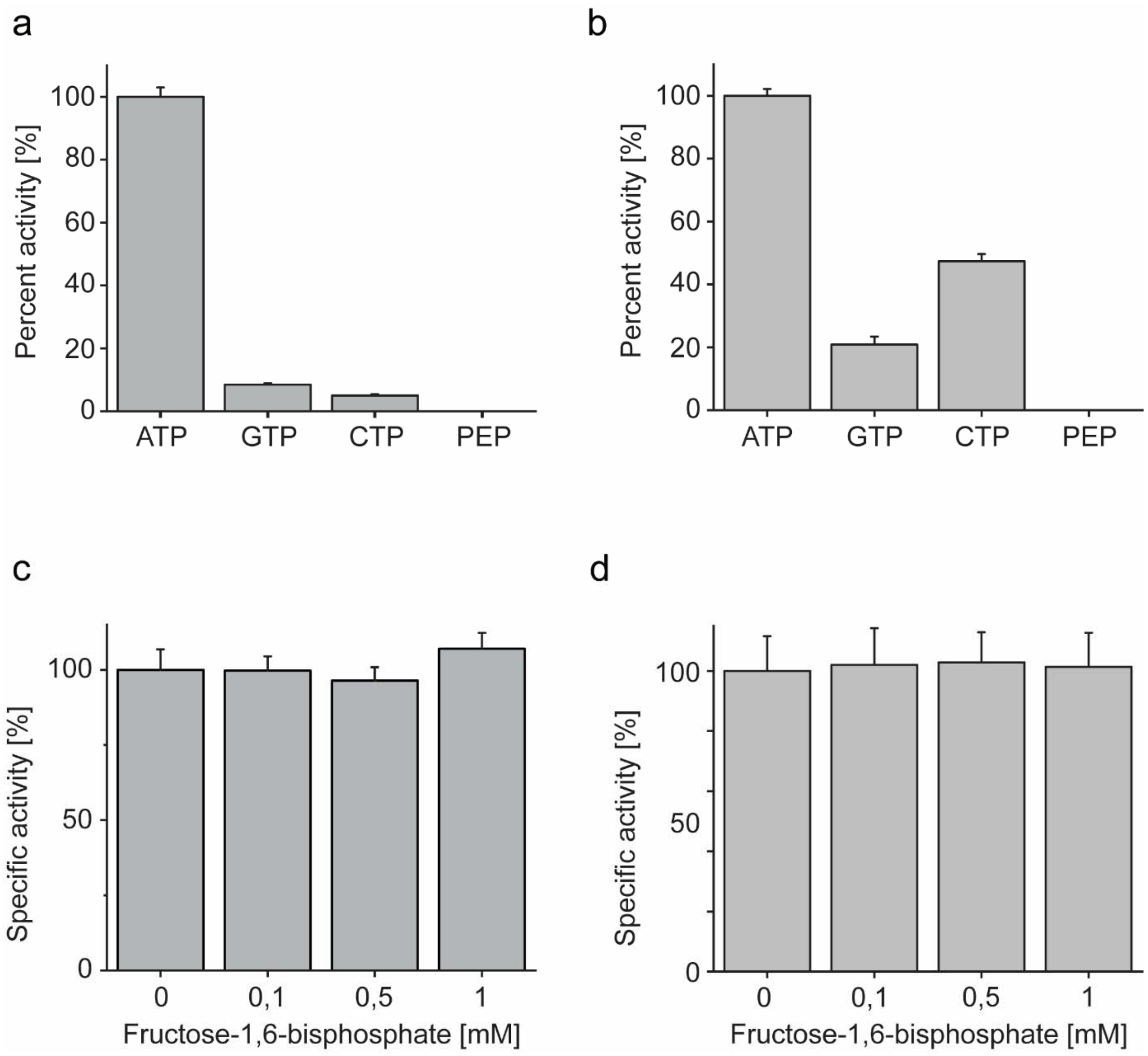


**Supplementary Fig. 4: Phosphate donor specificity and effect of fructose-1,6-bisphosphate on enzyme activity of the recombinant GK isoenzymes Saci_1117 and Saci_2033**. The enzyme activity of Saci_1117 (**a**) and Saci_2033 (**b**) with 2 mM glycerol as substrate and 5 mM of different phosphate donors was determined in a continuous assay at 75°C using G3PDH Saci_2032 as auxiliary enzyme. G3PDH couples the phosphorylation of glycerol by GK to the oxidation of G3P to DHAP by following the decrease in absorbance at 600 nm due to the reduction of DCPIP. The effect of fructose 1,6-bisphosphate on enzyme activity of Saci_1117 (**c**) and Saci_2033 (**d**) was determined in presence of increasing fructose 1,6-bisphosphate concentrations using the continuous coupled LDH/PK assay by following NADH oxidation as decrease in absorbance at 340 nm and 50°C. Experiments were performed in triplicate and error bars indicate the SD of the mean.

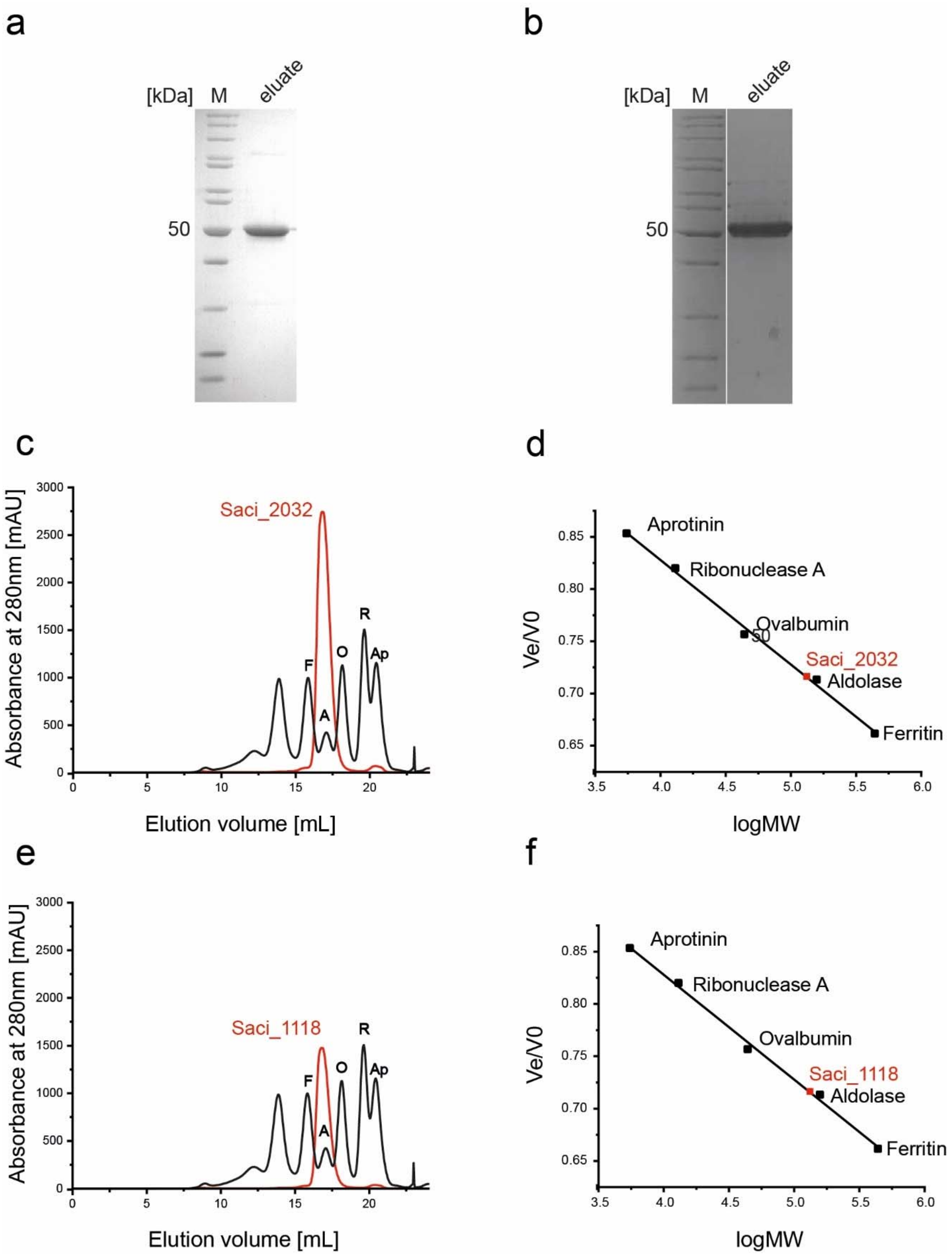


**Supplementary Fig. 5: SDS-Page and Size exclusion chromatography of the recombinant G3PDH Saci_1118 and Saci_2032 from *S. acidocaldarius* and the corresponding column calibration.** The glycerol-3-phospahte dehydrogenases (G3PDH) Saci_2032 **(a)** and Saci_1118 **(b)** were heterologously produced in *E. coli* Rosetta using pET15b and purified via His-tag affinity chromatography. Proteins (2 µg) were separated via SDS-PAGE and stained via Coomassie Blue. Marker (M): prestained PageRuler™ (Thermo Fischer scientific, USA). Recombinant proteins were separated on a Superose 6 10/300 column (**c,d**, Saci_2032 and **e,f**, Saci_1118). In the elution profiles (left panels) the G3PDHs

Saci_1118 and Saci_2032 from *S. acidocaldarius* are shown in red and the standard proteins for calibration in black. In the right panels the calibration curves are shown. and the standard proteins for calibration in black.

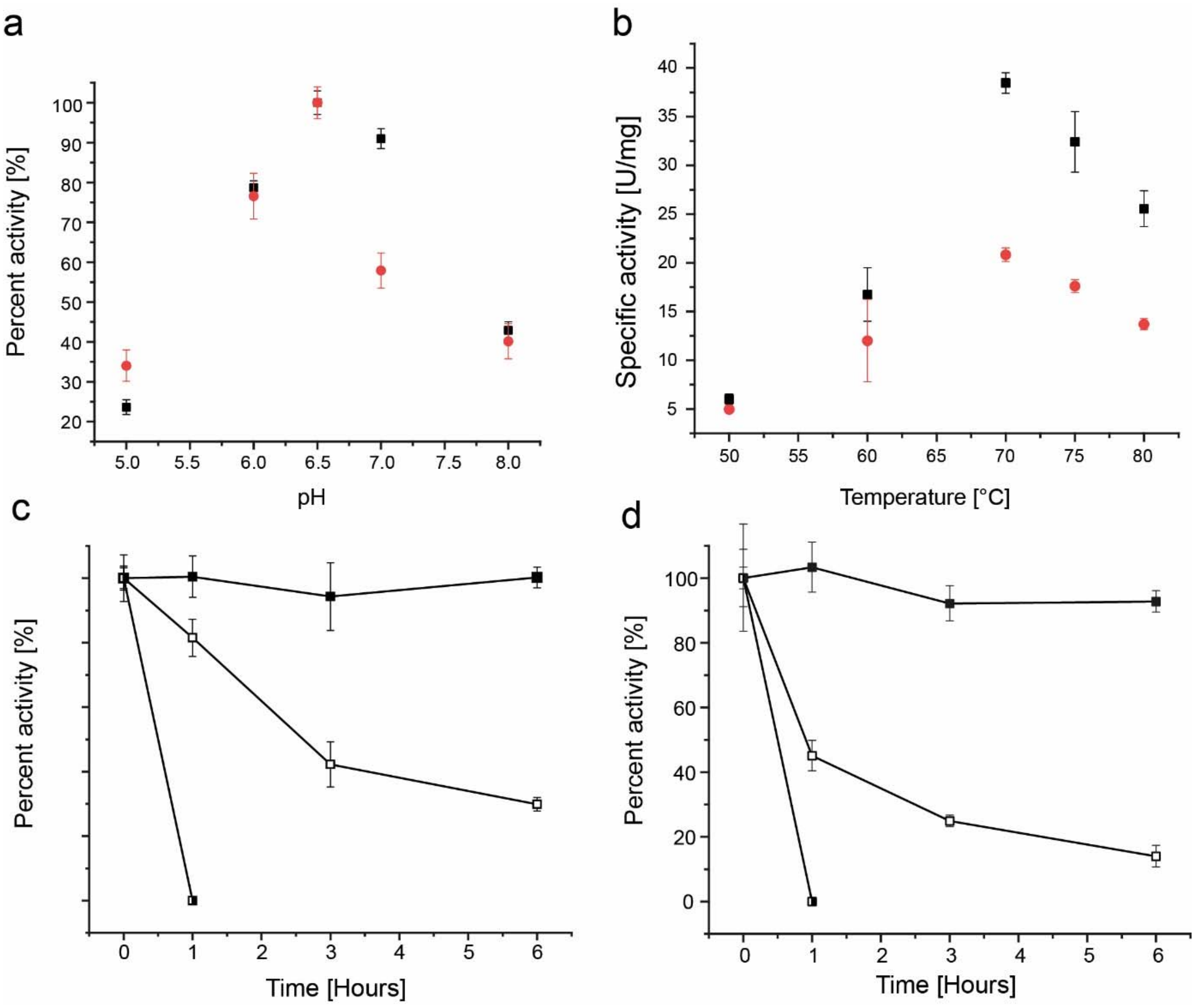


**Supplementary Fig. 6: Biochemical characterization of recombinant G3PDH Saci_2032.** (**a**) The pH dependence of Saci_1118 and Saci_2032 was determined in a continuous assay as G3P dependent reduction of DCPIP in a mixed buffer system (0.05 M MES, 0.05 M HEPES, 0.0 5M TRIS) at 70°C. (**b**) The optimum temperature of Saci _1118 and Saci_2032 was determined in the same assay system in 0.05 M MES-KOH pH 6.5 adjusted between 60°C to 80°C. **c**) Heat stability of Saci_2032 was determined by monitoring the residual activity upon incubation between 70°C to 90°C for up to 6 hours. The enzyme (protein concentration of 0.32 mg $ml^{-1}$) was incubated in 50 mM MES-KOH, pH 6.5 (temperature adjusted) and after the respective incubation times the residual G3PDH activity was determined with DCPIP as electron acceptor at 70°C as described above. (**d**) Heat stability of Saci_2032 was determined by monitoring the residual activity upon incubation between 70°C to 90°C for up to 6 hours. The enzyme (protein concentration of 0.18 mg $ml^{-1}$) was incubated in 50 mM MES-KOH, pH 6.5 (temperature adjusted) and after the respective incubation times the residual G3PDH activity was determined with DCPIP as electron acceptor at 70°C as described above. Experiments were performed in triplicate and error bars indicate the SD of the mean.

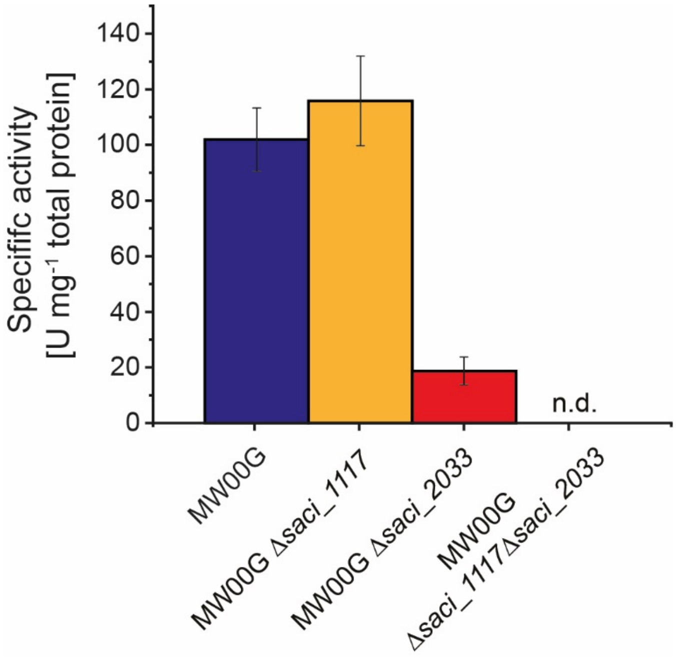


**Supplementary Fig. 7:** GK activity measured in crude extracts from parental MW00G and deletion mutants grown NZ-amine; n.d= not detectable.

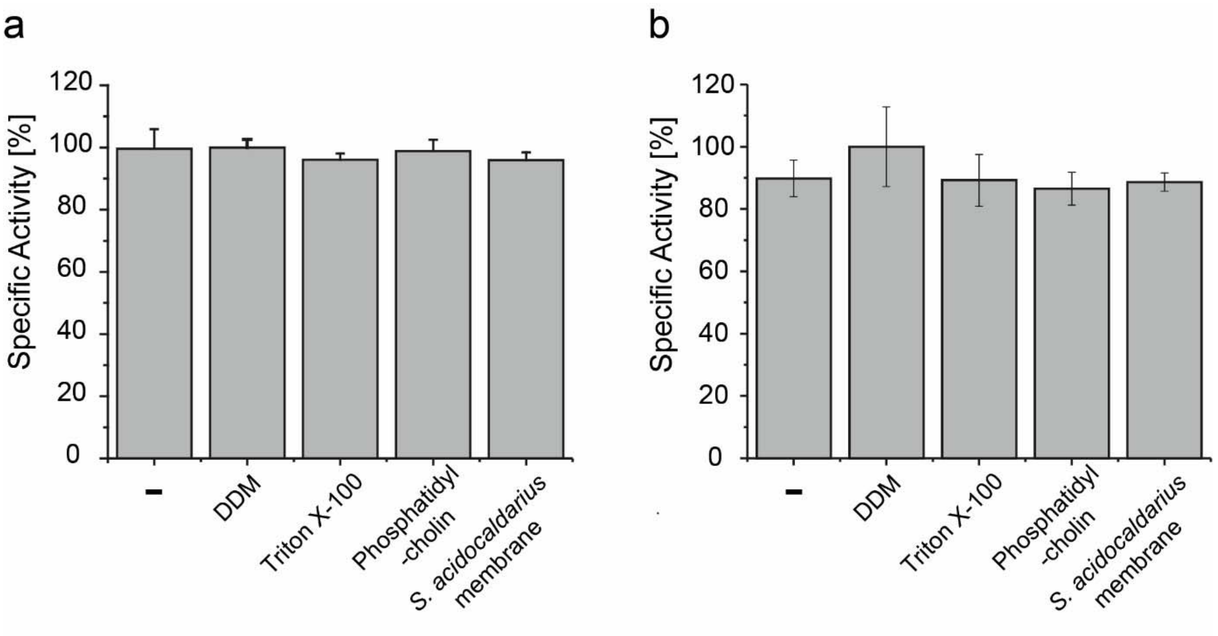


**Supplementary Fig. 8: Effect of non-ionic detergents and phospholipids on recombinant G3PDH Saci_2032 and Saci_1118.** Activity of Saci_2032 (**a**) and Saci_1118 (**b**) in the presence of potential activators was determined in a continuous assay by coupling the oxidation of G3P to the reduction of DCPIP in a MES-KOH buffer pH 6.5 at 50°C. 0.5 % DDM, 0.5 % of Triton X-100, 50 µg of phosphatidylcholine or 50 µg of isolated *S. acidocaldarius* membrane fractions were tested. Experiments were performed in triplicate and error bars indicate the SD of the mean.

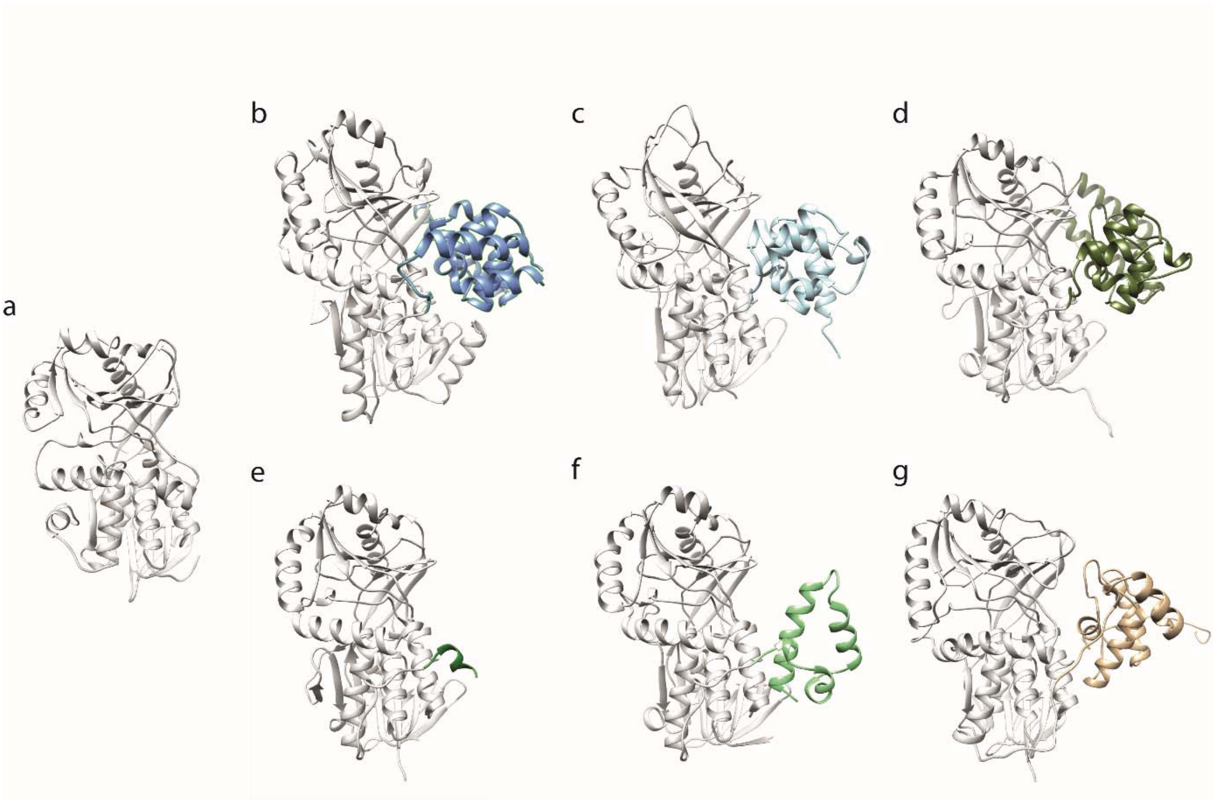


**Supplementary Fig. 9: Comparison of the overall fold of the catalytic subunits of the different types of FAD dependent G3P oxidizing enzymes.** The ribbon representations of crystal structures of (**a**) the *B. subtilis* glycine oxidase (1ng3) (Settembre et al., 2003) comprising the DAAO fold lacking any C-terminal extensions (shown in gray in all structures) is comparatively illustrated with those of (**b**) GlpO (2rgo) (Colussi et al., 2008), (**c**) GlpD (2qcu) (Yeh et al., 2008b), as well as the structural models (generated using CoLabFold (Mirdita et al., 2022)) of (**d**) *E. coli* GlpA, (**e**) the GlpA homologues of *T. pendens* and (**f**) *S. acidocaldarius*, and (g) the GlpTk from *T. kodakarensis* (TK1393). The C-terminal domains are highlighted using the same colour code as in Fig. 9-10 in the main text to emphasize the differences in size and fold.

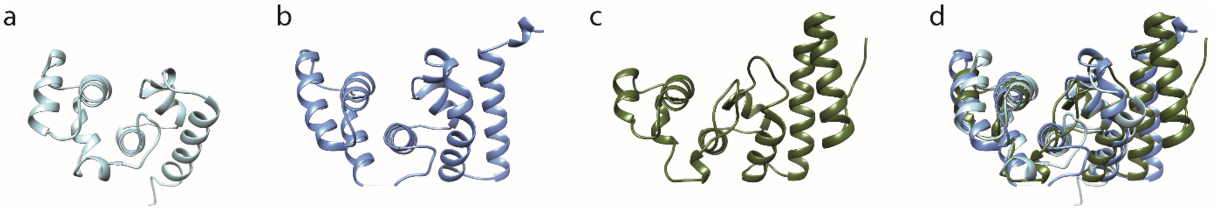


**Supplementary Fig. 10: Comparison of C-terminal domains of GlpD (a), GlpO (b), and GlpA (c).** The crystal structures of the C-termini of GlpD from *E. coli* (2qcu) (**a**) and GlpO from *Streptococcus* sp. (2rgo) (**b**) are shown, as well as the structural model of the C-terminus of GlpA from *E. coli* (**c**). The comparison reveals that the GlpD C-terminal fold is also present in GlpO and GlpA but extended by one and two helices, respectively. In (**d**) the superimposition of the three structures is shown in the same orientation as in (**a**)-(**c**). The dashed line in (**b**) indicates missing structural information for M518 in the C-terminus of *Streptococcus* sp. GlpO.

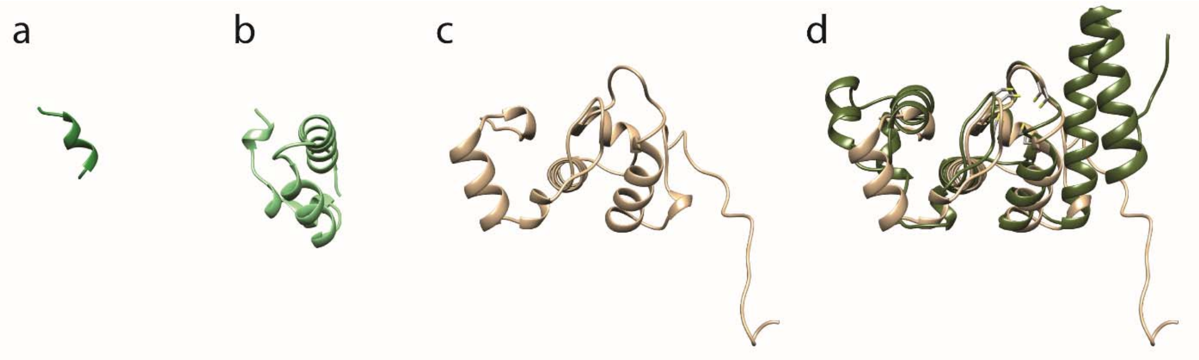


**Supplementary Fig. 11: Comparison of C-terminal extensions from GlpA-like G3PDHs from *T. pendens* (a) and *S. acidocaldarius* (Saci_2032, b), and of the GlpTk from *T. kodakarensis* (TK1393, c).** The ribbon representation of the structural models of the C-terminal are shown. To illustrate the presence of the bfd-fold in GlpTk and GlpA a superimposition of both C-termini is presented (**d**). Both structures match in their central parts as well as in the four conserved cysteine residues shown as stick models whereas the other parts of the structure differ remarkably.

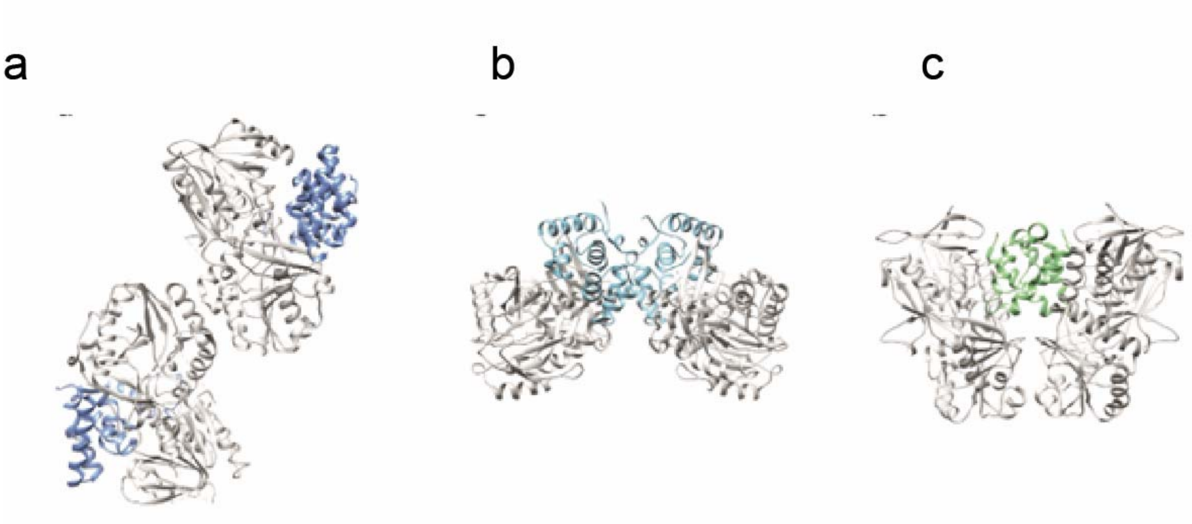


**Supplementary Fig. 12: Comparison of the dimer formation in the crystal structures of the (a) soluble, cytoplasmic GlpO (*Streptococcus*. spp., 2rgo) and (b) the monotopic, membrane bound GlpD (*E. coli*, 2qcu) with (c) the structural model of the GlpA-like Saci_2032 dimer.** The three structures illustrate the different role of the C-terminus (coloured according to the preceeding figures) in dimer formation as well as the resulting different spatial orientation of the DAAO-folds of the subunits to each other.
