## Supplementary File 1 for "Glycerol degradation in the thermoacidophilic crenarchaeon *Sulfolobus acidocaldarius* involves an unusual glycerol-3-phosphate dehydrogenase"

### Overview

| **ACE project** | **Title** |
| --- | --- |
| ACE_0602-01 | Identification of potential interaction partners of the CoxG homologs Saci_1119 and Saci_2031 in *Sulfolobus acidocaldarius* MW001 |
| ACE_0636-03 | Study of glycerol degradation in *S. acidocaldarius* MW00G |

### ACE_0602-01

#### File legend

| **ACE ID** | **Organism** | **Organ/ cell line** | **Treatment/ experimental setup** |
| --- | --- | --- | --- |
| ACE_0602-01_JB01 | *S. acidocaldarius* MW001 | Recomb. protein | Pulldown of overexpressed HA-tagged protein from *S.acidocaldarius* MW001 lysed in buffer B#_B incubated for 45 min. During pulldown, beads were washed using B#_A (2x) and MS standard water (3x)  JB01=neg. control empty vector (pSVAxylFX-CtHA), replicate 1 |
| ACE_0602-01_JB02 | *S. acidocaldarius* MW001 | Recomb. protein | Same setup as JB01, JB02 = neg. Control empty vector (pSVAxylFX-CtHA), replicate 2 |
| ACE_0602-01_JB03 | *S. acidocaldarius* MW001 | Recomb. protein | Same setup as JB01, JB03 = neg. control empty vector (pSVAxylFX-CtHA), replicate 3 |
| ACE_0602-01_JB04 | *S. acidocaldarius* MW001 | Recomb. protein | Same setup as JB01, JB04 = overexpressed Saci_2031 with C-terminal HA tag, (pSVA13204), replicate 1 |
| ACE_0602-01_JB05 | *S. acidocaldarius* MW001 | Recomb. protein | Same setup as JB01, JB05 = overexpressed Saci_2031 with C-terminal HA tag, (pSVA13204) replicate 2 |
| ACE_0602-01_JB06 | *S. acidocaldarius* MW001 | Recomb. protein | Same setup as JB01, JB06 = overexpressed Saci_2031 with C-terminal HA tag (pSVA13204), replicate 3 |
| ACE_0602-01_JB07 | *S. acidocaldarius* MW001 | Recomb. protein | Same setup as JB01, JB07 = overexpressed Saci_1119 with C-terminal HA tag (pSVA13202), replicate 1 |
| ACE_0602-01_JB08 | *S. acidocaldarius* MW001 | Recomb. protein | Same setup as JB01, JB08 = overexpressed Saci_1119 with C-terminal HA tag (pSVA13202), replicate 2 |
| ACE_0602-01_JB09 | *S. acidocaldarius* MW001 | Recomb. protein | Same setup as JB01, JB09 = overexpressed Saci_1119 with C-terminal HA tag (pSVA13202), replicate 3 |

#### LC_Settings

| MS device | Thermo Orbitrap Fusion Lumos |
| --- | --- |
| LC device | Thermo Easy-nLC 1200 |
| ion source | Thermo Nanospray Flex |
| **Analytical column** | Self-packed fused silica capillary, no frit; ESi Source Solutions PTC3-75-50-SP |
| column diameter | Length (L_C_) = 35 cm; ID = 75µm; OD = 360 µm; tip 5 µm |
| stationary phase | Reprosil-Pur 120 C18-AQ, Dr. Maisch GmbH |
| particle diameter (d_p_) | 1.9 µm |
| Pore size | 120 Å |
| Column ID | AC99 |
| Column oven | Sonation column oven PRSO-V2 |
| Column oven temp. | 50°C |
| **solvents** | A: 0.1% FA in UPLC water  B: 0.1% FA in 80% UPLC ACN |
| gradient | 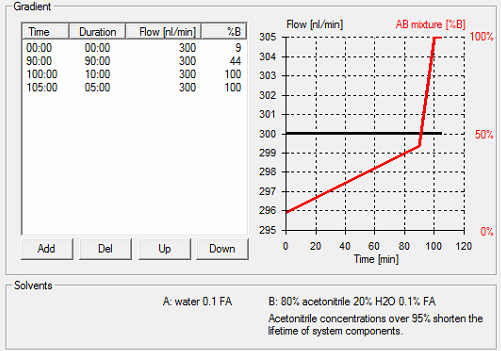 |

#### MS_Settings

| **Project** | **MS** | **general** | **MS1** | **MS2** | **MS2** | **MS3** | **Comments; special settings** |
| --- | --- | --- | --- | --- | --- | --- | --- |
| ACE_0602-01 | Lumos | Tune: v3.3.2782.28  Xcalibur: v4.3.73.11  Gradient: 105min | Analyzer: FT  Res.: 240000  SR: 375 - 1500  AGC: Standard  AcT: 50 ms  RF: 30  SF: --  DDM: CT/3sec | Analyzer: IT  Res./ScR: -/Rapid  SR: Auto  AGC: 300%  AcT: Auto  CS: +2 to +7  IsM: Q  IsW: 1.2  Frag.: sHCD  NCE: 20,30,45% |  |  | classic orbitrap experiment: MS1 in Orbitrap at high resolution and data dependent MS2 in Iontrap (IT) at rapid scan rate. Dynamic exclusion enabled (exclude after n times=1; Exclusion duration (s)= 30; mass tolerance= ± 10ppm)  Intensity Threshold: 5000  Ion transfer Tube Temp: 275 °C  Ion Source Voltage: 2500 V |

Note: **FT**= Fourier Transform (Orbitrap); **IT**= Iontrap; **Q**= Quadrupol; **Res.**= max. Resolution at 200 m/z (Lumos) or 400 m/z (Elite) [FWHM (full width at half maximum)]; **ScR**= scan rate for measurements in the IT; **SR**= scan range [m/z]; **AGC**= automatic gain control, max number of acquired ions per measurement; **AcT**= max. Ion acquisition time [ms]; **CS**= charge states used for fragmentation; **IsM**= Isolation mode (Q or IT), MS2 isolation and further is only done in IT; **IsW**= Isolation window [m/z], value followed by scan mode the isolation is based on (MS1, MS2 …) **Frag.**= Fragmentation method; **HCD**= Higher-energy collisional dissociation; **CID**= Collision-induced dissociation; **ETD**= Electron-transfer dissociation**; EThcD=** Electron-Transfer/Higher-Energy Collision Dissociation; **sHCD**= stepped HCD**; NCE**= normalized collision energy; **cycles**: number of MSn recorded or max cycle time; RF= RF Lens [%]; **SF**= Source Fragmentation [V]; **DDM**: Data dependent Mode (cycle time in seconds, CT/[s] or number of scans, NS); **NS**= Number of data dependent scans

### ACE_0636-03

#### File legend

| **ACE ID** | **Organism** | **Organ/ cell line** | **Treatment/ experimental setup** |
| --- | --- | --- | --- |
| ACE_0636-03_SN01 | *S. acidocaldarius* MW00G | MW00G | Growth on 0.2% NZA_biorep1 |
| ACE_0636-03_SN02 | S. acidocaldarius MW00G | MW00G | Growth on 0.2% NZA_biorep2 |
| ACE_0636-03_SN03 | S. acidocaldarius MW00G | MW00G | Growth on 0.2% NZA_biorep3 |
| ACE_0636-03_SN04 | S. acidocaldarius MW00G | MW00G | Growth on 0.2% NZA_biorep4 |
| ACE_0636-03_SN05 | S. acidocaldarius MW00G | MW00G | Growth on 10 mM glycerol_biorep1 |
| ACE_0636-03_SN06 | S. acidocaldarius MW00G | MW00G | Growth on 10 mM glycerol_biorep2 |
| ACE_0636-03_SN07 | S. acidocaldarius MW00G | MW00G | Growth on 10 mM glycerol_biorep3 |
| ACE_0636-03_SN08 | S. acidocaldarius MW00G | MW00G | Growth on 10 mM glycerol_biorep4 |
| ACE_0636-03_SN09 | S. acidocaldarius MW00G | MW00G | Growth on 0.2% D-xylose_biorep1 |
| ACE_0636-03_SN10 | S. acidocaldarius MW00G | MW00G | Growth on 0.2% D-xylose_biorep2 |
| ACE_0636-03_SN11 | S. acidocaldarius MW00G | MW00G | Growth on 0.2% D-xylose_biorep3 |
| ACE_0636-03_SN12 | S. acidocaldarius MW00G | MW00G | Growth on 0.2% D-xylose_biorep4 |
| ACE_0636-03_SN13 | S. acidocaldarius MW00G | MW00G | Growth on 20 mM glycerol_biorep1 |
| ACE_0636-03_SN14 | S. acidocaldarius MW00G | MW00G | Growth on 20 mM glycerol_biorep2 |
| ACE_0636-03_SN15 | S. acidocaldarius MW00G | MW00G | Growth on 20 mM glycerol_biorep3 |
| ACE_0636-03_SN16 | S. acidocaldarius MW00G | MW00G | Growth on 20 mM glycerol_biorep4 |
| ACE_0636-03_SN17 | S. acidocaldarius MW00G | MW00G | Growth on 40 mM glycerol_biorep1 |
| ACE_0636-03_SN18 | S. acidocaldarius MW00G | MW00G | Growth on 40 mM glycerol_biorep2 |
| ACE_0636-03_SN19 | S. acidocaldarius MW00G | MW00G | Growth on 40 mM glycerol_biorep3 |
| ACE_0636-03_SN20 | S. acidocaldarius MW00G | MW00G | Growth on 40 mM glycerol_biorep4 |

#### LC_Settings

| MS device | Thermo Orbitrap Fusion Lumos |
| --- | --- |
| LC device | Thermo Easy-nLC 1200 |
| ion source | Thermo Nanospray Flex |
| **Analytical column** | Self-packed fused silica capillary with an integrated sintered frit; CoAnn Technologies ICT36007515F-50-5 |
| column diameter | Length (L_C_) = 50 cm; ID = 75µm; OD = 360 µm; emitter tip 15 µm |
| stationary phase | Reprosil-Pur 120 C18-AQ, Dr. Maisch GmbH |
| particle diameter (d_p_) | 1.9 µm |
| Pore size | 120 Å |
| Column ID | AC103 |
| Column oven | Sonation column oven PRSO-V2 |
| Column oven temp. | 50°C |
| **solvents** | A: 0.1% FA in UPLC water  B: 0.1% FA in 80% UPLC ACN |
| gradient | 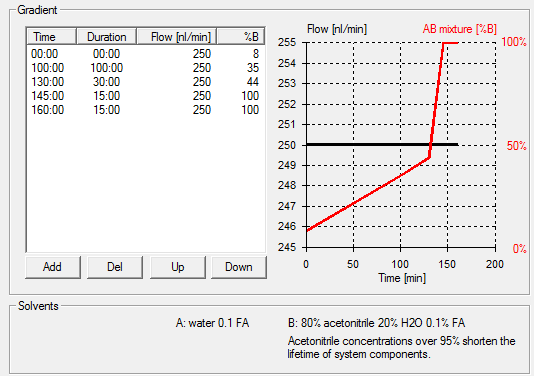 |

#### MS_Settings

| **Project** | **MS** | **general** | **MS1** | **MS2** | **MS2** | **MS3** | **Comments; special settings** |
| --- | --- | --- | --- | --- | --- | --- | --- |
| ACE_0636-03 | Lumos | Tune: v3.3.2782.28  Xcalibur: v4.3.73.11  Gradient: 160 min | Analyzer: FT  Res.: 120000  SR: 375 - 1750  AGC: Standard  AcT: 50 ms  RF: 30  SF: --  DDM: CT/3sec | Analyzer: IT  Res./ScR: -/Rapid  SR: Auto  AGC: 300%  AcT: Auto  CS: +2 to +7  IsM: Q  IsW: 1.2  Frag.: stepped HCD  NCE: 20,30,40% |  |  | classic orbitrap experiment: MS1 in Orbitrap at high resolution and data dependent MS2 in Iontrap (IT) at rapid scan rate. Dynamic exclusion enabled (exclude after n times=1; Exclusion duration (s)= 30; mass tolerance= ± 10ppm)  Intensity Threshold: 5000  Ion transfer Tube Temp: 275 °C  Ion Source Voltage: 2500 V |

Note: **FT**= Fourier Transform (Orbitrap); **IT**= Iontrap; **Q**= Quadrupol; **Res.**= max. Resolution at 200 m/z (Lumos) or 400 m/z (Elite) [FWHM (full width at half maximum)]; **ScR**= scan rate for measurements in the IT; **SR**= scan range [m/z]; **AGC**= automatic gain control, max number of acquired ions per measurement; **AcT**= max. Ion acquisition time [ms]; **CS**= charge states used for fragmentation; **IsM**= Isolation mode (Q or IT), MS2 isolation and further is only done in IT; **IsW**= Isolation window [m/z], value followed by scan mode the isolation is based on (MS1, MS2 …) **Frag.**= Fragmentation method; **HCD**= Higher-energy collisional dissociation; **CID**= Collision-induced dissociation; **ETD**= Electron-transfer dissociation**; EThcD=** Electron-Transfer/Higher-Energy Collision Dissociation; **sHCD**= stepped HCD**; NCE**= normalized collision energy; **cycles**: number of MSn recorded or max cycle time; RF= RF Lens [%]; **SF**= Source Fragmentation [V]; **DDM**: Data dependent Mode (cycle time in seconds, CT/[s] or number of scans, NS); **NS**= Number of data dependent scans
